# Robust group- but limited individual-level (longitudinal) reliability and insights into cross-phases response prediction of conditioned fear

**DOI:** 10.1101/2022.03.15.484434

**Authors:** Maren Klingelhöfer-Jens, Mana R. Ehlers, Manuel Kuhn, Vincent Keyaniyan, Tina B. Lonsdorf

## Abstract

Here we follow the call to target measurement reliability as a key prerequisite for individual-level predictions in translational neuroscience by investigating i) longitudinal reliability at the individual and ii) group level, iii) cross-sectional reliability and iv) response predictability across experimental phases. 120 individuals performed a fear conditioning paradigm twice six month apart. Analyses of skin conductance responses, fear ratings and BOLD-fMRI with different data transformations and included numbers of trials were conducted. While longitudinal reliability was generally poor to moderate at the individual level, it was good for acquisition but not extinction at the group-level. Cross-sectional reliability was satisfactory. Higher responding in preceding phases predicted higher responding in subsequent experimental phases at a weak to moderate level depending on data specifications. In sum, the results suggest the feasibility of individual-level predictions for (very) short time intervals (e.g., cross-phases) while predictions for longer time intervals may be problematic.

---

The crucial role of adequate statistical power in neuroscience (Button et al., 2013) has gained momentum in the past years and resulted in calls for and efforts to increase sample sizes (Moriarity & Alloy, 2021; Szucs & Ioannidis, 2020). Measurement reliability, however, has only recently gained attention as a crucial determinant of the maximally observable effect size (Fröhner, Teckentrup, Smolka, & Kroemer, 2019; Hedge, Powell, & Sumner, 2018; Zuo, Xu, & Milham, 2019). This is particularly important for individual-level predictions and clinical questions such as “Why do some individuals develop pathological anxiety while others do not?” or “Why do some patients benefit from treatment while others relapse?”

In the laboratory, such fear and treatment related processes can be studied using the fear conditioning paradigm, which is considered a key paradigm “offering the best current opportunity for translating neuroscience discoveries into clinical applications” (cf. Anderson & Insel, 2006, p. 319). To date, both clinical and experimental research has primarily focused on group-level, basic, generic principles (Lonsdorf & Merz, 2017). Tackling clinical questions regarding individual prediction of symptom development or treatment outcome, however, requires a shift towards and a validation of research methods tailored to individual differences - such as a focus on measurement reliability (Zuo, Xu, & Milham, 2019).

In differential fear conditioning protocols (see Lonsdorf et al., 2017), one stimulus is repetitively paired with an aversive unconditioned stimulus (US; e.g., electro-tactile stimulation), and as a consequence becomes a conditioned stimulus (CS+) while another stimulus, the CS-, is never paired with the US. After this acquisition training phase, CSs are presented without the US (extinction training) leading to a gradual waning of the conditioned response. Critically, the fear memory (CS+/US association) is not erased, but a competing inhibitory extinction memory (CS+/noUS association) is assumed to be formed during extinction training (Milad & Quirk, 2012; Myers & Davis, 2007). Subsequently, return of fear (RoF) can be induced by procedural manipulations such as a time delay (spontaneous recovery), a contextual change (renewal) or a (re-)presentation of an aversive event (reinstatement) and conditioned responding can be subsequently probed in a RoF test phase. During the RoF test, the absence (i.e., extinction retention) or the return of conditioned responding (i.e., return of fear) can be observed (Bouton, 2004; Lonsdorf et al., 2017).

Experimental research employing fear conditioning paradigms has a strong translational value in aiming to aid the development of optimized intervention and prevention programs with extinction learning assumed to be the active component of exposure therapy (Graham & Milad, 2011; Milad & Quirk, 2012; Rachman, 1989; Vervliet, Craske, & Hermans, 2013) and experimental RoF suggested as a model of clinical relapse (Scharfenort, Menz, & Lonsdorf, 2016; Vervliet, Craske, & Hermans, 2013). However, successful clinical translation - particularly treatment outcome predictions - requires that both the experimental paradigm and the measures employed allow for individual-level predictions over and above prediction of group-averages (Fröhner, Teckentrup, Smolka, & Kroemer, 2019; Hedge, Powell, & Sumner, 2018; Lonsdorf & Merz, 2017). A prerequisite for this is that the measures show stability within and reliable differences between individuals over time. The degree of temporal stability over repetitions can be assessed by longitudinal reliability (also termed test-retest reliability). Importantly, longitudinal reliability (for details see Table 1) also has implications for the precision with which associations of one variable (e.g., conditioned responding) with another (individual difference) variable can be measured because the correlation between those two variables cannot exceed the correlations within any of these two variables [i.e., the longitudinal reliability; Spearman (1910)].

**Table 1:**
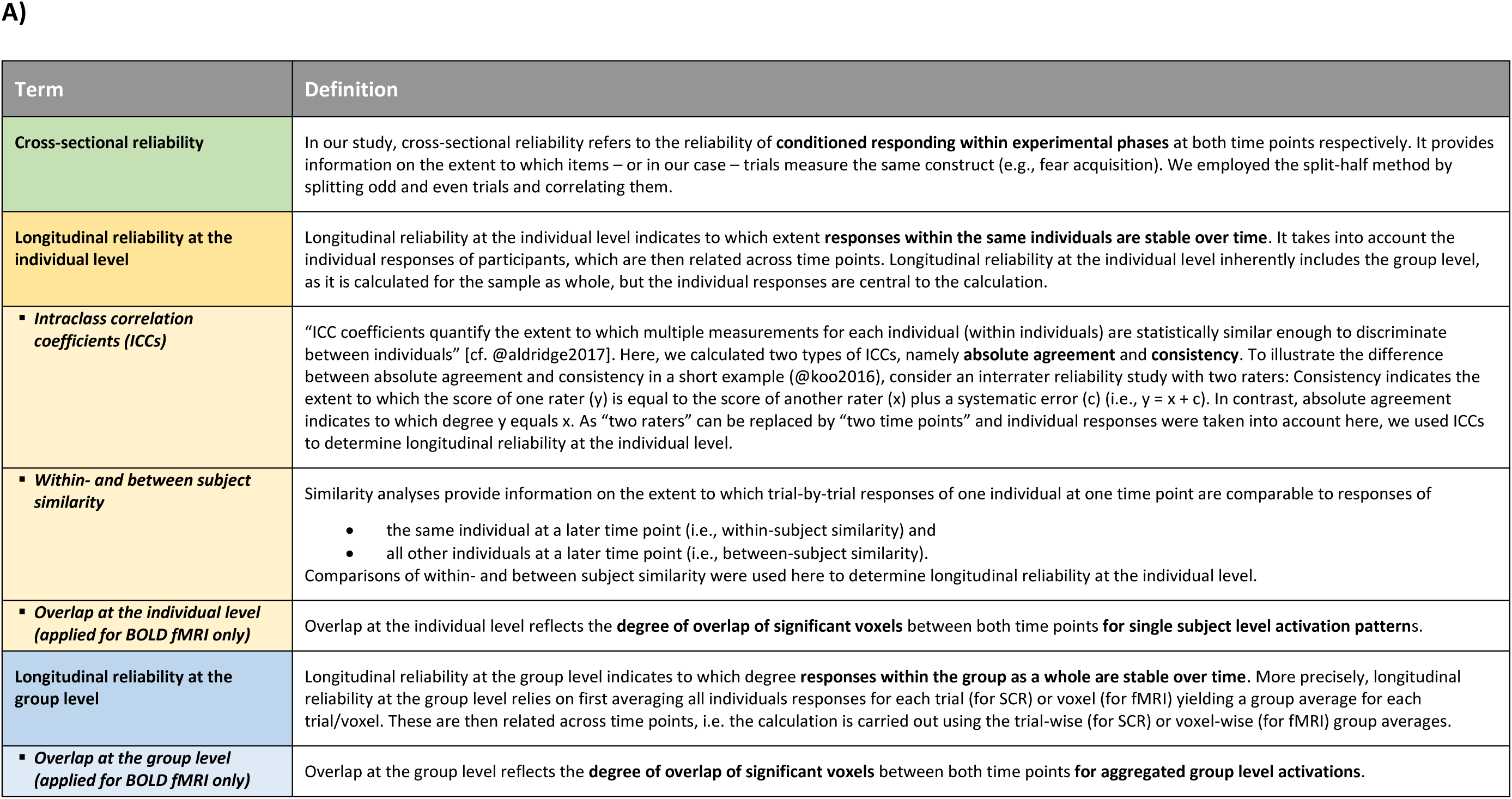
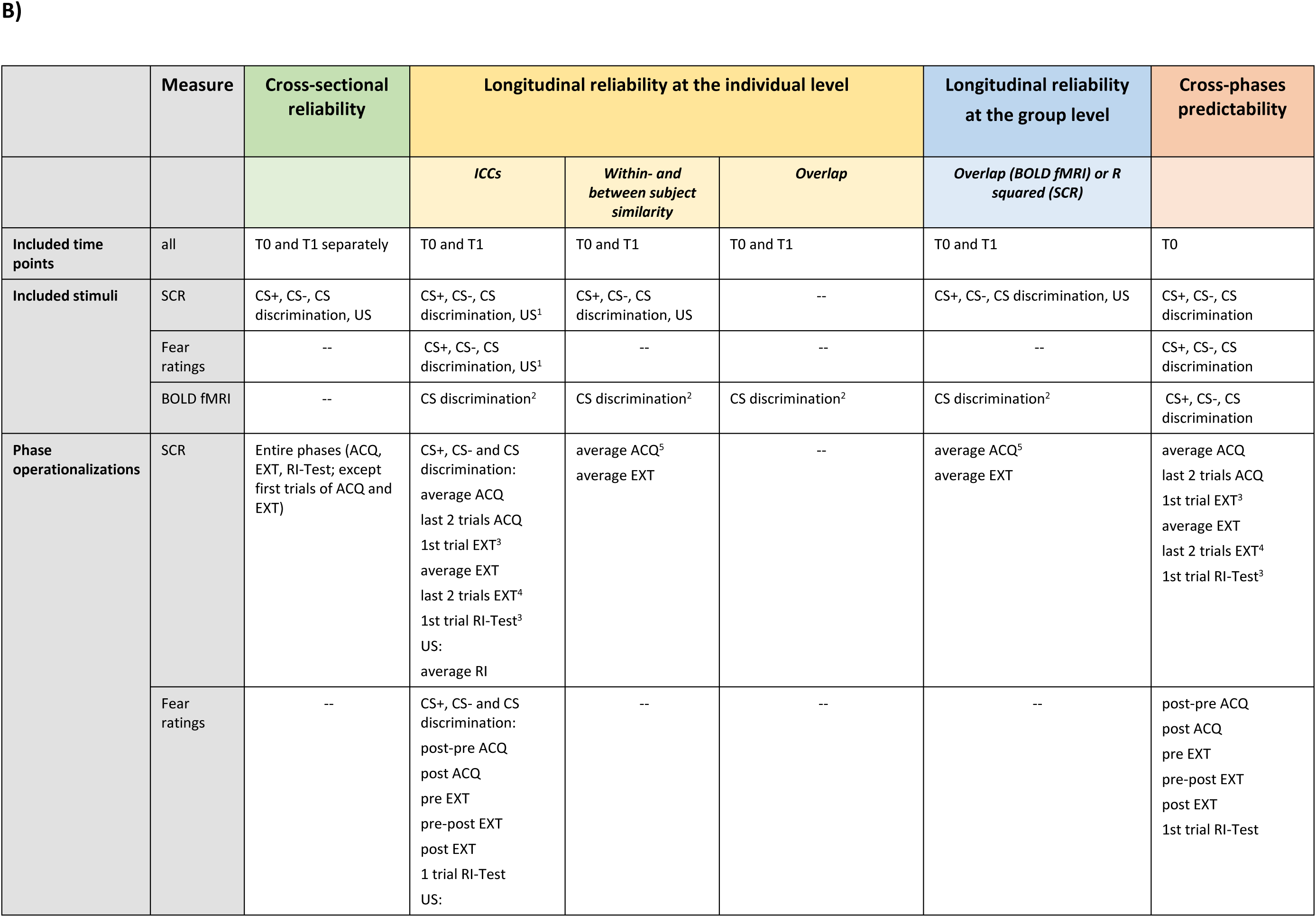
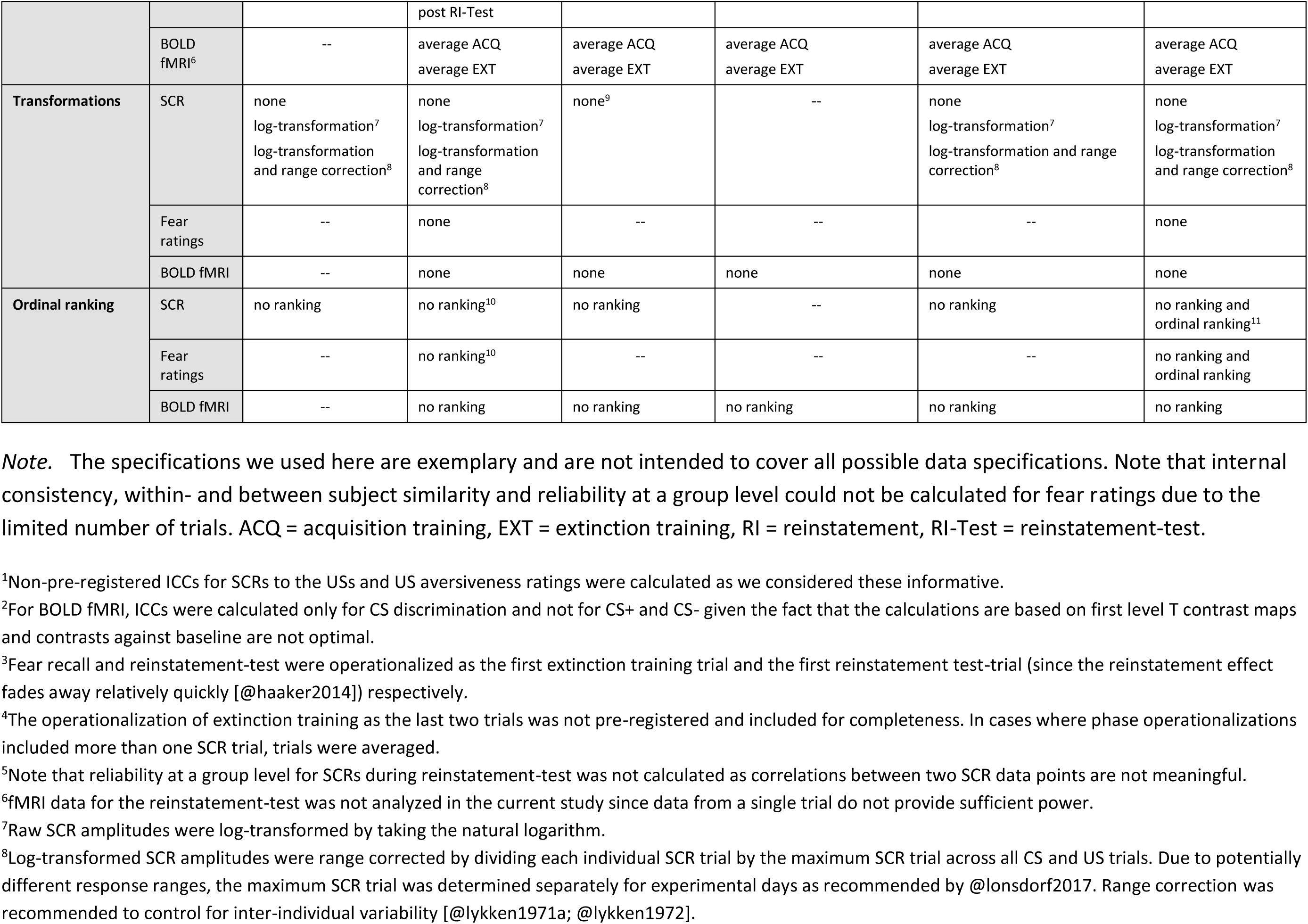
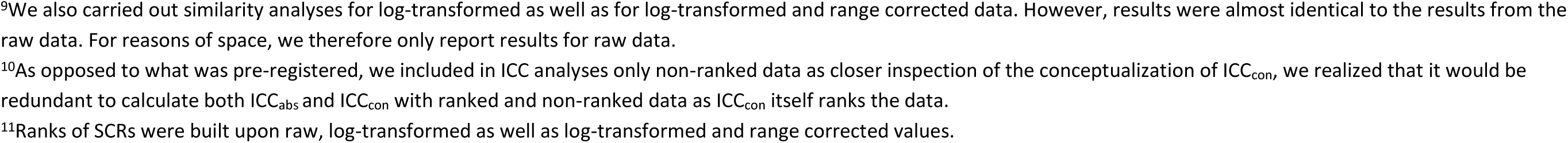
Definitions of key terms (A) and data specifications applied across analyses (B).

| Term | Definition |
| --- | --- |
| **Cross-sectional reliability** | In our study, cross-sectional reliability refers to the reliability of **conditioned responding within experimental phases** at both time points respectively. It provides information on the extent to which items – or in our case – trials measure the same construct (e.g., fear acquisition). We employed the split-half method by splitting odd and even trials and correlating them. |
| **Longitudinal reliability at the individual level** | Longitudinal reliability at the individual level indicates to which extent **responses within the same individuals are stable over time**. It takes into account the individual responses of participants, which are then related across time points. Longitudinal reliability at the individual level inherently includes the group level, as it is calculated for the sample as whole, but the individual responses are central to the calculation. |
| ▪ **Intraclass correlation coefficients (ICCs)** | “ICC coefficients quantify the extent to which multiple measurements for each individual (within individuals) are statistically similar enough to discriminate between individuals” [cf. @aldridge2017]. Here, we calculated two types of ICCs, namely **absolute agreement** and **consistency**. To illustrate the difference between absolute agreement and consistency in a short example (@koo2016), consider an interrater reliability study with two raters: Consistency indicates the extent to which the score of one rater (y) is equal to the score of another rater (x) plus a systematic error (c) (i.e., $y = x + c$). In contrast, absolute agreement indicates to which degree y equals x. As “two raters” can be replaced by “two time points” and individual responses were taken into account here, we used ICCs to determine longitudinal reliability at the individual level. |
| ▪ **Within- and between subject similarity** | Similarity analyses provide information on the extent to which trial-by-trial responses of one individual at one time point are comparable to responses of<br>• the same individual at a later time point (i.e., within-subject similarity) and<br>• all other individuals at a later time point (i.e., between-subject similarity).<br>Comparisons of within- and between subject similarity were used here to determine longitudinal reliability at the individual level. |
| ▪ **Overlap at the individual level (applied for BOLD fMRI only)** | Overlap at the individual level reflects the **degree of overlap of significant voxels** between both time points **for single subject level activation patterns**. |
| **Longitudinal reliability at the group level** | Longitudinal reliability at the group level indicates to which degree **responses within the group as a whole are stable over time**. More precisely, longitudinal reliability at the group level relies on first averaging all individuals responses for each trial (for SCR) or voxel (for fMRI) yielding a group average for each trial/voxel. These are then related across time points, i.e. the calculation is carried out using the trial-wise (for SCR) or voxel-wise (for fMRI) group averages. |
| ▪ **Overlap at the group level (applied for BOLD fMRI only)** | Overlap at the group level reflects the **degree of overlap of significant voxels** between both time points **for aggregated group level activations**. |

B)
|  | Measure | Cross-sectional reliability | Longitudinal reliability at the individual level |  |  | Longitudinal reliability at the group level | Cross-phases predictability |
| --- | --- | --- | --- | --- | --- | --- | --- |
|  |  |  | ICCs | Within- and between subject similarity | Overlap | Overlap (BOLD fMRI) or R squared (SCR) |  |
| Included time points | all | T0 and T1 separately | T0 and T1 | T0 and T1 | T0 and T1 | T0 and T1 | T0 |
| Included stimuli | SCR | CS+, CS-, CS discrimination, US | CS+, CS-, CS discrimination, US<sup>1</sup> | CS+, CS-, CS discrimination, US | -- | CS+, CS-, CS discrimination, US | CS+, CS-, CS discrimination |
|  | Fear ratings | -- | CS+, CS-, CS discrimination, US<sup>1</sup> | -- | -- | -- | CS+, CS-, CS discrimination |
|  | BOLD fMRI | -- | CS discrimination<sup>2</sup> | CS discrimination<sup>2</sup> | CS discrimination<sup>2</sup> | CS discrimination<sup>2</sup> | CS+, CS-, CS discrimination |
| Phase operationalizations | SCR | Entire phases (ACQ, EXT, RI-Test; except first trials of ACQ and EXT) | CS+, CS- and CS discrimination:<br>average ACQ<br>last 2 trials ACQ<br>1st trial EXT<sup>3</sup><br>average EXT<br>last 2 trials EXT<sup>4</sup><br>1st trial RI-Test<sup>3</sup><br>US:<br>average RI | average ACQ<sup>5</sup><br>average EXT | -- | average ACQ<sup>5</sup><br>average EXT | average ACQ<br>last 2 trials ACQ<br>1st trial EXT<sup>3</sup><br>average EXT<br>last 2 trials EXT<sup>4</sup><br>1st trial RI-Test<sup>3</sup> |
|  | Fear ratings | -- | CS+, CS- and CS discrimination:<br>post-pre ACQ<br>post ACQ<br>pre EXT<br>pre-post EXT<br>post EXT<br>1 trial RI-Test<br>US: | -- | -- | -- | post-pre ACQ<br>post ACQ<br>pre EXT<br>pre-post EXT<br>post EXT<br>1st trial RI-Test |

|  |  |  | post RI-Test |  |  |  |  |
| --- | --- | --- | --- | --- | --- | --- | --- |
|  | BOLD fMRI<sup>6</sup> | -- | average ACQ<br>average EXT | average ACQ<br>average EXT | average ACQ<br>average EXT | average ACQ<br>average EXT | average ACQ<br>average EXT |
| **Transformations** | SCR | none<br>log-transformation<sup>7</sup><br>log-transformation<br>and range correction<sup>8</sup> | none<br>log-transformation<sup>7</sup><br>log-transformation<br>and range<br>correction<sup>8</sup> | none<sup>9</sup> | -- | none<br>log-transformation<sup>7</sup><br>log-transformation and range<br>correction<sup>8</sup> | none<br>log-transformation<sup>7</sup><br>log-transformation<br>and range correction<sup>8</sup> |
|  | Fear ratings | -- | none | -- | -- | -- | none |
|  | BOLD fMRI | -- | none | none | none | none | none |
| **Ordinal ranking** | SCR | no ranking | no ranking<sup>10</sup> | no ranking | -- | no ranking | no ranking and<br>ordinal ranking<sup>11</sup> |
|  | Fear ratings | -- | no ranking<sup>10</sup> | -- | -- | -- | no ranking and<br>ordinal ranking |
|  | BOLD fMRI | -- | no ranking | no ranking | no ranking | no ranking | no ranking |

*Note.* The specifications we used here are exemplary and are not intended to cover all possible data specifications. Note that internal consistency, within- and between subject similarity and reliability at a group level could not be calculated for fear ratings due to the limited number of trials. ACQ = acquisition training, EXT = extinction training, RI = reinstatement, RI-Test = reinstatement-test.
<sup>1</sup>Non-pre-registered ICCs for SCRs to the USs and US aversiveness ratings were calculated as we considered these informative.
<sup>2</sup>For BOLD fMRI, ICCs were calculated only for CS discrimination and not for CS+ and CS- given the fact that the calculations are based on first level T contrast maps and contrasts against baseline are not optimal.
<sup>3</sup>Fear recall and reinstatement-test were operationalized as the first extinction training trial and the first reinstatement test-trial (since the reinstatement effect fades away relatively quickly [aaker2014]) respectively.
<sup>4</sup>The operationalization of extinction training as the last two trials was not pre-registered and included for completeness. In cases where phase operationalizations included more than one SCR trial, trials were averaged.
<sup>5</sup>Note that reliability at a group level for SCRs during reinstatement-test was not calculated as correlations between two SCR data points are not meaningful.
<sup>6</sup>fMRI data for the reinstatement-test was not analyzed in the current study since data from a single trial do not provide sufficient power.
<sup>7</sup>Raw SCR amplitudes were log-transformed by taking the natural logarithm.
<sup>8</sup>Log-transformed SCR amplitudes were range corrected by dividing each individual SCR trial by the maximum SCR trial across all CS and US trials. Due to potentially different response ranges, the maximum SCR trial was determined separately for experimental days as recommended by lonsdorf2017. Range correction was recommended to control for inter-individual variability [lykken1971a; lykken1972].
<sup>9</sup>We also carried out similarity analyses for log-transformed as well as for log-transformed and range corrected data. However, results were almost identical to the results from the raw data. For reasons of space, we therefore only report results for raw data.
<sup>10</sup>As opposed to what was pre-registered, we included in ICC analyses only non-ranked data as closer inspection of the conceptualization of $ICC_{con}$ , we realized that it would be redundant to calculate both $ICC_{abs}$ and $ICC_{con}$ with ranked and non-ranked data as $ICC_{con}$ itself ranks the data.
<sup>11</sup>Ranks of SCRs were built upon raw, log-transformed as well as log-transformed and range corrected values.

In fear conditioning, surprisingly little is known about **longitudinal reliability at the individual level** - which reflects the temporal stability of (averaged) **individual responding** (typically assessed through e.g., intra-class correlation coefficients, ICCs, see Table 1) - with time intervals in prior work ranging from three weeks to eight months (Fredrikson, Annas, Georgiades, Hursti, & Tersman, 1993; Ridderbusch et al., 2021; Torrents-Rodas et al., 2014; Zeidan et al., 2012).

Generally (details in Supplementary Table 1), individual-level longitudinal reliability of risk ratings, skin conductance responses (SCRs) and fear potentiated startle (FPS) was within the same range (Torrents-Rodas et al., 2014) whereas it was numerically somewhat lower for the BOLD response as compared to different rating types (Ridderbusch et al., 2021). Longitudinal reliability at the individual level appeared higher for acquisition training than for extinction training (SCRs: Fredrikson, Annas, Georgiades, Hursti, & Tersman, 1993; Zeidan et al., 2012) and higher for extinction training than for reinstatement-test (for BOLD but not ratings: Ridderbusch et al., 2021) and higher for CS+ than CS- responses (SCRs: Fredrikson, Annas, Georgiades, Hursti, & Tersman, 1993) and CS discrimination (ratings and BOLD fMRI: Ridderbusch et al., 2021; SCRs: Zeidan et al., 2012).

However, it is difficult to extract a comprehensive picture from these four studies as they differ substantially in sample size (*N* = 18 - 100), paradigm specifications, experimental phases reported, outcome measures, time intervals and employed reliability measures (see Supplementary Table 1).

Given that the majority of research in fear conditioning targets group-level generic mechanisms and does not necessarily focus on individual response patterns, it is even more surprising that, to our knowledge, no study to date has investigated longitudinal reliability at the group level and only few studies targeted (Fredrikson, Annas, Georgiades, Hursti, & Tersman, 1993) cross-sectional reliability (i.e., split-half reliability, see Table 1). In contrast to the typically investigated longitudinal reliability at the individual level, **longitudinal reliability at the group level** indicates the extent to which responses **averaged across the group** as a whole are stable over time. Even though it has to be acknowledged that the group average is not necessarily representative of any individual in the group and the same group average may arise from different and even opposite individual responses at both time points in the same group, group level reliability is important to establish in addition to individual-level reliability as much work focuses on group-level inference.

In addition to the importance of establishing longitudinal reliability (i.e., reliability of measurements between different time points) as a key prerequisite for individual level predictions across longer time intervals, it is often implicitly assumed that responding in one experimental phase predicts responding in a subsequent phase. As a result it has been suggested to routinely ‘correct’ for responding during fear acquisition training when studying performance in later experimental phases such as extinction training or retention/RoF test (critically discussed in Lonsdorf et al., 2019). However, empirical evidence on this cross-phases predictability (see Table 1) is scarce to date.

Evidence from experimental work on cross-phase predictability in rodents and humans is mixed. In rodents, freezing during acquisition training and 24h-delayed extinction training were uncorrelated (Plendl & Wotjak, 2010) and responding during extinction training did not predict extinction retention (i.e., lever-pressing suppression: Bouton, García-Gutiérrez, Zilski, & Moody, 2006; or freezing behavior: Shumake, Furgeson-Moreira, & Monfils, 2014). Similarly, in humans, extinction performance (FPS, SCRs and US expectancy ratings) did not predict performance at 24h-retention test (Prenoveau, Craske, Liao, & Ornitz, 2013). Yet, a computational modeling approach suggests that the quality of extinction learning predicts spontaneous recovery in SCRs (Gershman & Hartley, 2015).

Also evidence from work in patient samples is mixed (for a review, see Craske et al., 2008). The extent of fear reduction **within** therapeutic sessions was unrelated to overall treatment outcome in some studies (Kozak, Foa, & Steketee, 1988; Pitman et al., 1996; Riley et al., 1995), while others observed an association (Foa et al., 1983). Similarly, significant correlations of fear reduction **between** therapeutic sessions with treatment outcome were observed for reported distress (Rauch, Foa, Furr, & Filip, 2004) and heart rate, but not for SCR (Kozak, Foa, & Steketee, 1988; Lang, Melamed, & Hart, 1970) and for self-reported fear post treatment, but not at follow-up (Foa et al., 1983). In addition, evidence that responding in different phases is related comes from pharmacological manipulations with the cognitive enhancer D-cycloserine which facilitates learning and/or consolidation. D-cyloserine promoted long term extinction retention (Rothbaum et al., 2014; Smits, Rosenfield, Otto, Marques, et al., 2013; Smits, Rosenfield, Otto, Powers, et al., 2013) only if within-session learning was achieved.

With this pre-registered study, we follow the call for more appreciation and systematic investigation of measurement reliability (Zuo, Xu, & Milham, 2019). We address longitudinal and cross-sectional reliability as well as predictability of cross-phase responding in SCRs, fear ratings and the BOLD response. For this purpose, we re-analyzed data from 120 participants that underwent a differential fear conditioning paradigm twice (at time points T0 and T1, six months apart) - with habituation and acquisition training on day 1 and extinction, reinstatement and reinstatement-test on day 2 to allow for fear memory consolidation prior to extinction.

Specifically, we i) assessed cross-sectional reliability of SCRs at both time points and ii) systematically assessed longitudinal reliability of SCRs, fear ratings and BOLD fMRI at the individual level by calculating ICCs. Based on previous results, we expect poor to moderate longitudinal reliability at the individual level for all outcome measures. This is complemented by investigations of response similarity (SCR and BOLD fMRI) and the degree of overlap of activated voxels at both time points (BOLD fMRI) as additional measurements of longitudinal reliability at the individual level (see Table 1). As findings from fear conditioning studies are mostly based on group results, we also iii) assessed whether SCR and BOLD fMRI show longitudinal reliability at the group level. Finally, we iv) expect that individual-level responding during an experimental phase is significantly predictive of individual-level responding during subsequent experimental phases. All hypotheses are tested across different data specifications for CS type, phase operationalization and transformations to account for procedural heterogeneity in the literature (see Supplementary Table 1).

## Results

For a comprehensive overview of the different reliability measures used here and of the analyses conducted, see Table 1.

### Satisfactory cross-sectional reliability

To assess cross-sectional reliability of SCRs, trials were split into odd and even trials (i.e., odd-even approach), averaged for each individual subject and then correlated (Pearson correlation coefficient). This was done separately for each time point and experimental phase. Cross-sectional reliability at T0 (see Figure 1A) and T1 (see Figure 1B) of raw SCRs to the CS+ and CS- were acceptable to good during both acquisition and extinction training, but poor to questionable for reinstatement-test. Cross-sectional reliability for CS discrimination, however, was generally unacceptable to poor, but excellent for the US. Log-transformation did not impact cross-sectional reliability but log-transformation and range resulted mostly in reduced reliability (see Supplementary Figure 1).

**Figure 1.**
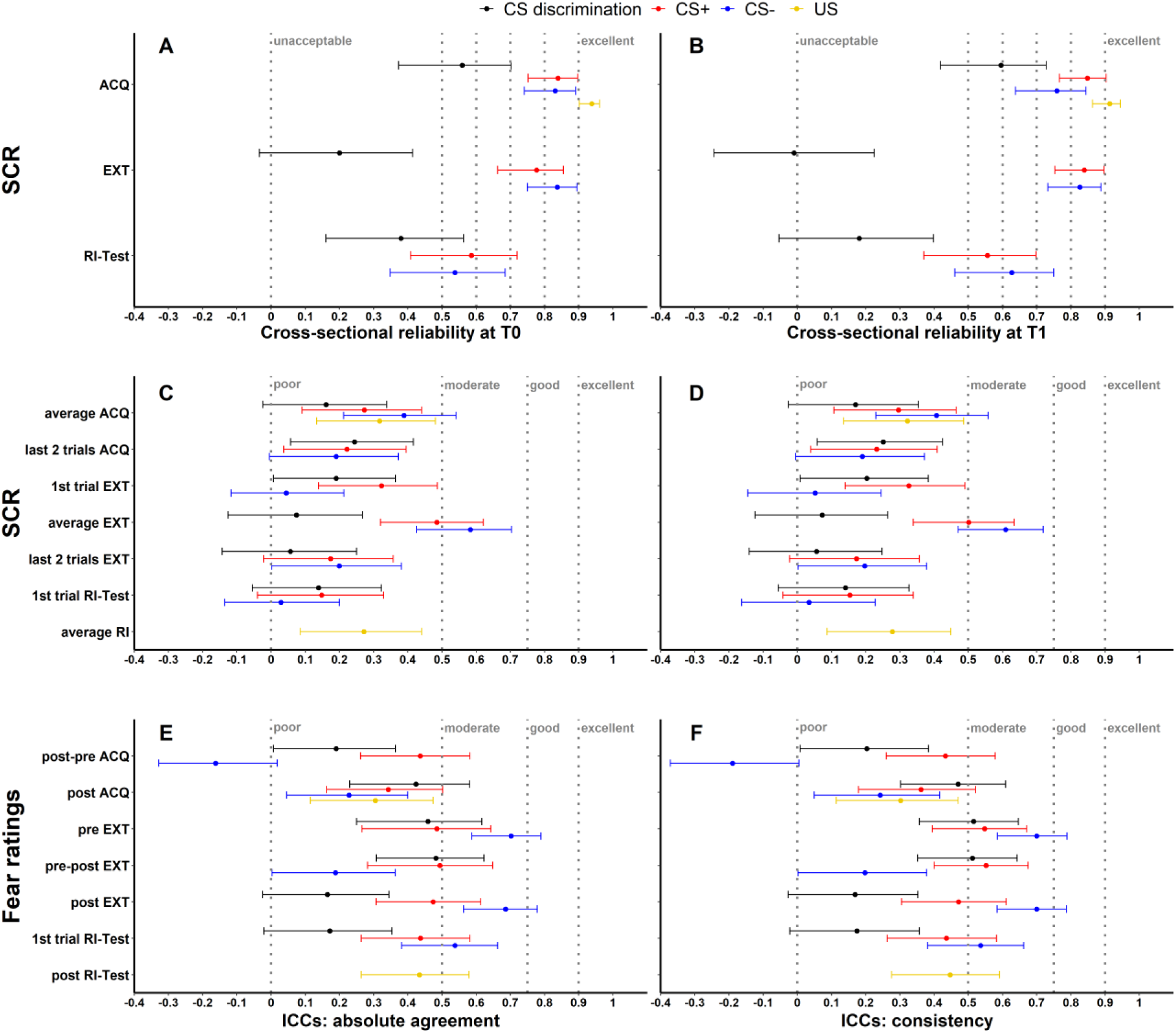
Illustration of cross-sectional reliability for SCRs at T0 (A) and T1 (B) as well as ICC_abs_ and ICC_con_ for SCRs (C,D) and fear ratings (E,F) color coded for stimulus-type. Note that assessment of cross-sectional reliability was not possible for fear ratings as only two ratings (pre, post) were available. Error bars represent 95% confidence intervals indicating significance, when zero is not included in the interval. The y-axis comprises the different phases or phase operationalizations. Cross-sectional reliability is interpreted using benchmarks (Kline, 2013) for unacceptable (< 0.5), poor (> 0.5 but < 0.6), questionable (> 0.6 but < 0.7), acceptable (> 0.7 but < 0.8), good (> 0.8 but < 0.9) and excellent (≥ 0.9). ICCs were interpreted as poor (< 0.5), moderate (> 0.5 but < 0.75), good (> 0.75 but < 0.9) and excellent (≥ 0.9) (Koo & Li, 2016). Note that cross-sectional reliability and ICCs for SCRs are shown for raw data only. Results of log-transformed as well as log-transformed and range corrected data are presented in Supplementary Figures 7 and 8 for completeness. ACQ = acquisition training, EXT = extinction training, RI = reinstatement, RI-Test = reinstatement-test, pre = prior to the experimental phase, post = subsequent to the experimental phase.

### Longitudinal reliability at the individual level

Longitudinal reliability at the individual level refers to the time-stability of individual responses which is assessed here through several measures (see Table 1).

#### Low intra-class correlation coefficients for SCRS, fear ratings and BOLD response

As a first measure, absolute agreement ICCs (ICC_abs_) and consistency ICCs (ICC_con_) were calculated across both time-points (T0, T1) for all data specifications (see Figure 1) while for BOLD fMRI these were only calculated for CS discrimination (see methods for justification).

#### SCR and fear ratings

Across most data specifications, ICC_abs_ and ICC_con_ indicated generally poor longitudinal reliability at the individual level (SCR: range_ICCabs_ = -0.086 - 0.616, range_ICCcon_ = -0.103 - 0.657; fear ratings: range_ICCabs_ = -0.162 - 0.702, range_ICCcon_ = -0.19 - 0.7, for detailed results see also Supplementary Table 3 and 4) for both SCRs and fear ratings with few exceptions of moderate reliability (see Figure 1). ICCs for log-transformed and raw SCRs were similar (see Supplementary Figure 2C-F) while log-transformation and range correction resulted in increased reliability for some data specifications (e.g., CS+ and CS- responses averaged across acquisition training) but in reduced reliability for others (e.g., CS- responses during fear recall, i.e., the first extinction trial).

Exploratory, non-preregistered analyses of trial-by-trial SCRs generally revealed only minor changes in ICCs when more SCR trials were included stepwise into the calculations (see Supplementary Figures 3 to 8) with few exceptions: Including more trials numerically shifted ICC point estimates for SCRs to the CS+ and CS- during acquisition (log-transformed and range corrected data) and extinction training (all transformation-types) from a poor to a moderate range. Note, however, that this was - at large - only statistically significant when comparing ICCs based on the first (i.e., single trial at T0 and T1) and the maximum number of trials (as indicated by non-overlapping 95% CI error bars). Interestingly, ICC point estimates for reinstatement-test (all transformation-types) were numerically lower with an increasing number of trials, likely because of the transitory nature of the reinstatement effect.

#### BOLD fMRI

For BOLD fMRI, both ICC-types suggest poor reliability for CS discrimination during acquisition (both ICC_abs_ and ICC_con_ = 0.18) and extinction training (both ICC_abs_ and ICC_con_ = 0.01). For individual ROIs (insula, amygdala, hippocampus, dmPFC and vmPFC), ICCs were even lower (all ICCs ≤ 0.001; for full results see Supplementary Table 5). In sum, individual-level responses in the same paradigm six months apart showed - at large - poor to moderate temporal stability in all three outcome measures.

#### Higher within- than between-subject similarity in BOLD fMRI but not SCRs

While ICCs provide information on the absolute quantity of longitudinal reliability at the individual level, comparison of within-subject and between-subject similarity as a complementary measure of longitudinal reliability at the individual level (see Table 1) reflects the extent to which responses in SCR and BOLD activation of one individual at T0 were more similar to themselves at T1 than to other individuals at T1.

#### SCR

For SCRs, within-subject similarity (i.e., within-subject correlation of trial-by-trial SCR across time points) and between-subject similarity (i.e., correlation of trial-by-trial SCR between one individual at T0 and all other individuals at T1; see Figure 2) did not differ significantly for most data specifications. This was true for CS discrimination (*t*(64) = 1.78, *p* = .079, *d* = 0.22) as well as SCRs to the CS+ (*t*(61) = 0.84, *p* = .407, *d* = 0.11) and CS- (*t*(55) = 1.50, *p* = .138, *d* = 0.20) during acquisition training and for CS discrimination (*t*(44) = -0.23, *p* = .823, *d* = -0.03) and SCRs to the CS+ (*t*(39) = 0.25, *p* = .801, *d* = 0.04) during extinction training. This indicates that SCRs of one particular individual at T0 were mostly not more similar to their own SCRs than to those of other individuals at T1. The only exceptions where within-subject similarities were significantly higher than between subject similarity were SCRs to the US during acquisition training (*t*(70) = 2.54, *p* = .013, *d* = 0.30) and to the CS- during extinction training (*t*(31) = 2.05, *p* = .049, *d* = 0.36). Note, however, that within-subject similarity had a very wide spread pointing to substantial individual differences (while this variance is removed in calculations of between-subject similarity).

**Figure 2.**
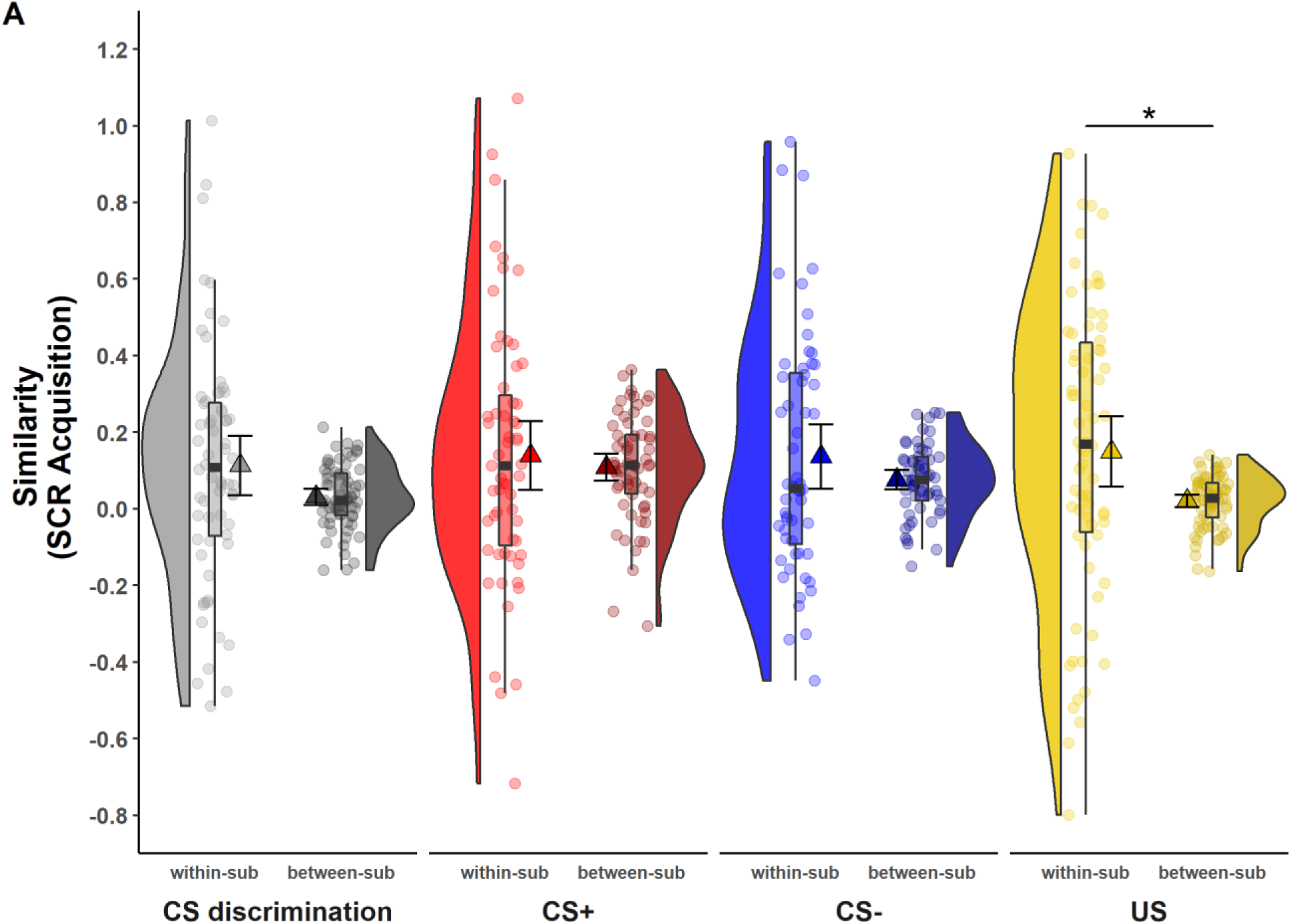
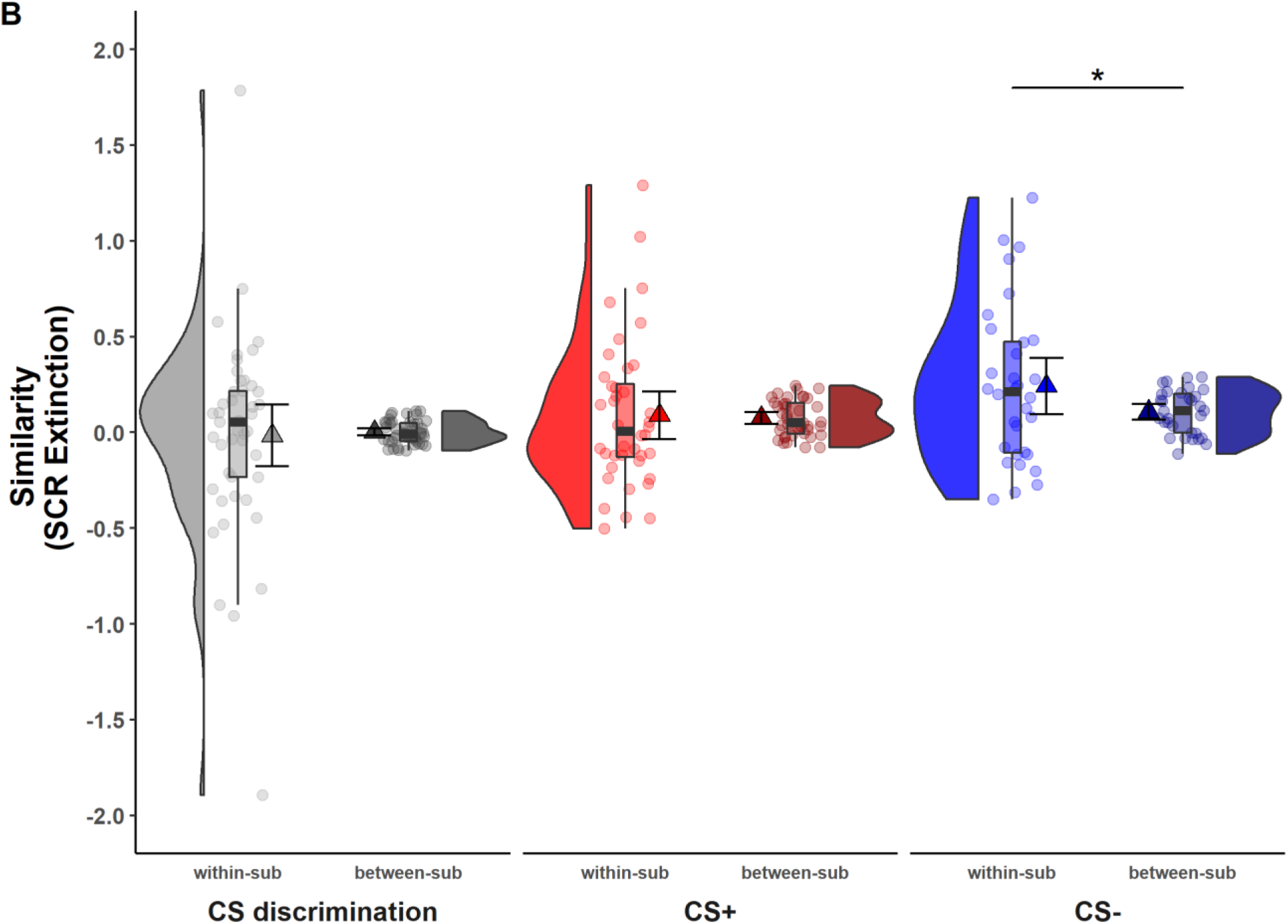
Illustration of within-subject and between-subject similarity for raw SCRs during (A) acquisition and (B) extinction training separately for CS discrimination (grey), CS+ (red), CS- (blue) and US responses (yellow). Results for log-transformed as well as log-transformed and range corrected SCRs were almost identical to the results from raw data and are hence not reported here. Single data points represent Fisher r-to-z transformed correlations between single trial SCRs of each subject at T0 and T1 (within-subject similarity) or averaged r-to-z transformed correlations between single trial SCRs of one subject at T0 and all other subjects at T1 (between-subject similarity). Triangles represent mean correlations, corresponding error bars represent 95% confidence intervals. Boxes of boxplots represent the interquartile range (IQR) crossed by the median as bold line, ends of whiskers represent the minimum/maximum value in the data within the range of 25th/75th Percentiles 1.5∗IQR. Distributions of the data are illustrated with densities next to the boxplots. One data point was above 3.5 (within-subject similarity of SCRs to the CS+) and is not shown in the figure. ∗*p* < .05. Note that the variances differ strongly between within- and between-subject similarity because between-subject similarity is based on correlations averaged across subjects, whereas within-subject similarity is based on non-averaged correlations calculated for each subject. Note also that similarity calculations were based on different sample sizes for acquisition and extinction training and CS discrimination as well as SCRs to the CS+, CS- and US respectively (for details, see methods). within-sub = within-subject; between-sub = between-subject.

#### fMRI data

In contrast to what was observed for SCRs, within-subject similarity was significantly higher than between-subject similarity in the whole brain (*p* < .001) and in all ROIs for fear acquisition training (all *p*’s < .048; see Figure 3A and Supplementary Table 6). This suggests that while absolute values for similarity might be low, individual brain activation patterns during fear acquisition training at T0 are still more similar to the same subject’s activation pattern at T1 than to any others at T1. For extinction training, however, no significant differences between within- and between-subject similarity were found for any ROI or the whole brain (all *p*’s > .221; see Figure 3B and Supplementary Table 6).

**Figure 3.**
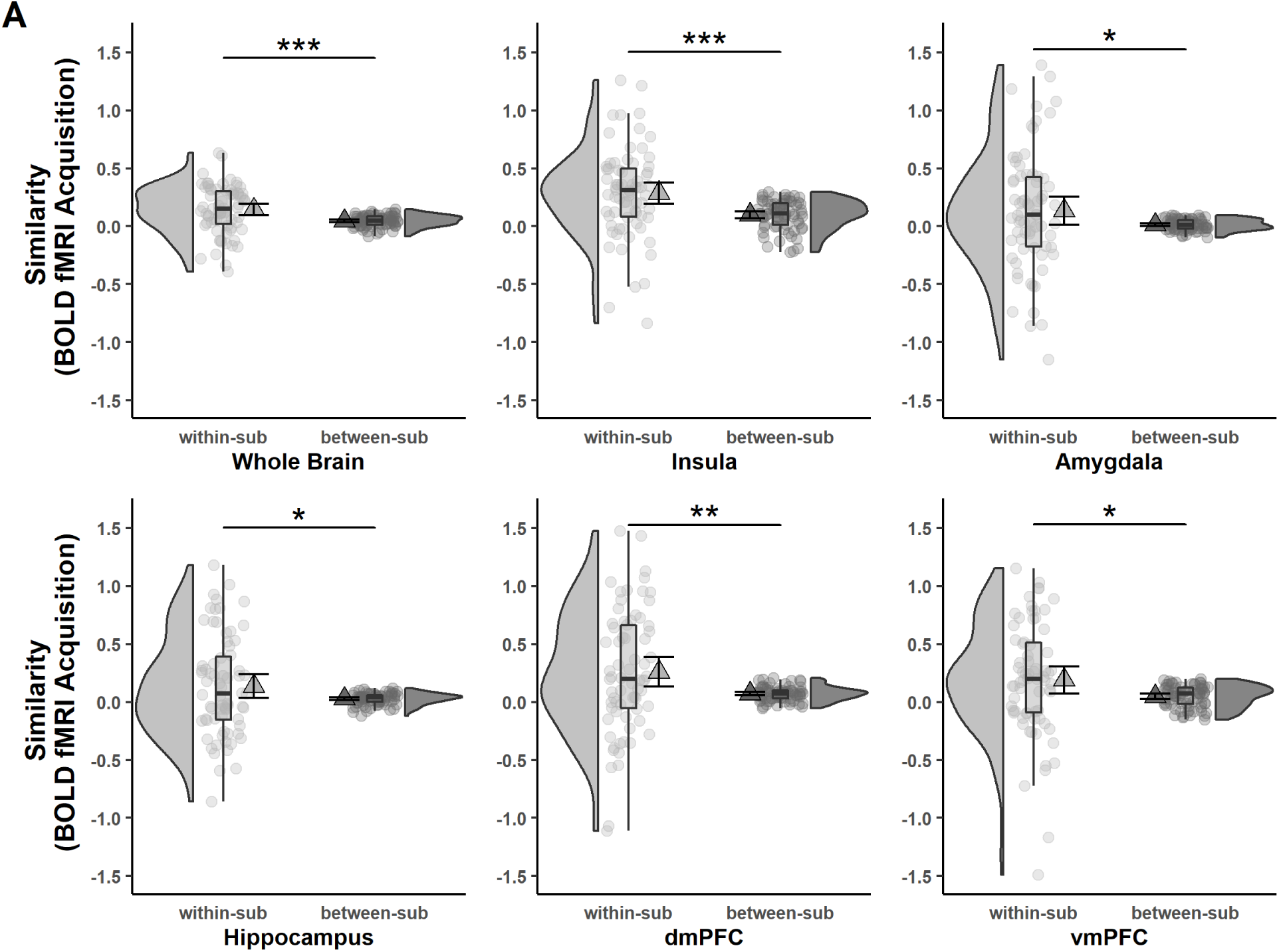
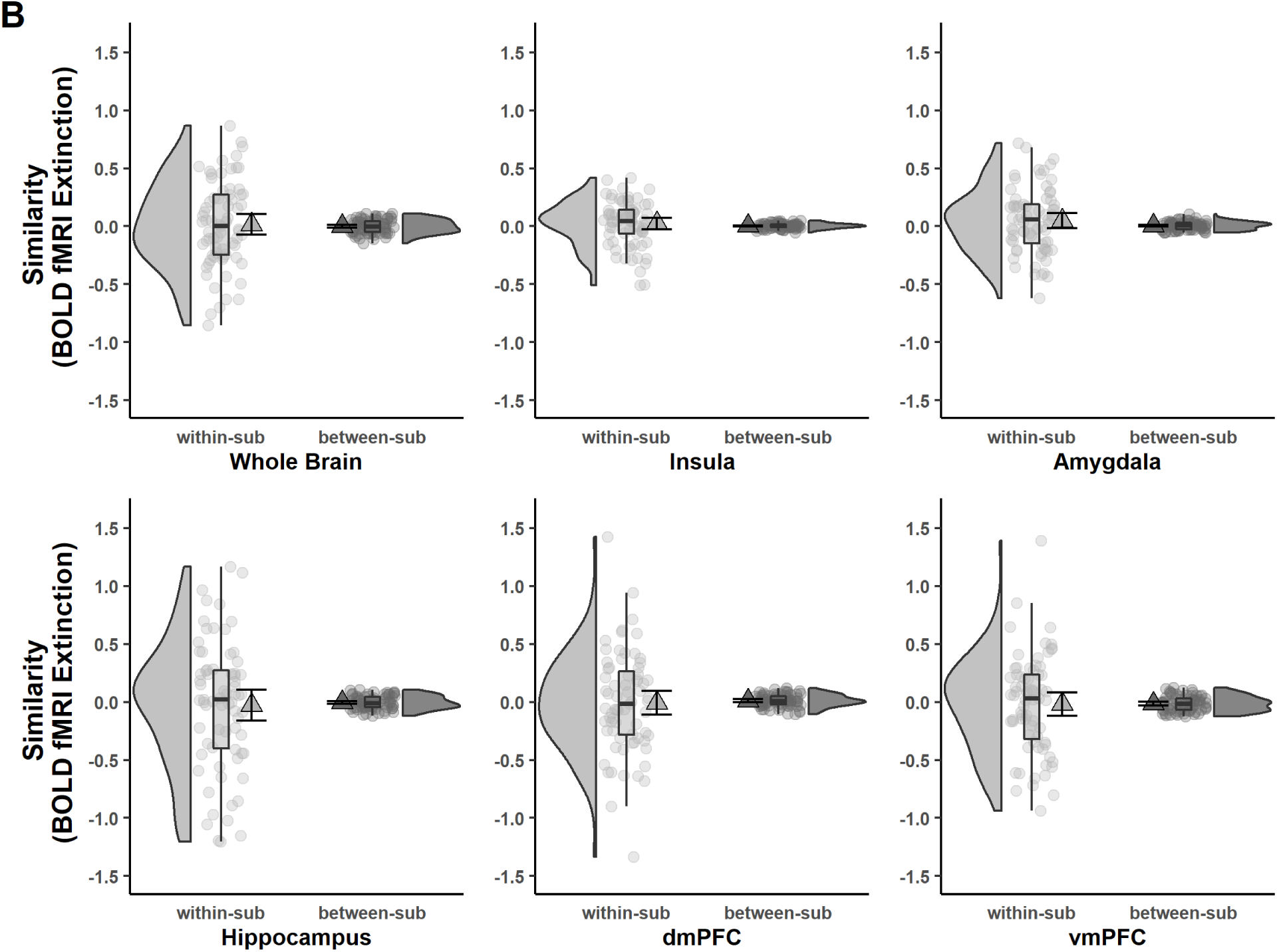
Acquisition (A) and extinction (B) within-subject and between-subject similarities (Fisher r-to-z transformed) of voxel-wise brain activation patterns (based on beta maps) for CS discrimination at T0 and T1 for the whole brain and different ROIs. Triangles represent mean correlations, corresponding error bars represent 95% confidence intervals. Single data points represent Fisher r-to-z transformed correlations between the first-level response patterns of brain activation of each subject at T0 and T1 (within-subject similarity) or averaged r-to-z transformed correlations between the first-level response patterns of brain activation of one subject at T0 and all other subjects at T1 (between-subject similarity). Boxes of boxplots represent the interquartile range (IQR) crossed by the median as bold line, ends of whiskers represent the minimum/maximum value in the data within the range of 25th/75th Percentiles 1.5∗IQR. Distributions of the data are illustrated with densities next to the boxplots. fMRI data for the reinstatement-test were not analyzed in the current study since data from a single trial do not provide sufficient power. ∗*p* < .05, ∗∗*p* < .01, ∗∗∗*p* < .001. dmPFC = dorsomedial prefrontal cortex; vmPFC = ventromedial prefrontal cortex; within-sub = within-subject; between-sub = between-subject.

#### Low overlap at the individual level between both time points

As opposed to similarity measures (see above) which reflect the correlation of activated voxels between time points, overlap at the individual level denotes the degree of overlap of significantly activated voxels.

The overlap at the individual level was low with the Jaccard coefficient indicating 7.60 % and 0.70 % whole brain overlap for acquisition and extinction training respectively (see Table 2A). Of note, individual values ranged from 0 - 39.65% overlap during acquisition, suggesting large interindividual differences in overlap.

**Table 2:** Overlap in significantly activated voxels at the individual and group level across both time points for CS discrimination.

| Level | Phase | Coeff. | ROI |  |  |  |  |  |
| --- | --- | --- | --- | --- | --- | --- | --- | --- |
|  |  |  | Whole Brain | Insula | Amygdala | Hippocampus | dmPFC | vmPFC |
| **(A) Individual** | Acq | Jaccard | 0.076 | 0.075 | 0.011 | 0.012 | 0.043 | 0.039 |
|  |  | Dice | 0.131 | 0.121 | 0.018 | 0.021 | 0.070 | 0.061 |
|  | Ext | Jaccard | 0.007 | 0.001 | 0.000 | 0.000 | 0.005 | 0.005 |
|  |  | Dice | 0.014 | 0.001 | 0.000 | 0.000 | 0.010 | 0.009 |
| **(B) Group** | Acq | Jaccard | 0.620 | 0.595 | 0.294 | 0.323 | 0.526 | 0.045 |
|  |  | Dice | 0.765 | 0.745 | 0.448 | 0.472 | 0.689 | 0.086 |
|  | Ext | Jaccard | 0.046 | 0.000 | 0.000 | 0.000 | 0.044 | 0.000 |
|  |  | Dice | 0.089 | 0.000 | 0.000 | 0.000 | 0.085 | 0.000 |

*Note.* Results are shown for the whole brain as well as for selected ROIs for fear acquisition training and extinction training. Both coefficients range from 0 (no overlap) to 1 (perfect overlap). Note that the Jaccard can be interpreted as % (Maitra, 2010). dmPFC = dorsomedial prefrontal cortex; vmPFC = ventromedial prefrontal cortex.

While overlap during acquisition for individual ROIs was comparable to the whole brain, Jaccard and Dice coefficients indicate close to 0 overlap at extinction (see Table 2A).

### Robust longitudinal reliability at the group level

While longitudinal reliability at the individual level relies on (mean) individual subject responding at both time points, longitudinal reliability at the group level relies on the percentage of explained variance of group averaged trials at T1 by group averaged trials at T0 (i.e., R squared for SCR) or the degree of group level overlap of significant voxels expressed as Dice and Jaccard indices (i.e., BOLD fMRI).

#### SCR

For acquisition training (see Figure 4A), 40.66% (*F*(1,11) = 7.54, *p* = .019), 63.59% (*F*(1,11) = 19.21, *p* = .001) and 75.67% (*F*(1,11) = 34.20, *p* < .001) of the variance of SCRs at T1 could be explained by SCRs at T0 for CS discrimination, CS+ and CS- respectively indicating robust longitudinal reliability of SCRs at the group level for CS responding during acquisition. Interestingly, only 19.53% (*F*(1,12) = 2.91, *p* = .114) of the variance of SCRs to the US could be explained. For extinction training, in contrast, only 19.58% (*F*(1,11) = 2.68, *p* = .130) and 21.70% (*F*(1,11) = 3.05, *p* = .109) of the SCR variance at T1 could be explained by SCRs at T0 for CS discrimination and CS+ respectively indicating only limited longitudinal reliability at the group level. However, with 67.35% (*F*(1,11) = 22.69, *p* = .001) explained variance at T1, longitudinal reliability of SCRs to the CS- seemed to be more robust as compared to CS discrimination and responses to the CS+ (see Figure 4B).

**Figure 4.**
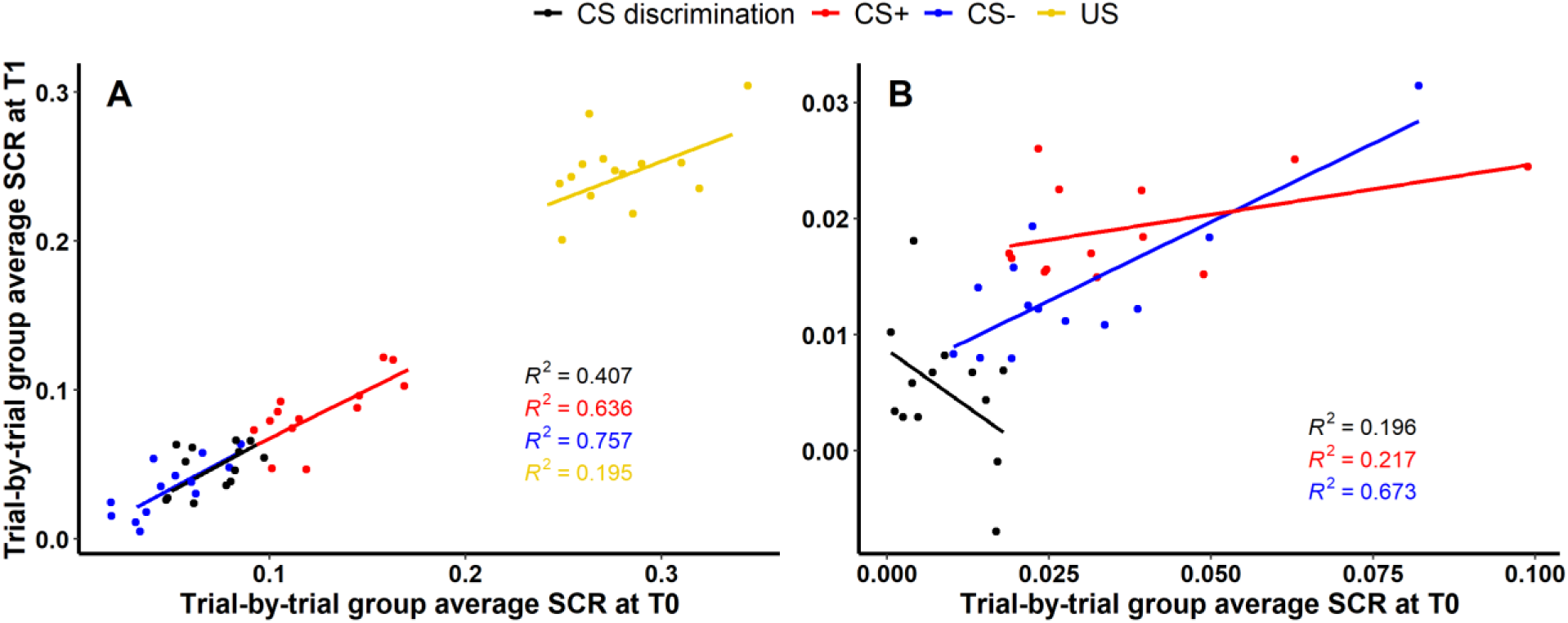
Scatter plots illustrating longitudinal reliability at the group level during (A) acquisition and (B) extinction training for raw SCRs (in μS). Results for log-transformed as well as log-transformed and range corrected data are presented in the Supplement (Supplementary Figure 9). Longitudinal reliability at the group level refers to the extent of explained variance in linear regressions comprising SCRs at T0 as independent and SCRs at T1 as dependent variable. Results are shown for trial-by-trial group average SCRs to the CS+ (red), CS- (blue), the US (yellow) and CS discrimination (black). Single data points represent pairs of single trials at T0 and T1 averaged across participants. Note that no US was presented during extinction training and hence, no reliability of the US is shown in (B).

#### BOLD fMRI

In stark contrast to the low overlap of individual level activation (see Table 2A), the overlap at the group level was rather high with 62 % for the whole brain (Jaccard) for CS discrimination during acquisition training (see Table 2B). Similar to what was observed for overlap at the individual level, a much lower overlap for extinction training as compared to acquisition training was observed for the whole brain (4.60 % overlap) and all ROIs (all close to zero).

### Cross-phases predictability of conditioned responding

Finally, we investigated if responding in any given experimental phase predicted responding in subsequent experimental phases. To this end, simple linear regressions with robust standard errors were computed for both SCRs as well as fear ratings and all data specifications (see Figure 5 and Supplementary Tables 7 and 8). To approximate these analyses, correlations of patterns of BOLD brain activation between experimental phases were calculated (see Figure 6).

**Figure 5.**
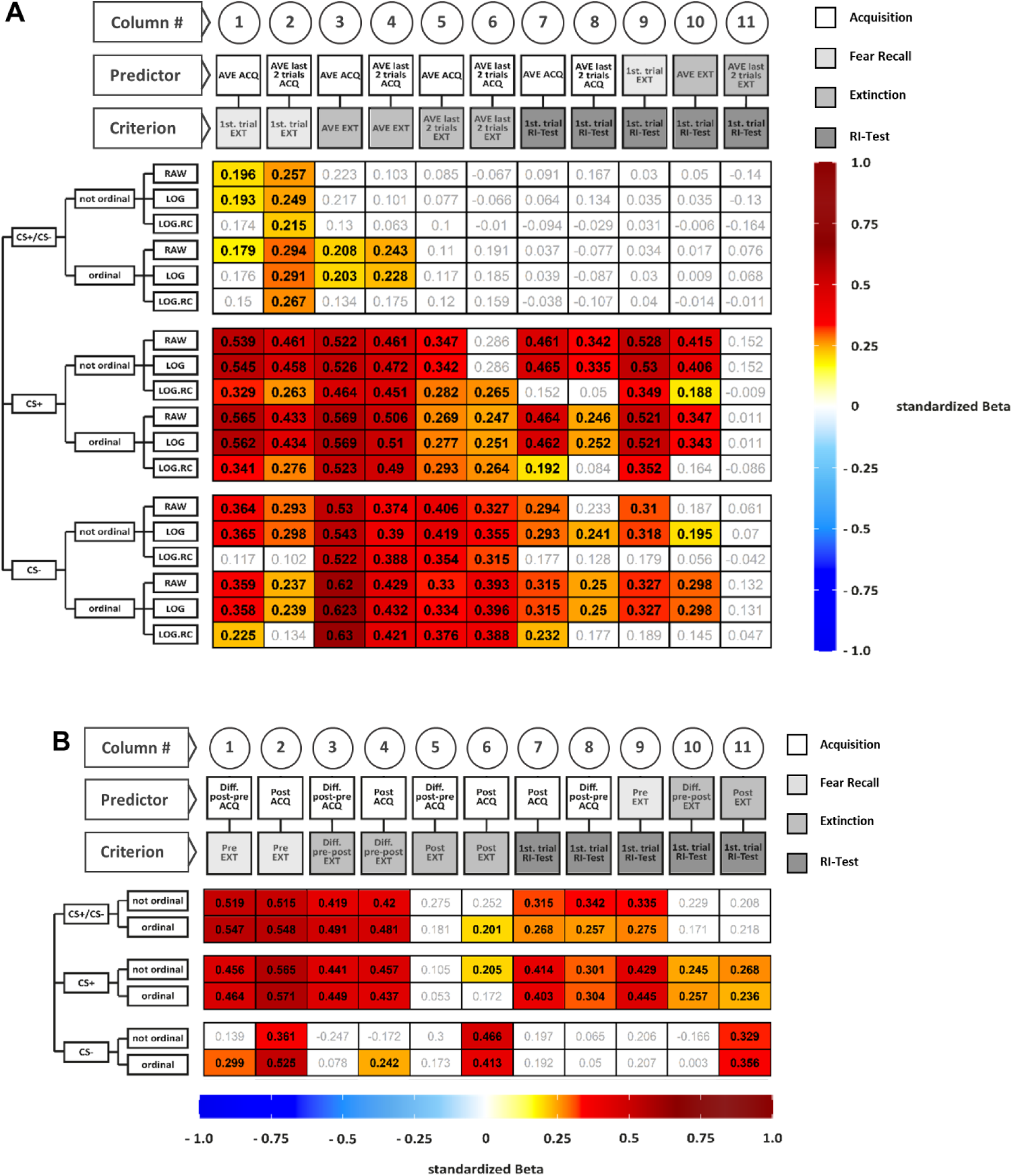
Illustration of standardized betas derived from regressions including SCRs (A) and fear ratings (B) for all data specifications. Colored cells indicate statistical significance of standardized betas, non-colored cells indicate non-significance. Standardized betas are color-coded for their direction and magnitude showing positive values from yellow to red and negative values from light blue to dark blue. Darker colors indicate higher betas. Tables containing regression parameters beyond the standardized betas depicted in Figure 5A and Figure 5B are presented in Supplementary Tables 9 and 10. AVE = average, LOG = log-transformed data, LOG.RC = log-transformed and range corrected data, not ordinal = not ordinally ranked data, ordinal = ordinally ranked data.

**Figure 6.**
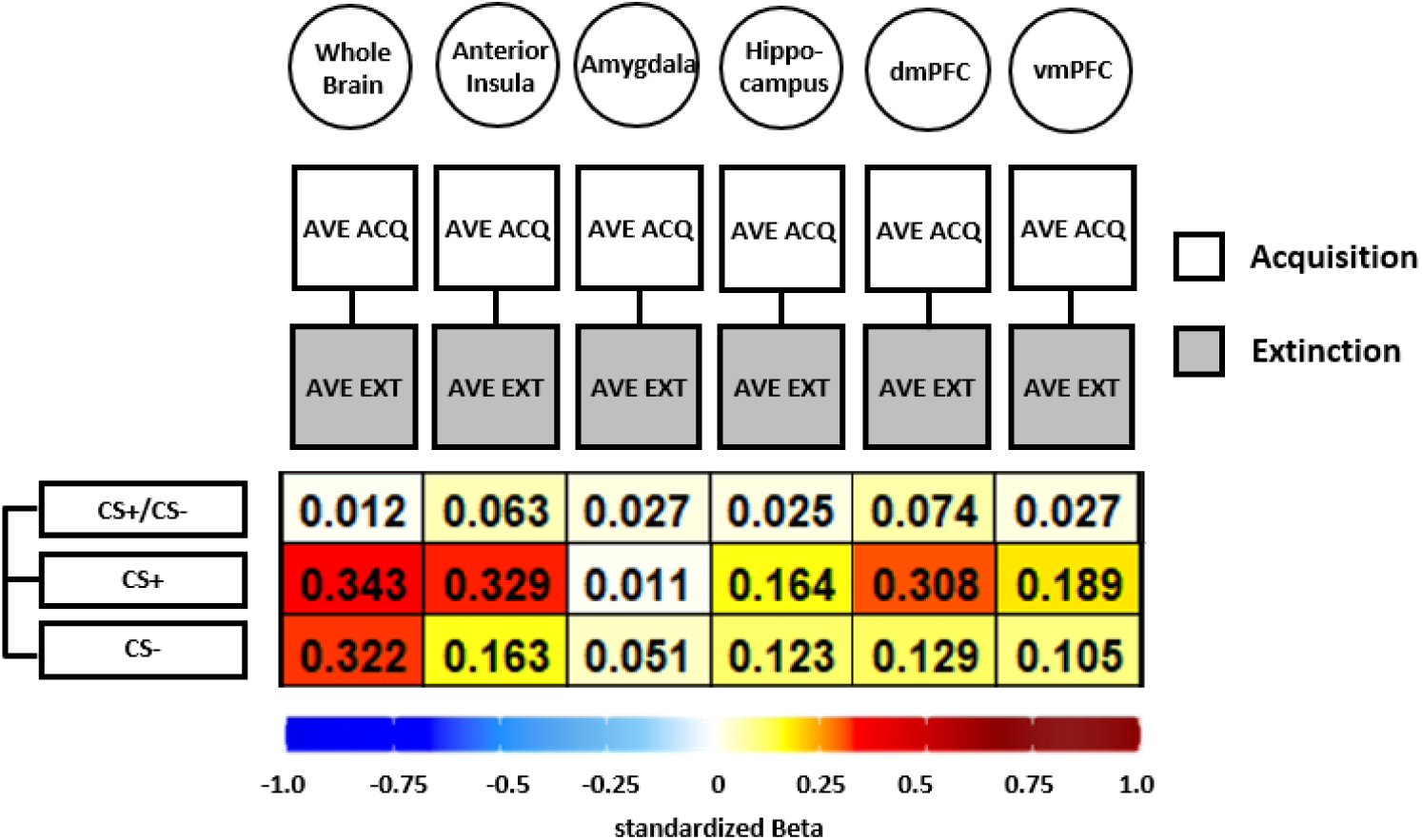
Illustration of standardized betas derived from correlation analyses between brain activation patterns during acquisition and extinction training in different ROIs and different data specifications. Standardized betas are color-coded for their direction and magnitude showing positive values from yellow to red and negative values from light blue to dark blue. Darker colors indicate higher betas.

#### SCR

Stronger CS discrimination in SCRs during (delayed) fear recall (i.e., first trial of extinction training) was significantly predicted by both average and end-point performance (i.e., last two trials) during acquisition training in most data specifications (Figure 5A, columns 1 and 2). In contrast, average CS discrimination during extinction training was significantly predicted only by acquisition training performance if data were ordinally ranked (columns 3 and 4). Strikingly, all predictions of extinction end-point performance (columns 5 and 6) as well as performance at reinstatement-test (columns 7 - 11) were non-significant – irrespective of phase operationalizations and data transformation.

The majority of predictions of SCRs to the CS+ and CS- were significant with few exceptions (see white cells in Figure 5A) - irrespective of experimental phases, their operationalization and data transformation. The non-significant regressions for the most part involved log-transformed and range corrected data. Strikingly, extinction end-point performance never predicted performance at reinstatement-test - irrespective of data transformation (column 11).

#### Fear ratings

Higher ratings for the CS+ as well as higher CS discrimination during acquisition training significantly predicted higher CS+ ratings and CS discrimination at fear recall (Figure 5B, columns 1 and 2), extinction training (columns 3 and 4) and at reinstatement-test (columns 7 - 8) and higher responding to the CS+ and higher CS discrimination at fear recall significantly predicted higher responding at reinstatement-test (column 9) - irrespective of data transformations. In contrast, predictions of CS discrimination and CS+ ratings after extinction training were mostly non-significant (columns 5 and 6). Higher CS+ ratings during extinction training significantly predicted higher ratings at reinstatement-test which was not true for CS discrimination (columns 10 and 11).

Higher CS- ratings after acquisition significantly predicted higher CS- ratings at fear recall as well as after extinction training and CS- ratings after extinction significantly predicted the performance at reinstatement-test - irrespective of ranking of the data (columns 2, 6 and 11). Furthermore, when based on ordinally ranked data, the difference between ratings prior to and after acquisition predicted CS- ratings at fear recall and CS- ratings after acquisition training predicted the difference between CS- ratings prior to and after extinction (column 1 and 4). All other predictions were non-significant.

In sum, all significant predictions observed were positive with weak to moderate associations and indicate that higher responding in preceding phases predicted higher responding in subsequent phases for both SCRs and fear ratings.

#### fMRI

In short, all associations were positive, showing that higher BOLD response during acquisition was associated with higher BOLD responding during extinction training (see Figure 6). However, the standardized beta coefficients are mostly below or around 0.3, indicating weak associations for all ROIs and CS specifications that were near absent for CS discrimination, CS+ and CS-. Analysis of CS+ and CS- data was included here as the analysis is based on beta maps and not T-maps (as in previous analyses) where a contrast against baseline is not optimal.

#### Cross-phases predictability depends on data specifications

Pooled across all other data specifications, some interesting patterns can be extracted: First, standardized betas were significantly lower for raw (*t*(65) = 8.08, *p* < .001, *d* = 0.99) and log-transformed (*t*(65) = 8.26, *p* < .001, *d* = 1.02) as compared to log-transformed and range corrected SCRs while standardized betas derived from the former did not differ significantly (*t*(65) = -0.26, *p* = .794, *d* = -0.03). Second, standardized betas derived from ranked and not-ranked analyses were comparable for fear ratings (*t*(32) = 1.26, *p* = .218, *d* = 0.22) and but not for SCRs with significantly higher betas for non-ranked as opposed to ranked SCRs (*t*(98) = 2.37, *p* = .020, *d* = 0.24). Third, standardized betas for CS discrimination were significantly lower than for CS+ and CS- for both SCRs (CS+: *t*(65) = -15.31, *p* < .001, *d* = -1.88 and CS-: *t*(65) = -12.34, *p* < .001, *d* = -1.52) and BOLD fMRI (CS+: *t*(5) = -3.72, *p* = .014, *d* = -1.52 and CS-: *t*(5) = -2.62, *p* = .047, *d* = -1.07), while for ratings, standardized betas for CS discrimination were higher than for the CS- (*t*(21) = 3.11, *p* = .005, *d* = 0.66) and comparable to those for the CS+ (*t*(21) = -0.57, *p* = .572, *d* = -0.12). Furthermore standardized betas were larger for the CS+ than for the CS- for SCRs (*t*(65) = 3.79, *p* < .001, *d* = 0.47) and ratings (*t*(21) = 3.12, *p* = .005, *d* = 0.67) and comparable for BOLD fMRI (*t*(5) = 2.17, *p* = .082, *d* = 0.89). Fourth, standardized betas derived from regressions predicting fear recall were significantly higher than for reinstatement-test for both SCRs (*t*(124) = 4.35, *p* < .001, *d* = 0.86) and fear ratings (*t*(40) = 5.15, *p* < .001, *d* = 1.76).

## Discussion

Our work follows recent calls for increasing attention to measurement reliability as a crucial determinant of statistical power in psychology and the neurosciences (Fröhner, Teckentrup, Smolka, & Kroemer, 2019; Hedge, Powell, & Sumner, 2018; Moriarity & Alloy, 2021; Zuo, Xu, & Milham, 2019). Measurement reliability is of key importance when addressing clinically-relevant questions involving individual-level predictions such as predicting treatment success from extinction (e.g., Lange et al., 2020; Scheveneels, Boddez, Vervliet, & Hermans, 2016) or biomarkers (Cano-Catala et al., 2021). Not only translational questions call for measurement tools that are sufficiently stable within a person over time, but also correlations with a third (individual difference) variable (Nunnally, 1970; Vul, Harris, Winkielman, & Pashler, 2009) because reliability puts an upper bound to the maximally observable correlation (i.e., the correlation approaches 1 only if both measures are perfectly reliable). However, in fear conditioning research only little is known about stability over time (i.e, longitudinal reliability, often referred to as test-retest reliability) and almost nothing is known about cross-sectional reliability and predictability across experimental phases.

Here, we aimed to fill this gap by systematically scrutinizing the reliability of cross-session as well as within-session responding. To this end, we investigated longitudinal reliability at the individual level which reflects the temporal stability of individual responding. Previous work has mostly focused on longitudinal reliability at the individual level as assessed for instance by ICCs (see Supplementary Table 1; Fredrikson, Annas, Georgiades, Hursti, & Tersman, 1993; Ridderbusch et al., 2021; Torrents-Rodas et al., 2014; Zeidan et al., 2012). Here we complement this traditional approach with i) analyses of response similarity and ii) the degree of overlap of individual-level brain activation patterns. Furthermore, we extend this focus by exploring longitudinal reliability at the group level as well as cross-sectional reliability.

In addition to reliability, we also directly investigated predictability of responding from one experimental phase to subsequent experimental phases. We took heterogeneous methodological approaches in the literature into account by systematically employing different data specifications (i.e., phase operationalizations, transformations).

Generally, longitudinal group-level reliability was good or acceptable for SCRs and the BOLD response while longitudinal individual-level reliability as assessed by ICCs, pattern similarity and individual-level BOLD activation overlap was generally poor across outcome measures and data specifications - particularly during extinction training. This is in line with previous work in fear conditioning (Fredrikson, Annas, Georgiades, Hursti, & Tersman, 1993; Ridderbusch et al., 2021; Torrents-Rodas et al., 2014; Zeidan et al., 2012) that also reported poor to moderate longitudinal individual-level reliability - in all three outcome measures (SCRs, fear ratings, BOLD fMRI) and across experimental phases.

Our complementary analyses beyond traditional ICCs yielded that for SCRs within- and between-subject similarity was comparable. This indicates that responses of one individual at baseline T0 were not more similar to responses of the same individual at T1 than compared to others at T1. For BOLD fMRI, however, within-subject similarity was higher than between-subject similarity during acquisition but not extinction training, indicating that acquisition-related individual BOLD activation patterns at T0 were more similar to their own activation patterns at T1 than to other individuals’ activation patterns. Hence, while traditional statistics of longitudinal reliability (i.e., ICCs) for SCR and BOLD fMRI during acquisition training were rather poor, the significantly higher similarity of neural activity within one individual as compared to others indicates that BOLD fMRI might be more sensitive to detect similarity at individual-level responses within participants than SCRs - maybe due to the reliance on a more spatial (i.e., voxel-by-voxel) than temporal (i.e., trial-by-trial) level as for SCRs.

It is noteworthy that longitudinal reliability at the individual level was slightly better for log-transformed and at the same time range-corrected SCRs (as opposed to raw and only log-transformed data) while - in contrast to what has been shown for other paradigms and outcome measures (Baker et al., 2021; see also https://shiny.york.ac.uk/powercontours/) - an increasing number of trials included in the calculation of ICCs did not generally improve reliability. Together, this suggests that longitudinal reliability at the individual level is relatively stable across different data transformations and paradigm specifications (e.g., number of trials within the range used here, i.e, 1 to maximum 14) which is important information facilitating the integration of previous work using different time intervals, reliability indices and paradigms (Fredrikson, Annas, Georgiades, Hursti, & Tersman, 1993; see Supplementary Table 1; Ridderbusch et al., 2021; Torrents-Rodas et al., 2014; Zeidan et al., 2012) as well as for optimization of paradigms for future work.

In stark contrast to this generally low individual-level longitudinal reliability, we observed robust longitudinal reliability at the group level for both SCRs and BOLD fMRI between both time points with substantial (i.e., up to 62%) overlap in group-level BOLD fMRI activation pattern (whole brain and ROI-based) as well as substantial (i.e., up to 76%) explained variance at T1 by variance at T0 for SCRs. However, this was generally only true for acquisition but not extinction training.

Reports regarding this discrepancy between group-level and individual-level longitudinal reliability were recently highlighted for a number of (classic) experimental paradigms (Fröhner, Teckentrup, Smolka, & Kroemer, 2019; Hedge, Powell, & Sumner, 2018; Herting, Gautam, Chen, Mezher, & Vetter, 2018; Plichta et al., 2012). Our results add fear conditioning and extinction as assessed by SCRs and BOLD fMRI to this list and have important implications for translational questions aiming for individual-level predictions - particularly since findings obtained at a group level are not necessarily representative for any individual within the group (Fisher, Medaglia, & Jeronimus, 2018). Additionally, we extend these methodological findings regarding long-term individual-level predictions by an empirical investigation of predictability of individual-level responding between experimental phases.

We observed significant weak to moderate associations between responding in different experimental phases for SCR, fear ratings and BOLD fMRI revealing that higher responses in previous phases were generally associated with higher responses in subsequent phases in all outcome measures. However, a remarkable amount of predictions were non-significant - which was particularly true for CS discrimination in SCRs and BOLD. This may be explained by difference scores (i.e., CS+ minus CS-) being generally less reliable (Lynam, Hoyle, & Newman, 2006) due to a subtraction of meaningful variance (Moriarity & Alloy, 2021) particularly in highly-correlated predictors (Thomas & Zumbo, 2012).

Mixed findings in the literature support both the independence of conditioned responding in different experimental phases (Bouton, García-Gutiérrez, Zilski, & Moody, 2006; Plendl & Wotjak, 2010; Prenoveau, Craske, Liao, & Ornitz, 2013; Shumake, Furgeson-Moreira, & Monfils, 2014) but also their dependence - particularly in clinical samples (Foa et al., 1983; Rauch, Foa, Furr, & Filip, 2004; Rothbaum et al., 2014; Smits, Rosenfield, Otto, Marques, et al., 2013; Smits, Rosenfield, Otto, Powers, et al., 2013). These diverging findings in experimental and clinical studies might reveal a translational gap. However, an alternative explanation might be revealed by our systematic investigation covering multiple operationalizations and data specifications. More precisely, our work may suggest that the strengths of associations between responding in different phases depended on the specific outcome measure and its specifications (e.g., responses specified as CS discrimination, CS+ or CS-). Yet another explanation - in particular for predictions spanning a 24h delay in experimental phases - might be that individual differences in consolidation efficacy may underlie differences in predictability which has implications for the common practice of correcting responses during one experimental phase for responding during preceding experimental phases (discussed in Lonsdorf, Merz, & Fullana, 2019).

Importantly, taken together with our observation of robust cross-sectional reliability (see also Fredrikson, Annas, Georgiades, Hursti, & Tersman, 1993), this pattern of findings suggests that individual-level predictions at short intervals may be feasible, but problematic over longer time-periods as suggested by the limited stability over time in our data.

Yet, before discussing implications of our results in detail, some reflections on potential (methodological) reasons for i) low individual-level but robust group-level reliability and ii) on the role of time interval lengths deserve attention:

First, low individual level but robust group level longitudinal reliability might be (in part) due to different averaging procedures which impacts error variance (Kennedy et al., 2021) with group-level data based strongly on aggregated data compared to individual-level data.

Second, different operationalizations of the same measurement might have different reliability (Kragel, Han, Kraynak, Gianaros, & Wager, 2021). For instance, amygdala habituation has been shown to be a more reliable measure than average amygdala activation (Plichta et al., 2014) and more advanced analytical approaches such as intra-individual neural response variability (Månsson et al., 2021) and multivariate imaging techniques (Kragel, Han, Kraynak, Gianaros, & Wager, 2021; Marek et al., 2020; Noble, Scheinost, & Constable, 2021; Visser, Bathelt, Scholte, & Kindt, 2021) have been suggested to have better (longitudinal) reliability than more traditional analyses approaches. Finally, longitudinal reliability refers to measurements obtained under the same conditions and hence it is both plausible and well established that higher reliability is observed at short test-retest intervals (see also Noble, Scheinost, & Constable, 2021). Longer intervals are more susceptible to true changes of the measurand - for instance due to environmental influences such as seasonality, temperature, hormonal status or life events (see Specht, Egloff, & Schmukle, 2011; Vaidya, Gray, Haig, & Watson, 2002). Indeed most longitudinal reliability studies in the fMRI field used shorter intervals (< 6 month; see Elliott et al., 2020; Noble, Scheinost, & Constable, 2021) than our 6-month interval and hence our results should be conceptualized as longitudinal stability rather than a genuine test-retest reliability. The good to excellent cross-sectional reliability speaks against excessive noisiness inherent to our measures and suggests a true change of the measurand during our retest interval and hence a potentially stronger state than trait dependency.

What do our findings imply? Poor measurement reliability has major consequences as it results in smaller effect sizes and lower statistical power for individual difference questions and longitudinal studies (Elliott et al., 2020). Fear conditioning research has been highlighted as the “best current opportunity for translating neuroscience discoveries into clinical applications” (cf. Anderson & Insel, 2006, p. 319) and most of the pressing translational questions are based on individual-level predictions such as predicting treatment success. Our results, however, suggest that measurement reliability may only allow for individual-level predictions for (very) short (as in our cross-phases predictability analysis) but not longer time intervals (such as our 6 months retest interval). Importantly, however, good group-level reliability also allows for group-level predictions over longer time intervals. A potential solution to make use of both good group level reliability and individual-level predictions might be the implementation of homogenous (latent) subgroups (e.g., Galatzer-Levy, Bonanno, Bush, & Ledoux, 2013) which might be a promising future avenue.

We argue that we may need to take a (number of) step(s) back and develop paradigms and data processing pipelines explicitly tailored to individual difference research and focus more strongly on measurement reliability. More precisely, multiverse-type investigations (Steegen, Tuerlinckx, Gelman, & Vanpaemel, 2016) that systematically scrutinize the impact of several alternative and equally justifiable processing and analytical decisions in a single dataset (Kuhn, Gerlicher, & Lonsdorf, 2021 PREPRINT) - as also done here for transformations and number of trials - may be helpful to achieve this overarching aim. This could be complemented by systematically varying design specifications (Harder, 2020) which are extensively heterogeneous in fear conditioning research (Lonsdorf et al., 2017). Calibration approaches, as recently suggested (Bach, Melinščak, Fleming, & Voelkle, 2020) follow a similar aim.

Such work on measurement questions can (often) be done cost and resource effective in existing or openly available data - which, however, requires cross-lab data sharing and data management homogenization plans. Devoting resources and funds to measurement optimization is a valuable investment into the prospect of this field contributing to improved mental health (Moriarity & Alloy, 2021) and to resume the path to successful translation from neuroscience discoveries into clinical applications.

## Materials and methods

### Pre-registration

This project has been pre-registered on the Open Science Framework (OSF) (August 03, 2020; retrieved from https://doi.org/10.17605/OSF.IO/NH24G). Deviations from the pre-registered protocol are made explicit in brief in the methods section and reasons are specified in Supplementary Table 2 as recommended by Nosek, Ebersole, DeHaven, and Mellor (2018), who notes that such deviations are common and occur even in the most predictable analysis plans.

### Participants

Participants were selected from a large cohort providing participants for subsequent studies in the Collaborative Research Center CRC 58. Participants from this sample were recruited for this study through a phone interview. Only healthy individuals between 18 and 50 years of age without a history of childhood trauma according to the Childhood Trauma Questionnaire (CTQ; critical cutoffs as identified by Bernstein et al., 2003). Additional exclusion criteria were claustrophobia, cardiac pacemaker, non-MR-compatible metal implants, brain surgery, left handedness, participation in pharmacological studies within the past 2 weeks, medication except for oral contraceptives, internal medical disorders, chronic pain, neurological disorders, psychiatric disorders, metabolic disorders, acute infections, complications with anaesthesia in the past and pregnancy. Participants were right-handed and had normal or corrected to normal vision. All participants gave written informed consent to the protocol which was approved by the local ethics committee (PV 5157, Ethics Committee of the General Medical Council Hamburg). The study was conducted in accordance with the Declaration of Helsinki. All participants were naïve to the experimental setup and received a financial compensation of 170 € for completion of experiments at both time points (T0 and T1).

The total sample consisted of 120 participants (female*_N_* = 79, male*_N_* = 41, age*_M_* = 24.46, age*_SD_* = 3.73, age*_range_* = 18 - 34). At T0 on day 1 and day 2, in total 13 participants were excluded due to technical issues (day 1: *N* = 0; day 2: *N* = 3), deviating protocols (day 1: *N* = 2; day 2: *N* = 0) and SCR non-responding (day 1: *N* = 3; day 2: *N* = 5, see below for definition of ‘non-responding’). Accordingly, the final data set for the cross-sectional analysis of T0 data consists of 107 subjects (female*_N_* = 70, male*_N_* = 37, age*_M_* = 24.30, age*_SD_* = 3.68, age_range_ = 18 - 34). 84.11% of these participants were aware and 6.54% were unaware of CS-US contingencies. The remaining 9.35% subjects uncertain of the CS-US contingencies were classified as semi-aware. CS-US contingency awareness of participants was assessed with a standardized post-experimental awareness interview (adapted from Bechara et al. (1995)). On average, the US aversiveness was rated on day 1 with a value of 19.82 (*SD* = 3.28) and on day 2 with a value of 16.46 (*SD* = 4.75) on a visual analogue scale (VAS) ranging from 0 to 25. The US intensity was 8.04 mA (*SD* = 8.28) on average. Averaged STAI-S (Strait-Trait Anxiety Inventory - State; Spielberger, 1983) scores were 35.38 (*SD* = 5.26) on day 1 and 35.57 (*SD* = 6.69) on day 2.

At T1, 16 subjects were excluded due to technical issues (day 1: *N* = 1; day 2: *N* = 1), deviating protocols (day 1: *N* = 3; day 2: *N* = 0) and SCR non-responding (day 1: *N* = 5; day 2: *N* = 6; see below for definition of ‘non-responding’). 88.73% of these participants were aware and 1.41% were unaware of CS-US contingencies. The remaining 9.86% were classified as semi-aware. US aversiveness was rated with *M* = 19.96 (*SD* = 2.99) on day 1 and with *M* = 17.73 (*SD* = 3.90) on day 2 (VAS = 0 - 25). On average, the US intensity amounted to 9.76 mA (*SD* = 13.18). Averaged STAI-S scores were 36.33 (*SD* = 6.09) on day 1 and 35.83 (*SD* = 7.10) on day 2.

Additionally, twenty participants dropped out between T0 and T1 leaving 71 subjects for longitudinal analyses (female*_N_* = 41, male*_N_* = 30, age*_M_* = 24.63, age*_SD_* = 3.77, age*_range_* = 18 - 32).

### Experimental design

Here, we re-analyzed pre-existing data that are part of a larger longitudinal study that spanned six time points. Part of the data has been used previously in method focused work (Kuhn, Gerlicher, & Lonsdorf, 2021; Lonsdorf, Gerlicher, Klingelhöfer-Jens, & Krypotos, 2021; Lonsdorf et al., 2019) and work assessing the association between physiological and subjective measures of fear acquisition as well as extinction and brain morphology (Ehlers, Nold, Kuhn, Klingelhöfer-Jens, & Lonsdorf, 2020). In the current study, we included data from a two-day fear conditioning experiment which were collected at two time points (T0 and T1) six months apart. The two-day experimental procedure and the stimuli were identical at both time points. Measures acquired during the full longitudinal study that are not relevant for the current work such as questionnaires, hair and salivary cortisol are not described in detail here.

### Experimental protocol and stimuli

The protocol consisted of a habituation and a fear acquisition training phase on day 1 and an extinction training, reinstatement, and reinstatement-test phase on day 2. Acquisition and extinction training included 28 trials each (14 CS+/ 14 CS-), habituation and the reinstatement-test phase 14 trials each (7 CS+/ 7 CS-). Acquisition training was designed as delay conditioning with the US being presented 0.2 s before CS+ offset with 100% reinforcement rate (i.e., all CS+ presentations followed by the US). CSs were two light grey fractals (RGB [230, 230, 230]), 492*492 pixels) presented in a pseudo-randomized order, with no more than two identical stimuli in a row, for 6 - 8 s (mean: 7 s). During the inter-trial interval (ITI), a white fixation cross was shown for 10 to 16 s (mean: 13 s). The reinstatement consisted of three trials with a duration of 5 s each presented after a 10 s ITI. Reinstatement USs were delivered 4.8 s after each trial onset. The reinstatement was followed by a 13 s ITI before the next CS was presented during reinstatement-test. All stimuli were presented on a grey background (RGB [100, 100, 100]) by using Presentations software (2010) (Version 14.8, Neurobehavioral Systems, Inc., Albany California, USA) keeping the context constant to avoid renewal effects (Haaker, Golkar, Hermans, & Lonsdorf, 2014). Visual stimuli were identical for all participants, but allocation to CS+/CS- and CS-type of the first trial of each phase were counterbalanced across participants.

The electrotactile US consisted of a train of three 2 ms electro-tactile rectangular pulses with an interpulse interval of 50 ms generated by a Digitimer DS7A constant current stimulator (Welwyn Garden City, Hertfordshire, UK) and was administered to the back of the right hand of the participants through a 1 cm diameter platinum pin surface electrode. The electrode was attached between the metacarpal bones of the index and middle finger. The US was individually calibrated in a standardized stepwise procedure controlled by the experimenter aiming at an unpleasant, but still tolerable level rated by the participants between 7 and 8 on scale from zero (= stimulus was not unpleasant at all) to 10 (= stimulus was the worst one could imagine within the study context). Participants were, however, not informed that we aimed at a score of 7 to 8.

### Outcome measures

#### Skin conductance responses

SCRs were acquired continuously during each phase of conditioning using a BIOPAC MP 100 amplifier (BIOPAC Systems, Inc., Goleta, California, USA) and Spike 2 software (Cambridge Electronic Design, Cambridge, UK). For analogue to digital conversion, a CED2502-SA was used. Two self-adhesive hydrogel Ag/AgCl-sensor recording SCR electrodes (diameter = 55 mm) were attached on the palm of the left hand on the distal and proximal hypothenar. A 10 Hz lowpass filter and a gain of 5 Ω were applied. Data were recorded at 1000 Hz and later down-sampled to 10 Hz. Subsequently, SCRs were scored semi-manually using the custom-made computer program EDA View (developed by Prof. Dr. Matthias Gamer, University of Würzburg). The program is used to quantify the SCR amplitude based on the trough-to-peak method (TTP) with the trough occurring at 0.9 - 3.5 s after CS onset and 0.9 -2.5 s after US onset (Boucsein et al., 2012; Sjouwerman & Lonsdorf, 2019). The maximum rise time was set to maximally 5 s (Boucsein et al., 2012) unless the US occurred earlier. SCRs confounded by recording artifacts due to technical reasons, such as electrode detachment or responses moving beyond the sampling window, were discarded and scored as missing values. SCRs smaller than 0.01 μS within the defined time window were defined as zero-responses. Participants with zero responses to the US in more than two-thirds (i.e., more than 9 out of 14) of US acquisition trials were classified as non-responders on day 1. On day 2, non-responding was defined as no response to any of the three reinstatement USs.

SCR data were prepared for response quantification by using MATLAB (2016) version R2016b. No learning could have possibly taken place during the first CS presentations as the US occurred only after the CS presentation. Consequently, the first CS+ and CS- trial during acquisition training were excluded from analyses. Hence, a total of 26 trials (13 differential SCRs) for the acquisition training phase were included in the analyses. For US analyses, all 14 trials were entered into analyses.

Similarly, responses to the first CS+ and CS- during extinction training have to be considered a 24h delayed test of fear recall as no extinction learning could have taken place. Hence, the first trial and the remaining trials of the extinction were analyzed separately. CS discrimination was computed by subtracting (averaged) CS- responses from (averaged) CS+ responses.

#### Fear ratings

Fear ratings to the CSs were collected prior to and after acquisition and extinction training as well as after the reinstatement-test. Participants were asked “how much stress, fear and tension” they experienced when they last saw the CS+ and CS-. After reinstatement-test, ratings referred to i) the first CS presentation per CS-type directly after reinstatement as well as ii) the last CS presentation during reinstatement-test. After acquisition training and the reinstatement-test, subjects were also asked how uncomfortable they experienced the US itself. All ratings were given on a VAS ranging from zero (answer = none) to 100 (answer = maximum). For analyses, the rating scale was reduced to 0 - 25. Participants had to confirm the ratings via button press. A lack of confirmation resulted in exclusion of the trial from analyses. CS discrimination was computed by subtracting CS- from CS+ ratings.

### BOLD fMRI: data acquisition, preprocessing and first level analysis

The inclusion of BOLD fMRI data was not initially planned and is included here as an additional non-pre-registered outcome measure.

#### Data Acquisition

Functional data was acquired with a 3 Tesla PRISMA whole body scanner (Siemens Medical Solutions, Erlangen, Germany) using a 64-channel head coil and an echo planar imaging (EPI) sequence (repetition time (TR): 1980 ms, echo time (TE): 30 ms, number of slices: 54, slice thickness: 1.7 mm (1 mm gap), field of view = 132 x 132 mm). T1- weighted structural images were acquired using a magnetization prepared rapid gradient echo (MPRAGE) sequence (TR: 2300 ms, TE: 2.98 ms, number of slices: 240, slice thickness: 1 mm, field of view = 192 x 256 mm).

#### Preprocessing

fMRI data analysis was performed using SPM12 (Wellcome Department of Neuroimaging, London, United Kingdom) and MATLAB (2019). Preprocessing included realignment, coregistration, normalization to a group-specific DARTEL template and smoothing (6 mm FWHM).

#### First-level analysis

Regressors for the first level analysis of acquisition training data included separate regressors for the first CS+ and CS- trials and the remaining CS+ and CS- trials because no learning could have occurred at the first presentation of the CSs. Nuisance regressors included habituation trials, US presentation, fear ratings and motion parameters. Likewise, separate regressors for the first CS+ and CS- trials of extinction (because no extinction has taken place yet) as well as the remaining CS+ and CS- trials were included as regressors of interest in the first level analysis of extinction data acquired on day 2, while US, rating onset and motion parameters were included as regressors of no interest. No second-level analysis was completed in the current study, instead different analyses were carried out based on first level models as further detailed in the statistical analysis section.

#### Regions of Interest

A total of five regions of interest (ROIs; i.e., bilateral anterior insula, amygdala, hippocampus, dorsomedial prefrontal cortex [dmPFC], ventromedial prefrontal cortex [vmPFC]) were included in the current study. Amygdala and hippocampus anatomical masks were extracted from the Harvard-Oxford atlas (Desikan et al., 2006) at a maximum probability threshold of 0.5. The anterior insula was defined as the overlap between the thresholded anatomical mask from the Harvard Oxford atlas (threshold: 0.5) and a box of size 60 x 30 x 60 mm centered around MNIxyz = 0, 30, 0. As previously reported (Lonsdorf, Haaker, & Kalisch, 2014), the cortical ROIs dmPFC and vmPFC were created by using a box of size 20 x 16 x 16 mm centered on peak coordinates identified in prior studies of fear learning (dmPFC: MNIxyz = 0, 43, 29, Kalisch et al. (2009), Milad et al. (2009); vmPFC: MNIxyz = 0, 40, -12, e.g., Kalisch et al. (2006), Milad et al. (2007)) with the x coordinate set to 0 to obtain masks symmetric around the midline. All analyses of BOLD fMRI as described below were conducted separately not only for the whole brain but also for these five selected ROIs.

### Statistical analyses

For a comprehensive overview of which analysis was carried out for which outcome measures, stimuli, phases and data transformations, see Table 1.

#### Cross-sectional reliability

We assessed the cross-sectional reliability of SCRs for both time points and experimental phases separately (for details, see also Table 1): trials of the respective time point and phase were split into odd and even trials (i.e., odd-even approach) and averaged for each individual subject. Averaged odd and even trials were then correlated by using Pearson’s correlation coefficient. To obtain a rather conservative result, we refrained from applying the Spearman-Brown prophecy formula. Calculations of cross-sectional reliability were not possible for fear ratings and BOLD fMRI due to the limited number of data points for fear ratings and an experimental design that did not allow for a trial-by-trial analysis of BOLD fMRI data. Cross-sectional reliability was interpreted using benchmarks for unacceptable (< 0.5), poor (> 0.5 but < 0.6), questionable (> 0.6 but < 0.7), acceptable (> 0.7 but < 0.8), good (> 0.8 but < 0.9) and excellent (≥ 0.9) (Kline, 2013).

#### Longitudinal reliability at the individual and group level

While cross-sectional reliability indicates the extent to which all items of a test or - here, trials of an experimental phase - measure the same construct (Revelle, 1979), longitudinal reliability reflects the variability across two or more measurements of the same individual under the same conditions and is therefore indicative of the degree of correlation and agreement between measurements (Koo & Li, 2016). For calculations of longitudinal reliability, we included data from both time points T0 and T1 from the same experimental phase. To capture different aspects of longitudinal reliability, we chose a dual approach of calculating longitudinal reliability at both i) the individual level and ii) at the group level (for details see also Table 1). To this end, longitudinal reliability at the individual and group level indicates to which extent responses within the same individual and within the group as a whole are stable over time. More precisely, whereas longitudinal reliability at the individual level takes into account the individual responses of participants, which are then related across time points, reliability at the group level first averages the individual responses across the group and then relates them across time points. Reliability at the individual level inherently includes the group level, as it is calculated for the sample as whole, but the individual responses are central to the calculation. Contrarily, for reliability at the group level, the calculation is carried out using group averages.

**Reliability at the individual level** was investigated as i) ICCs encompassing both time points, ii) within- and between-subject similarity of individual trial-by-trial responding (i.e., SCRs) or BOLD fMRI activation patterns between time points and iii) as the degree of overlap of significant voxels between time points within an **individual** (for methodological details see below). **Reliability at the group level** was investigated as i) trial-by-trial group average SCRs and ii) the degree of overlap of significant voxels between time points within the **group as a whole** (for methodological details see below).

Assessments of cross-sectional reliability, within- and between-subject similarity, overlap at the individual and group level as well as longitudinal reliability of SCRs at a group level were not pre-registered but are included as they provide valuable additional and complementary information. Overlap and similarity analyses follow the methodological approach of Fröhner, Teckentrup, Smolka, and Kroemer (2019).

#### Longitudinal reliability at the individual level

##### Intra-class correlation coefficients

ICCs were determined separately for each experimental phase by including data from both time points T0 and T1. Generally, larger ICCs indicate higher congruency of within-subject responding between time points and increased distinction of subjects from each other (Noble, Scheinost, & Constable, 2021). Parsons, Kruijt, and Fox (2019) recommend the calculations of ICCs in cognitive-behavioral tasks through a two-way mixed-effects model of single rater type labeled ICC(2,1) (absolute agreement, in the following referred to as ICC_abs_) and ICC(3,1) (consistency, in the following referred to as ICC_con_) according to Shrout and Fleiss’s (1979) convention and to report their 95% confidence intervals (CIs). Due to their slightly different calculations, ICC_abs_ tends to be lower than ICC_con_ (see Table 1).

However, as the pre-registered mixed-effects approach resulted in non-convergence of some models for SCRs and ratings, we implemented an ANOVA instead of the mixed-effects approach to calculate ICC_abs_ and ICC_con_ (Shrout & Fleiss, 1979). To calculate ICCs for BOLD fMRI (additional not pre-registered analyses), the SPM based toolbox fmreli (Fröhner, Teckentrup, Smolka, & Kroemer, 2019) was used. BOLD fMRI ICCs were determined for each voxel and averaged across the whole brain and for selected ROIs.

Furthermore, we investigated whether or to what extent ICCs change when ICC calculations were based on different numbers of trials. To this end, we included (additional non-preregisted) analyses of trial-by-trial ICCs for SCRs in the supplementary material: First, ICCs were only computed for the first trial. Then, all subsequent trials of the respective phase were added stepwise to this first trial. After each step, trials were averaged and ICCs were calculated (see Supplementary Figures 3 to 8).

As suggested (Koo & Li, 2016), values less than 0.50 are interpreted as poor reliability, values between 0.50 and 0.75 as indicative of moderate reliability, values between 0.75 and 0.9 are interpreted as good reliability and values greater than 0.90 indicate excellent reliability. Note, however, that these values are rules of thumb rather than clear cut-offs.

##### Within- and between-subject similarity

Both ICCs and within-subject similarity indicate to which extent responses of the individual at one time point are comparable to responses of the same individual at a later time point. Both were calculated separately for each experimental phase by including data from both time points. There are, however, two main differences. Firstly, ICCs were calculated by decomposition of variances as applied for analysis of variance (ANOVA), whereas similarity was calculated as correlation of responses between both time points i) within one individual (within-subject similarity) and ii) between this individual and all other individuals (between-subject similarity). Secondly, while ICCs are interpreted in terms of absolute values using cutoffs that provide information on the quantity of longitudinal reliability, within-subject similarity was compared to between-subject similarity showing if responses of one subject at T0 were more similar to themselves at T1 than to responses of all others at T1.

The approach to the assessment of similarity was derived from the idea of representational similarity analysis (RSA) introduced by Kriegeskorte (2008) and previously used by Fröhner, Teckentrup, Smolka, and Kroemer (2019) for the comparison of fMRI BOLD activation patterns between different sessions.

Here, within-subject similarity was calculated by correlating (Pearson correlation coefficient) i) individual trial-by-trial SCRs and ii) the first-level response patterns of brain activation for CS discrimination (i.e. CS+ > CS-) of each individual subject between T0 and T1 resulting in one within-subject similarity value per subject (e.g., SCR acquisition trials of subject 1 at T0 were correlated with SCR acquisition trials of subject 1 at T1). Between-subject similarity was calculated by correlating trial-by-trial SCRs or the first-level response patterns of brain activation of each individual subject at T0 with those of all other individuals at T1 (e.g., SCR acquisition trials of subject 1 at T0 were correlated with SCR acquisition trials of subject 2 to 71 at T1). This resulted in 70 correlation coefficients for each subject. These correlation coefficients were then averaged to yield one correlation coefficient per subject as an indicator of between-subject similarity.

For comparisons of within- and between-subject similarity in SCR and BOLD fMRI, similarities were Fisher r-to-z transformed and compared by using paired t-tests or Welsh tests in cases where the assumption of equal variances was not met. Cohen’s d is reported as effect size.

Note that within-subject similarities of SCRs could not be calculated for participants with a single non-zero response at the same trial (e.g, trial 1) at both time points or only zero responses to the CS+ or CS- in one particular phase. This is because arrays that include only zeros can not be correlated and correlations of 1 (e.g., resulting from non-zero responses at the same trial at both time points) result in infinite Fisher r-to-z transformed correlations. Thus, different numbers of participants had to be included in the analyses of SCRs during acquisition (*N_CSdiscrimination_* = 65, *N_CS+_* = 62, *N_CS-_* = 56, *N_US_* = 71) and extinction training (*N_CSdiscrimination_* = 45, *N_CS+_* = 40, *N_CS-_* = 32).

##### Overlap at the individual level

For BOLD fMRI, overlap in individual subject BOLD activation patterns across both time points was calculated as a third indicator of reliability at the individual level. Thus, overlap was determined separately for experimental phases by including data from both time points T0 and T1. To this end, activation maps from first-level contrasts (here CS+ > CS- or CS discrimination) were compared such that the degree of overlap of significant voxels at a liberal threshold of *p_uncorrected_* < .01 between T0 and T1 was determined and expressed as the Dice and Jaccard coefficients (Fröhner, Teckentrup, Smolka, & Kroemer, 2019). Both coefficients range from 0 (no overlap) to 1 (perfect overlap), with the Jaccard index being easily interpretable as percent overlap (Fröhner, Teckentrup, Smolka, & Kroemer, 2019). While overlap reflects the degree of voxels activated at both time points, similarity measures (see above) are based on the correlation of activated voxels between time points and can be considered a continuous approach based on CS+ > CS- contrast specific beta values and not thresholded T-maps.

#### Longitudinal reliability at the group level

As opposed to longitudinal reliability at the individual level which indicates the stability of individual responses across time points, longitudinal reliability at the group level refers to the extent to which the group average responding is stable over time. Longitudinal reliability at the group level was calculated separately for experimental phases by including data from both time points T0 and T1.

We define longitudinal reliability at the group level i) for SCRs as the percentage of explained variance of group averaged trials at T1 by group averaged trials at T0 (i.e., R squared) and ii) for BOLD fMRI as the degree of overlap of group averaged activated voxels between both time points. Different analysis approaches were chosen as SCR and BOLD fMRI data are inherently different measures: trial-by-trial analyses in fMRI require slow-event related designs with long ITIs as well as fixed trial orders and ideally partial reinforcement rate to not confound CS and US responses (Visser et al., 2016). Hence, trial-by-trial analyses were not possible given our design and thus overlap at a group level was defined as overlap at voxel rather than at trial level.

For SCRs, simple linear regressions were computed with group averaged SCR trials at T0 as independent and group averaged SCR trials at T1 as dependent variable and R squared was extracted. This was done separately for experimental phases. Although the Pearson correlation coefficient is often calculated to determine longitudinal reliability, R squared, which like overlap can also be expressed as a percentage, appears closest to the concept of overlap of significant voxels at T0 and T1 as applied to BOLD fMRI data.

For overlap in BOLD fMRI at the group level, the degree of overlap of significant voxels between both time points was determined for aggregated group level activations instead of single subject level activation patterns (see ‘Overlap at the individual level’) and expressed using the Dice and Jaccard indices as described above.

#### Cross-phases predictability of conditioned responding

Simple linear regressions were calculated to assess the predictability of SCRs and fear ratings across experimental phases at T0. During data analysis, inspection of the data revealed heteroscedasticity. Therefore and deviating from the pre-registration, regressions with robust standard errors were calculated by using the HC3 estimator (Hayes & Cai, 2007). Two consecutive phases represent the independent and the dependent variable respectively, with the preceding phase as the independent variable and the following phase as the dependent variable. For SCR and fear ratings, standardized betas as derived from linear regressions are reported. In simple linear regression, as implemented here, standardized betas can be also interpreted as Pearson correlation coefficients.

For fMRI data, we adopted the cross-phases predictability analysis of SCR and fear ratings by calculating Pearson correlation coefficients between patterns of voxel activation (i.e. first level beta maps). Correlations were first calculated at the individual subject level and subsequently averaged.

Standardized betas (resulting from SCR and fear rating regressions) and correlation coefficients (resulting from BOLD fMRI correlational tests) were interpreted as demonstrating weak, moderate or strong associations between variables with values of < .40, ≥ .40. and ≥ .70 respectively (Dancey & Reidy, 2007). Tables containing regression parameters beyond the standardized betas depicted in Figure 5A and B are presented in the Supplement (see Supplementary Tables 9 and 10).

For SCR and fear rating predictions, we assessed if predictions differ in their strength or direction when they are summarized across certain data specifications (see Table 1). For BOLD fMRI, correlation coefficients were pooled across ROIs. T-tests or Welch tests in cases where the assumption of equal variances was not met were performed on individual Fisher r-to-z transformed standardized betas (SCR and fear ratings) or correlation coefficients (BOLD fMRI).

#### Data specifications

As the literature is characterized by heterogeneity with respect to which stimuli are included, how to operationalize experimental phases (i.e., number of trials included) and how to transform the data, we systematically employed different (with few exceptions) pre-registered data specifications here (see Table 1).

We included i) responses to the CS+, CS-, US and CS discrimination, ii) different phase operationalizations (i.e., number of trials), iii) different data transformations (none, log-transformed, log-transformed and range corrected) and iv) ordinally ranked vs. non-ranked data (to investigate whether the quality of predictions changes when the predictions were not based on the absolute values but on a coarser scale).

We acknowledge that the specifications used here are not intended to cover all potentially meaningful combinations as in a multiverse study (Lonsdorf, Gerlicher, Klingelhöfer-Jens, & Krypotos, 2021 PREPRINT; Steegen, Tuerlinckx, Gelman, & Vanpaemel, 2016) but can be viewed as a small manyverse.

For all statistical analyses described above, a level of *p* < .05 (two-sided) was considered significant. Since we were more interested in patterns of results and less in the result of one specific test, it was not necessary to correct for multiple comparisons. Moreover, multiverse approaches, as approximated in our study, are assumed to be insensitive to multiple comparisons (Lonsdorf, Gerlicher, Klingelhöfer-Jens, & Krypotos, 2021 PREPRINT).

For data analyses and visualizations as well as for the creation of the manuscript, we used R (Version 4.1.3; R Core Team, 2020) and the R-packages *apa* (Aust & Barth, 2020; Version 0.3.3; Gromer, 2020), *car* (Version 3.0.10; Fox & Weisberg, 2019; Fox, Weisberg, & Price, 2020), *carData* (Version 3.0.4; Fox, Weisberg, & Price, 2020), *cowplot* (Version 1.1.1; Wilke, 2020), *DescTools* (Version 0.99.42; Andri et mult. al., 2021), *dplyr* (Version 1.0.8; Wickham, François, Henry, & Müller, 2021), *effsize* (Torchiano, 2020), *flextable* (Version 0.6.10; Gohel, 2021), *gghalves* (Version 0.1.1; Tiedemann, 2020), *ggplot2* (Version 3.3.5; Wickham, 2016), *ggpubr* (Version 0.4.0; Kassambara, 2020), *ggsignif* (Version 0.6.3; Constantin & Patil, 2021), *gridExtra* (Version 2.3; Auguie, 2017), *here* (Version 1.0.1; Müller, 2020), *kableExtra* (Version 1.3.1; Zhu, 2020), *knitr* (Version 1.37; Xie, 2015), *lm.beta* (Version 1.5.1; Behrendt, 2014), *lmtest* (Version 0.9.38; Zeileis & Hothorn, 2002), *officedown* (Version 0.2.4; Gohel & Ross, 2022), *papaja* (Version 0.1.0.9997; Aust & Barth, 2020), *patchwork* (Version 1.1.0; Pedersen, 2020), *psych* (Version 2.0.9; Revelle, 2020), *reshape2* (Version 1.4.4; Wickham, 2007), *sandwich* (Zeileis, 2004, 2006; Version 3.0.1; Zeileis, Köll, & Graham, 2020), *stringr* (Version 1.4.0; Wickham, 2019), *tidyr* (Version 1.2.0; Wickham, 2020), *tinylabels* (Version 0.2.3; Barth, 2022), and *zoo* (Version 1.8.8; Zeileis & Grothendieck, 2005).

## Acknowledgements

The authors would like to thank Claudia Immisch for data collection, Mario Reutter for methodological discussions and comments on an earlier draft as well as Juliane Tkotz for supporting with reproducible manuscript writing. This work was supported by grants awarded by the German Research foundation to TBL (CRC 58 on “Fear, Anxiety and Anxiety Disorders,” Grant ID INST 211/633-2, Grant ID LO 1980/4-1 and Grant ID LO 1980/7-1).

## Data Availability Statement

The data that support the findings of this study are openly available in Zenodo at https://doi.org/10.5281/zenodo.6359920.

The authors made the following contributions. Maren Klingelhöfer-Jens: Conceptualization, Methodology, Software, Formal analysis, Data Curation, Writing - Original Draft, Visualization, Pre-registration of the study; Mana R. Ehlers: Conceptualization, Methodology, Formal analysis, Visualization, Writing - Original draft; Manuel Kuhn: Investigation, Software, Data Curation, Writing - Review & Editing; Vincent Keyaniyan: Methodology, Formal analysis, Writing -Review & Editing, Visualization, Pre-registration of the study; Tina B. Lonsdorf: Conceptualization, Methodology, Resources, Writing - Original draft, Supervision, Funding acquisition, Pre-registration of the study.

