## Supplementary Material for "Robust group- but limited individual-level (longitudinal) reliability and insights into cross-phases response prediction of conditioned fear"

**This Supplement belongs to an article which is a preprint and has not been peer-reviewed**

***Supplementary Table 1:*** Overview of experimental specifications and results of four previous studies reporting test-retest reliabilities in human fear conditioning research.

| | Fredrikson et al., 1993 | Zeidan et al., 2012 | Torrents-Rodas et al., 2014 | Ridderbusch et al., 2021 |
|---|---|---|---|---|
| **N/female/age** | 28/14/M = 28.5 (± 1.42) | 18/9/M = 38.0 (± 12.7) | 71/52/M = 22.4 (± 2.61) | 100/46/M = 33.1 (± 10.7) |
| **Reinforcement rate (%)** | 100 | 100 | Acquisition training: 75<br>Generalization: 50 | 60 |
| **Acquisition type** | Not reported | Not reported | Uninstructed but informed about the existence of contingencies[3] | Instructed (but not informed about the reinforcement rate) |
| **Extinction type** | Immediate | Immediate;<br>Extinction training consisted of 2 subphases separated by a 1-min rest period | None | 24h delayed; Extinction training consisted of 2 subphases (Ex1 and Ex2) |

| | | | | |
|---|---|---|---|---|
| **Additional phase(s)** | None | 24h delayed extinction recall immediately followed by renewal | Generalization (10 min. after acquisition training) | One re-acquisition trial prior to extinction training<br>Reinstatement-test (immediately after extinction training and reinstatement) |
| **CS quality** | Geometric shapes | Lamp in a room (2 colors) | 2 rings as CSs and 8 rings as GSs | Neutral faces on colored background (background color = CS type) |
| **CS duration (s)** | 8 | 12 | 8 | 6 |
| **ITI duration (s)** | 20 – 40 | 12 – 21 | 9 – 17 | 6 – 10 |
| **US type** | Auditory (110 dB white noise) | Electrotactile | Electrotactile | Electrotactile |
| **# of habituation trials CS+/CS-** | 4/4 | 4/4 | 6/6 | 2/2 |
| **# of acquisition trials CS+/CS-** | 8/8 | 5/5 | 12/12 | 10/10 |
| **# of extinction trials CS+/CS-** | 8/8 | 5/5 (in each of the 2 subphases) | No extinction phase | Extinction phase 1 (Ex1): 10/10<br>Extinction phase 2 (Ex2): 10/10 |
| **# of trials add. phase CS+/CS-** | No additional phase | Extinction recall: 5/5<br>Renewal: 5/5 | 12/12<br>6 times each GS | Re-acquisition: 1/0<br>Reinstatement-test: 10/10 |
| **SCR** | Yes | Yes | Yes | Yes |
| **FPS** | No | No | Yes | Yes |
| **Ratings** | No | No | Risk ratings | Expectancy, arousal, valence ratings |
| **fMRI** | No | No | No | Yes |
| **Reported measure(s)** | SCR | SCR | SCR, FPS, ratings | fMRI, ratings |

| | | | | |
|---|---|---|---|---|
| **# of measurement time points** | 2 | 3 | 2 | 2 |
| **Time gap between measurement time points** | 20 days | Time points 1 and 2:<br>17.9 ± 2.1 weeks<br>Time points 2 and 3:<br>14.5 ± 0.7 weeks | 5.8 - 9.0 months (M = 7.7) | 13 weeks |
| **Same stimuli used in retest** | Not reported | No[1] | Yes (half of the participants)<br>New set (other half of the participants<br>(new stimuli = lines with varying slopes) | No[5] |
| **Same allocation of stimuli to CSs** | Not reported | No[2] | Yes (applies to the use of the same stimulus set) | No[5] |
| **Reliability measure** | Pearson's r | ICC (no type specified) | G coefficient (range = 0 - 1) | ICC(1,1)[6] |
| **Included trials** | All | All | All | See results and notes below |
| **Test-Retest Habituation** | | | | |
| **CS+** | SCR (FIR): 0.62 | SCR (time points 1-3): 0.10<br>SCR (time points 1-2): 0.16 | Not reported | **fMRI**<br>No fMRI data for habituation<br>**Ratings**<br>No rating data for habituation |
| **CS-** | SCR (FIR): 0.72 | Not reported | | |
| **Test-Retest Acquisition** | | | | |
| **CS+** | SCR (FIR): 0.85<br>SCR (SIR): 0.51<br>SCR (TIR): 0.65 | SCR (time points 1-3): 0.68<br>SCR (time points 1-2): 0.64 | **Same stimulus set**[4]<br>SCR: 0.27<br>FPS: 0.34<br>Ratings: 0.23<br>**New stimulus set**[4]<br>SCR: 0.39 | **fMRI**<br>No fMRI data for acquisition training<br>**Ratings**<br>Expectancy: not reported<br>Arousal: no data for acquisition training |
| **CS-** | SCR (FIR): 0.57<br>SCR (SIR): 0.27<br>SCR (TIR): 0.29 | Not reported | | |

| | | | | |
|---|---|---|---|---|
| **CS discrimination** | Not included | SCR (time points 1-3): 0.43 | FPS: 0.40<br>Ratings: 0.46 | Valence: no data for acquisition training |
| **Test-Retest Extinction** | | | | |
| **CS+** | SCR (FIR): 0.62<br>SCR (SIR): 0.27<br>SCR (TIR): 0.83 | SCR (time points 1-3): -0.19<br>SCR (time points 1-2): -0.24 | No extinction phase | **<u>fMRI</u>**<br>**Ex1**<br>Right insula: 0.54<br>Left insula: 0.57<br>Middle cingulate cortex: 0.40<br>**Ex1 > Ex2**<br>Left insula: 0.22<br>Right insula: 0.14<br>Middle cingulate cortex: 0.29<br>**<u>Ratings</u>**<br>**pre Ex1**<br>Expectancy: no data for pre Ex1<br>Arousal: not reported<br>Valence: not reported<br>**Post Re-Acq, post Ex1 and post Ex2[7]**<br>Expectancy: 0.66<br>Arousal: 0.63<br>Valence: 0.56 |
| **CS-** | SCR (FIR): 0.37<br>SCR (SIR): -0.05<br>SCR (TIR): 0.09 | Not reported | | Not reported |
| **CS discrimination** | Not included | Not reported | | **<u>fMRI</u>**<br>**Ex1**<br>Right insula: 0.44<br>Left insula: 0.39<br>Middle cingulate cortex: 0.34<br>**Ex1 > Ex2**<br>Left insula: 0.20<br>Right insula: 0.01<br>Middle cingulate cortex: 0.13 |

| | | | | |
|---|---|---|---|---|
| | | | | <u>**Ratings**</u><br>**pre Ex1**<br>Expectancy: no data for pre Ex1<br>Arousal: 0.42<br>Valence: 0.02<br><br>**Post Re-Acq, post Ex1 and post Ex2[7]**<br>Expectancy: 0.64<br>Arousal: 0.43<br>Valence: 0.25 |
| **Test-Retest additional phase** | | **Extinction recall Renewal** | **Generalization** | **Re-Acquisition Reinstatement-Test** |
| **CS+** | No additional phase | **Extinction recall:**<br>SCR (time points 1-3): 0.46<br>SCR (time points 1-2): 0.72<br><br>**Renewal:**<br>SCR (time points 1-3): 0.67<br>SCR (time points 1-2): 0.66 | **Same stimulus set[4]**<br>SCR: 0.44<br>FPS: 0.22<br>Ratings: 0.22<br><br>**New stimulus set[4]**<br>SCR: 0.21<br>FPS: 0.16<br>Ratings: 0.25 | <u>**fMRI**</u><br>Not reported<br><br><u>**Ratings**</u><br>**post Re-Acquisition**<br>Expectancy: 0.51<br>Arousal: 0.53<br>Valence: 0.49 |
| **CS-** | | Not reported | | Not reported |
| **CS discrimination** | | **Extinction Recall**<br>SCR (time points 1-3): 0.23<br><br>**Renewal**<br>SCR (time points 1-3): 0.50 | | <u>**fMRI**</u><br>**RI-T:**<br>**Cingulate cortex cluster**<br>pre RI[8]: 0.01<br>post RI[9]: -0.05<br>pre vs. post RI[8,9]: -0.12<br><br><u>**Ratings**</u><br>**post Re-Acquisition**<br>Expectancy: 0.49<br>Arousal: not reported<br>Valence: not reported<br><br>**pre RI[10]**<br>Expectancy: 0.67 |

| | | | | |
|---|---|---|---|---|
| | | | | Arousal: 0.53<br>Valence: 0.39<br>**post RI**<br>Expectancy: 0.52<br>Arousal: 0.55<br>Valence: 0.34<br>**pre vs. post RI**[10]<br>Expectancy: 0.22<br>Arousal: 0.19<br>Valence: -0.03 |
| **Physiological response quantification** | | | | |
| **SCR quantification** | Trough-to-peak (TTP) | Baseline correction | Baseline correction | Trough-to-peak (TTP) |
| **SCR scoring criteria** | FIR: 1-4 s after CS onset<br>SIR: 5-9 s after CS onset<br>TIR: 1-4 s after CS termination | Baseline: means SCL during 2 s before trial onset subtracted from the highest SCL within the 12 s CS duration | Value at stimulus onset subtracted from the maximum value during 1-5 s after stimulus onset (only trials without risk ratings analyzed) | First response occurring 0.9-4 s after stimulus onset |
| **FPS specifications** | No FPS applied | No FPS applied | 5s after onset of odd trials and during ITIs (6 times per phase, IPIs 18-25 s) | Either 4.5 or 5 s after CS onset and during ITI (2, 3, 4, 5, or 6 s after CS offset); presented during all CS trials during habituation and during 8 of 10 CS trials during fear acquisition training |
| **FPS quantification** | | | Baseline correction | Trough-to-peak (TTP) |
| **FPS scoring criteria** | | | Value at response peak (Response onset in a time window 20-100 ms after probe onset with a peak between 20 and 150 ms after probe onset) subtracted from a baseline value (averaged during the 50 ms preceding the probe) | Response in a time window 20-120 ms after probe onset with a maximum peak within 150 ms after onset |
| **Ratings provided** | No ratings provided | No ratings provided | During even trials | Expectancy: before each CS trial |

| | | | | |
|---|---|---|---|---|
| | | | | Arousal and Valence: post Re-Acq, pre Ex1, post Ex1, post Ex2, post RI, post RIT |

*Note.* # = number; FIR = first interval response, occurring 1-4s after CS onset; SIR = second interval response, occurring 5-9s after CS onset; TIR = third interval response, occurring 1-4s after CS termination; GS = generalization stimulus; Ex1 = first extinction phase; Ex2 = second extinction phase; pre/post = prior and subsequent to respective phases; Re-Acq = re-acquisition; RI-T = reinstatement-test; RI = reinstatement.

[1] "Conditioning context and color of the CS+ were different for each of the 3 sessions and counterbalanced across visits." (Zeidan et al., 2012, p. 314)

[2] "The conditioning context and the color of the CS+ were different for each of the three test sessions and counterbalanced across visits." (Zeidan et al., 2012, p. 315)

[3] "They were not instructed about the CS−US contingency, but were told that they might learn to predict the shock if they pay attention to the presented stimuli." (Torrents-Rodas et al., 2014, p. 699)

[4] The G coefficient includes both responses to the CS+ and CS-.

[5] "The whole experimental protocol (t1) was repeated after an interval of an average of 13 weeks (second measurement: t2), using two different visual stimuli as CSs to avoid re -acquisition." (Ridderbusch et al., 2021, p. 3)

[6] One-way random effects model with single measures.

[7] Post re-acq, post Ex1 and post Ex2 = "extinction training effect" (see Ridderbusch et al., 2021).

[8] Pre RI means for fMRI: last half of Ex2 trials (5 trials).

[9] Post RI means for fMRI: first half of RI-T (5 trials).

[10] Pre RI means for ratings: post Ex2.

### Deviations from the pre-registration

***Supplementary Table 2:*** Deviations from pre-registration.

| Pre-registration | Deviation type | Manuscript | Justification |
|---|---|---|---|
| Not pre-registered | Additional analyses | Analyses of cross-sectional reliability | Considered to provide additional valuable information |
| Mixed-effects approach for calculation of ICCS | Changes analysis approach | ANOVA approach for calculation of ICCS | Statistical approach changed due to model non-convergence problems |
| Calculate ICCs for ranked and non-ranked data | Omitted pre-registered specification | Non-ranked ICCs only | During closer inspection of the conceptualization of $ICC_{con}$, we realised that it would be redundant to calculate both $ICC_{abs}$ and $ICC_{con}$ with ranked and non-ranked data as $ICC_{con}$ itself ranks the data. Hence, we decided to calculate ICCs based on non-ranked data only. |
| Not pre-registered | Additional analyses | Inclusion of ICC for SCRs to the US and US aversiveness ratings | Considered to provide valuable information |
| Not pre-registered | Additional analyses | Additional phase operationalization: last two extinction trials | Considered to provide valuable information for completeness |
| Not pre-registered | Additional analyses | Trial-by trial ICCs for SCRs | Considered to provide additional valuable information |
| Not pre-registered | Additional analyses | Inclusion of analysis focusing on reliability at the group level for SCRs | Considered to provide additional valuable information |
| Not pre-registered | Additional outcome measure | Inclusion of fMRI as an outcome measure and corresponding reliability analyses as well as within-session predictability analyses | Considered to provide valuable information |
| Multiple linear regression with SCRs or fear ratings during both acquisition and extinction training as multiple predictors for responses at reinstatement-test | Changes in analysis approach | Simple linear regressions including SCRs or fear ratings during acquisition training as predictors and responding during reinstatement-test as criterion. Further checks of statistical assumptions revealed heteroscedasticity of the data. Therefore, we conducted simple linear regressions with robust standard errors instead of using classical OLS estimators | Due to multicollinearity of the predictors resulting from significant associations of responding during acquisition and extinction training the pre-registered analyses were not suitable |
| Not pre-registered | Additional analyses | We compare different patterns of SCR, fear rating and fMRI data after pooling them for certain data specifications/ROIs in predictability analyses | Considered to provide additional valuable information |

**Cross-sectional reliability of log-transformed as well as log-transformed and range corrected SCRs**

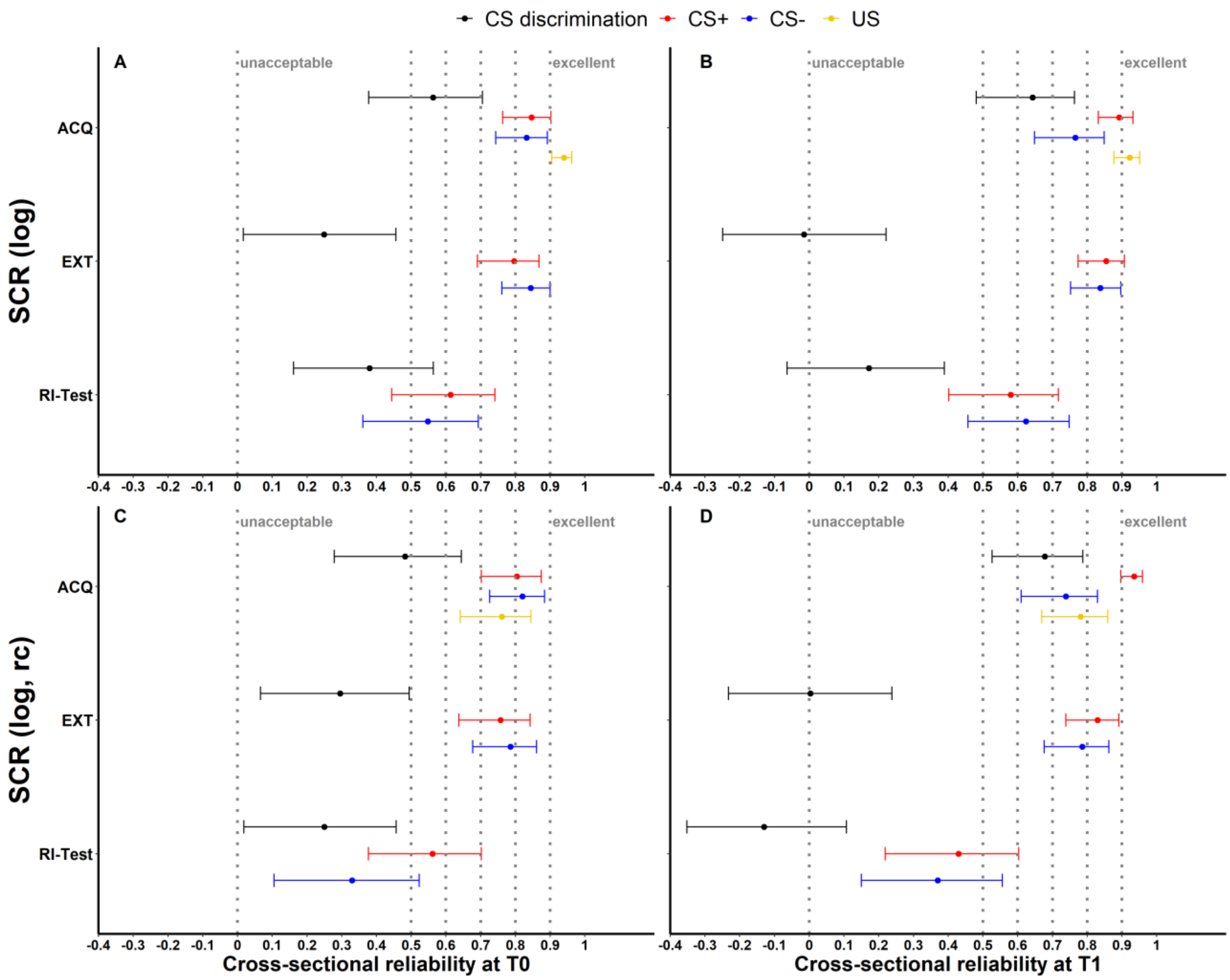


*Supplementary Figure 1.* Illustration of (A-B) cross-sectional reliability for log-transformed (log) as well as (C-D) log-transformed and range corrected (log rc) SCRs at T0 and T1 color coded for stimulus-type. Error bars represent 95% confidence intervals indicating significance, when zero is not included in the interval. The y-axis comprises the different experimental phases. Cross-sectional reliability is interpreted using benchmarks (Kline, 2013) for unacceptable (< 0.5), poor (> 0.5 but < 0.6), questionable (> 0.6 but < 0.7), acceptable (> 0.7 but < 0.8), good (> 0.8

but < 0.9) and excellent (≥ 0.9). ACQ = acquisition training, EXT = extinction training, RI-Test = reinstatement-test.

### Detailed results of ICC calculations: SCR

***Supplementary Table 3:*** *$ICC_{abs}$ and $ICC_{con}$ for all data specifications of SCRs.*

| Outcome | Ampl.-type | Stim.-type | Phase | Op. | $ICC_{abs}$ | | | | $ICC_{con}$ | | | |
|---|---|---|---|---|---|---|---|---|---|---|---|---|
| | | | | | Value | Lower 95% CI | Upper 95% CI | p-value | Value | Lower 95% CI | Upper 95% CI | p-value |
| SCR | raw | CS dis. | Acq | average | 0.160 | -0.020 | 0.340 | .077 | 0.170 | -0.030 | 0.350 | .077 |
| SCR | raw | CS+ | Acq | average | 0.270 | 0.090 | 0.440 | .006 | 0.300 | 0.110 | 0.460 | .006 |
| SCR | raw | CS- | Acq | average | 0.390 | 0.210 | 0.540 | .000 | 0.410 | 0.230 | 0.560 | .000 |
| SCR | raw | US | Acq | average | 0.317 | 0.133 | 0.481 | .003 | 0.322 | 0.135 | 0.487 | .003 |
| SCR | raw | CS dis. | Acq | last 2 trials | 0.240 | 0.060 | 0.420 | .017 | 0.250 | 0.060 | 0.430 | .017 |
| SCR | raw | CS+ | Acq | last 2 trials | 0.220 | 0.040 | 0.390 | .025 | 0.230 | 0.040 | 0.410 | .025 |
| SCR | raw | CS- | Acq | last 2 trials | 0.190 | 0.000 | 0.370 | .055 | 0.190 | -0.010 | 0.370 | .055 |
| SCR | raw | CS dis. | Ext | 1st trial | 0.190 | 0.010 | 0.360 | .044 | 0.200 | 0.010 | 0.380 | .044 |
| SCR | raw | CS+ | Ext | 1st trial | 0.320 | 0.140 | 0.490 | .003 | 0.330 | 0.140 | 0.490 | .003 |
| SCR | raw | CS- | Ext | 1st trial | 0.040 | -0.120 | 0.210 | .331 | 0.050 | -0.140 | 0.250 | .331 |
| SCR | raw | CS dis. | Ext | average | 0.070 | -0.130 | 0.270 | .272 | 0.070 | -0.120 | 0.260 | .272 |
| SCR | raw | CS+ | Ext | average | 0.480 | 0.320 | 0.620 | .000 | 0.500 | 0.340 | 0.630 | .000 |
| SCR | raw | CS- | Ext | average | 0.580 | 0.430 | 0.700 | .000 | 0.610 | 0.470 | 0.720 | .000 |
| SCR | raw | CS dis. | Ext | last 2 trials | 0.060 | -0.140 | 0.250 | .321 | 0.060 | -0.140 | 0.250 | .321 |
| SCR | raw | CS+ | Ext | last 2 trials | 0.170 | -0.020 | 0.360 | .073 | 0.170 | -0.020 | 0.360 | .073 |
| SCR | raw | CS- | Ext | last 2 trials | 0.200 | 0.000 | 0.380 | .048 | 0.200 | 0.000 | 0.380 | .048 |
| SCR | raw | CS dis. | RI-T | 1st trial | 0.140 | -0.050 | 0.320 | .119 | 0.140 | -0.060 | 0.330 | .119 |
| SCR | raw | CS+ | RI-T | 1st trial | 0.150 | -0.040 | 0.330 | .098 | 0.150 | -0.040 | 0.340 | .098 |
| SCR | raw | CS- | RI-T | 1st trial | 0.030 | -0.130 | 0.200 | .389 | 0.030 | -0.160 | 0.230 | .389 |
| SCR | raw | US | RI | average | 0.271 | 0.085 | 0.440 | .009 | 0.278 | 0.087 | 0.449 | .009 |

| Outcome | Ampl.-type | Stim.-type | Phase | Op. | $ICC_{abs}$ | | | | $ICC_{con}$ | | | |
|---|---|---|---|---|---|---|---|---|---|---|---|---|
| | | | | | Value | Lower 95% CI | Upper 95% CI | p-value | Value | Lower 95% CI | Upper 95% CI | p-value |
| SCR | log | CS dis. | Acq | average | 0.180 | 0.000 | 0.350 | .054 | 0.190 | 0.000 | 0.370 | .054 |
| SCR | log | CS+ | Acq | average | 0.290 | 0.100 | 0.450 | .004 | 0.310 | 0.130 | 0.480 | .004 |
| SCR | log | CS- | Acq | average | 0.400 | 0.230 | 0.550 | .000 | 0.420 | 0.250 | 0.570 | .000 |
| SCR | log | US | Acq | average | 0.320 | 0.137 | 0.482 | .003 | 0.327 | 0.140 | 0.491 | .003 |
| SCR | log | CS dis. | Acq | last 2 trials | 0.230 | 0.040 | 0.400 | .024 | 0.230 | 0.040 | 0.410 | .024 |
| SCR | log | CS+ | Acq | last 2 trials | 0.210 | 0.020 | 0.380 | .032 | 0.220 | 0.030 | 0.400 | .032 |
| SCR | log | CS- | Acq | last 2 trials | 0.190 | 0.000 | 0.370 | .052 | 0.190 | 0.000 | 0.370 | .052 |
| SCR | log | CS dis. | Ext | 1st trial | 0.180 | 0.000 | 0.350 | .052 | 0.190 | 0.000 | 0.370 | .052 |
| SCR | log | CS+ | Ext | 1st trial | 0.310 | 0.120 | 0.470 | .004 | 0.310 | 0.120 | 0.480 | .004 |
| SCR | log | CS- | Ext | 1st trial | 0.030 | -0.130 | 0.200 | .368 | 0.040 | -0.160 | 0.230 | .368 |
| SCR | log | CS dis. | Ext | average | 0.060 | -0.140 | 0.260 | .301 | 0.060 | -0.130 | 0.250 | .301 |
| SCR | log | CS+ | Ext | average | 0.490 | 0.320 | 0.620 | .000 | 0.500 | 0.340 | 0.640 | .000 |
| SCR | log | CS- | Ext | average | 0.590 | 0.430 | 0.710 | .000 | 0.620 | 0.480 | 0.720 | .000 |
| SCR | log | CS dis. | Ext | last 2 trials | 0.070 | -0.130 | 0.260 | .283 | 0.070 | -0.130 | 0.260 | .283 |
| SCR | log | CS+ | Ext | last 2 trials | 0.200 | 0.010 | 0.380 | .044 | 0.200 | 0.010 | 0.380 | .044 |
| SCR | log | CS- | Ext | last 2 trials | 0.230 | 0.040 | 0.410 | .027 | 0.230 | 0.030 | 0.410 | .027 |
| SCR | log | CS dis. | RI-T | 1st trial | 0.170 | -0.020 | 0.350 | .076 | 0.170 | -0.030 | 0.350 | .076 |
| SCR | log | CS+ | RI-T | 1st trial | 0.160 | -0.030 | 0.340 | .083 | 0.160 | -0.030 | 0.350 | .083 |
| SCR | log | CS- | RI-T | 1st trial | 0.040 | -0.120 | 0.210 | .337 | 0.050 | -0.150 | 0.240 | .337 |
| SCR | log | US | RI | average | 0.299 | 0.116 | 0.464 | .004 | 0.308 | 0.120 | 0.475 | .004 |
| SCR | log rc | CS dis. | Acq | average | 0.230 | 0.050 | 0.410 | .018 | 0.250 | 0.050 | 0.420 | .018 |
| SCR | log rc | CS+ | Acq | average | 0.490 | 0.310 | 0.630 | .000 | 0.530 | 0.370 | 0.660 | .000 |
| SCR | log rc | CS- | Acq | average | 0.610 | 0.470 | 0.730 | .000 | 0.630 | 0.500 | 0.740 | .000 |
| SCR | log rc | US | Acq | average | 0.112 | -0.086 | 0.301 | .176 | 0.111 | -0.086 | 0.300 | .176 |
| SCR | log rc | CS dis. | Acq | last 2 trials | 0.050 | -0.150 | 0.240 | .337 | 0.050 | -0.150 | 0.240 | .337 |
| SCR | log rc | CS+ | Acq | last 2 trials | 0.300 | 0.120 | 0.470 | .004 | 0.310 | 0.120 | 0.480 | .004 |

| Outcome | Ampl.-type | Stim.-type | Phase | Op. | $ICC_{abs}$ | | | | $ICC_{con}$ | | | |
|---|---|---|---|---|---|---|---|---|---|---|---|---|
| | | | | | Value | Lower 95% CI | Upper 95% CI | p-value | Value | Lower 95% CI | Upper 95% CI | p-value |
| SCR | log rc | CS- | Acq | last 2 trials | 0.190 | -0.010 | 0.370 | .056 | 0.190 | -0.010 | 0.370 | .056 |
| SCR | log rc | CS dis. | Ext | 1st trial | 0.200 | 0.020 | 0.370 | .033 | 0.220 | 0.020 | 0.400 | .033 |
| SCR | log rc | CS+ | Ext | 1st trial | 0.270 | 0.080 | 0.440 | .012 | 0.270 | 0.070 | 0.440 | .012 |
| SCR | log rc | CS- | Ext | 1st trial | -0.090 | -0.250 | 0.090 | .804 | -0.100 | -0.290 | 0.090 | .804 |
| SCR | log rc | CS dis. | Ext | average | 0.350 | 0.160 | 0.510 | .002 | 0.340 | 0.160 | 0.510 | .002 |
| SCR | log rc | CS+ | Ext | average | 0.540 | 0.380 | 0.670 | .000 | 0.560 | 0.410 | 0.680 | .000 |
| SCR | log rc | CS- | Ext | average | 0.620 | 0.440 | 0.740 | .000 | 0.660 | 0.530 | 0.760 | .000 |
| SCR | log rc | CS dis. | Ext | last 2 trials | 0.210 | 0.020 | 0.390 | .037 | 0.210 | 0.020 | 0.390 | .037 |
| SCR | log rc | CS+ | Ext | last 2 trials | 0.360 | 0.170 | 0.520 | .001 | 0.350 | 0.170 | 0.510 | .001 |
| SCR | log rc | CS- | Ext | last 2 trials | 0.440 | 0.270 | 0.590 | .000 | 0.440 | 0.270 | 0.590 | .000 |
| SCR | log rc | CS dis. | RI-T | 1st trial | 0.230 | 0.040 | 0.410 | .023 | 0.240 | 0.040 | 0.410 | .023 |
| SCR | log rc | CS+ | RI-T | 1st trial | 0.170 | -0.020 | 0.360 | .071 | 0.170 | -0.020 | 0.360 | .071 |
| SCR | log rc | CS- | RI-T | 1st trial | 0.150 | -0.030 | 0.330 | .086 | 0.160 | -0.030 | 0.350 | .086 |
| SCR | log rc | US | RI | average | 0.093 | -0.106 | 0.284 | .221 | 0.092 | -0.105 | 0.282 | .221 |

Note. Ampl. = Amplitude, Stim. = Stimulus, Op. = Operationalization, CI = Confidence Interval, CS dis. = CS discrimination, log = log-transformed, log rc = log-transformed and range corrected, Acq = Acquisition training, Ext = Extinction training, RI = Reinstatement, RI-T = Reinstatement-Test.

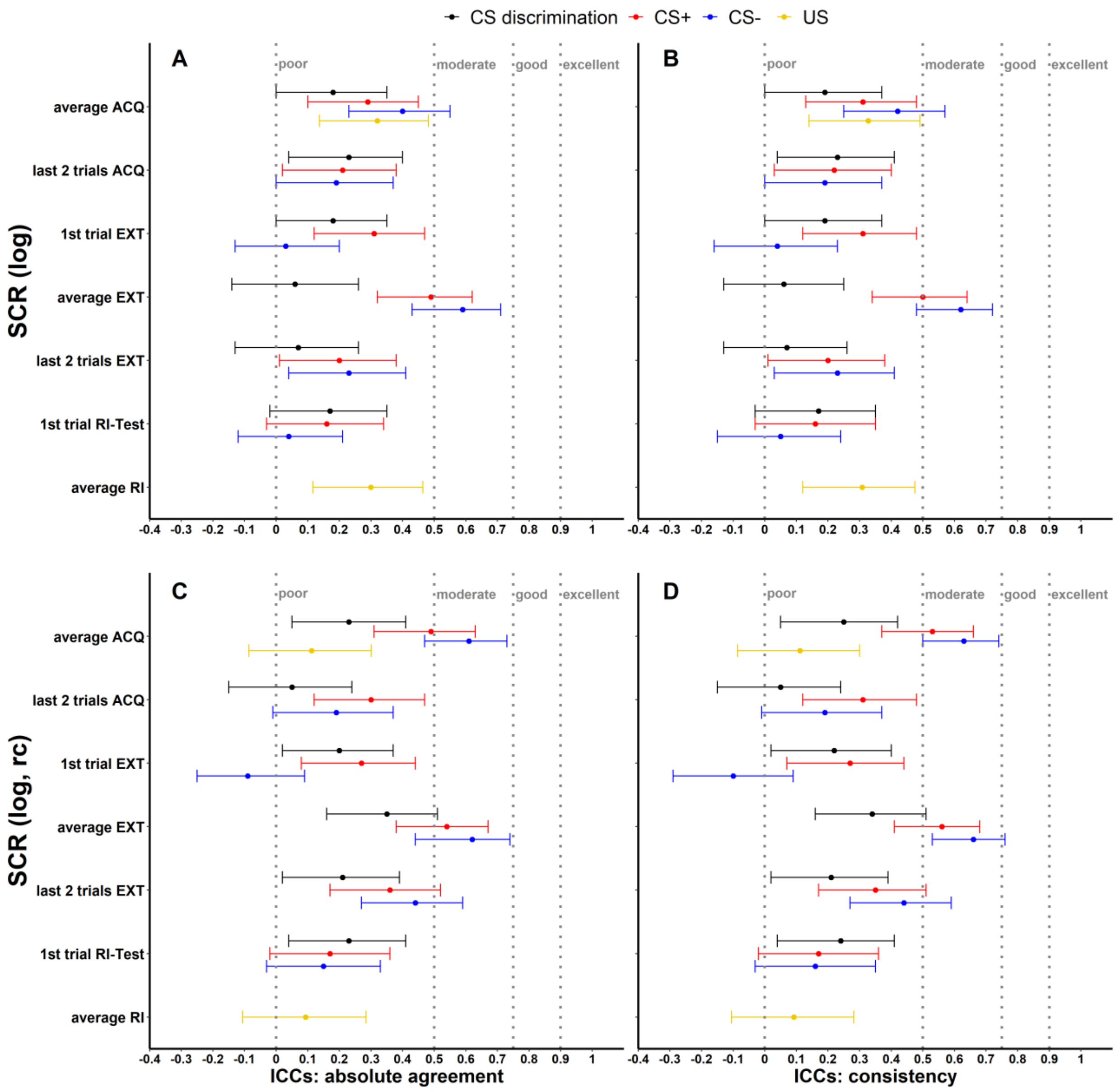


*Supplementary Figure 2.* Illustration of (A-B) ICCs of log-transformed (log) as well as (C-D) log-transformed and range corrected (log, rc) SCRs color coded for stimulus-type. The y-axis comprises the different phase operationalizations. A and C display $ICC_{abs}$, B and D display $ICC_{con}$. ICCs < 0.5, < 0.75, < 0.9 and > 0.9 were interpreted as poor, moderate, good and excellent respectively (Koo & Li, 2016). Error bars represent 95% confidence intervals indicating significance of ICCs, when zero is not included in the interval. ACQ = acquisition training, EXT = extinction training, RI = reinstatement, RI-Test = reinstatement-test.

### ICCs of trial-by-trial SCRs

#### ICCs of trial-by-trial raw SCRs

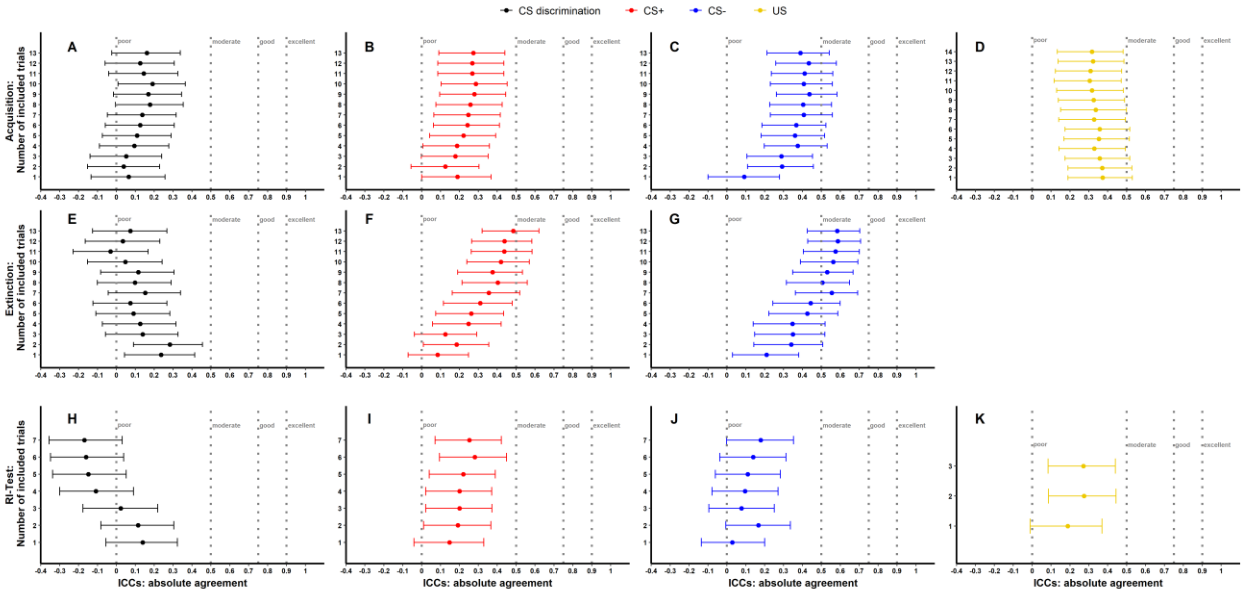


*Supplementary Figure 3.* Illustration of $ICC_{abs}$ of trial-by-trial raw SCRs for phases (A-D: Acquisition, E-G: Extinction, H-J: Reinstatement-Test, K: Reinstatement) and stimulus-types separately. Trials were averaged starting with the first (i.e., reinstatement-test and US trials) or second trial (i.e., acquisition and extinction training), adding all preceding trials trial-by-trial and averaged. ICCs < 0.5, < 0.75, < 0.9 and > 0.9 (Koo & Li, 2016) were interpreted as poor, moderate, good and excellent respectively. Error bars represent 95% confidence intervals. Non-overlapping error bars indicate significant differences between ICCs within one figure. RI-Test = reinstatement-test.

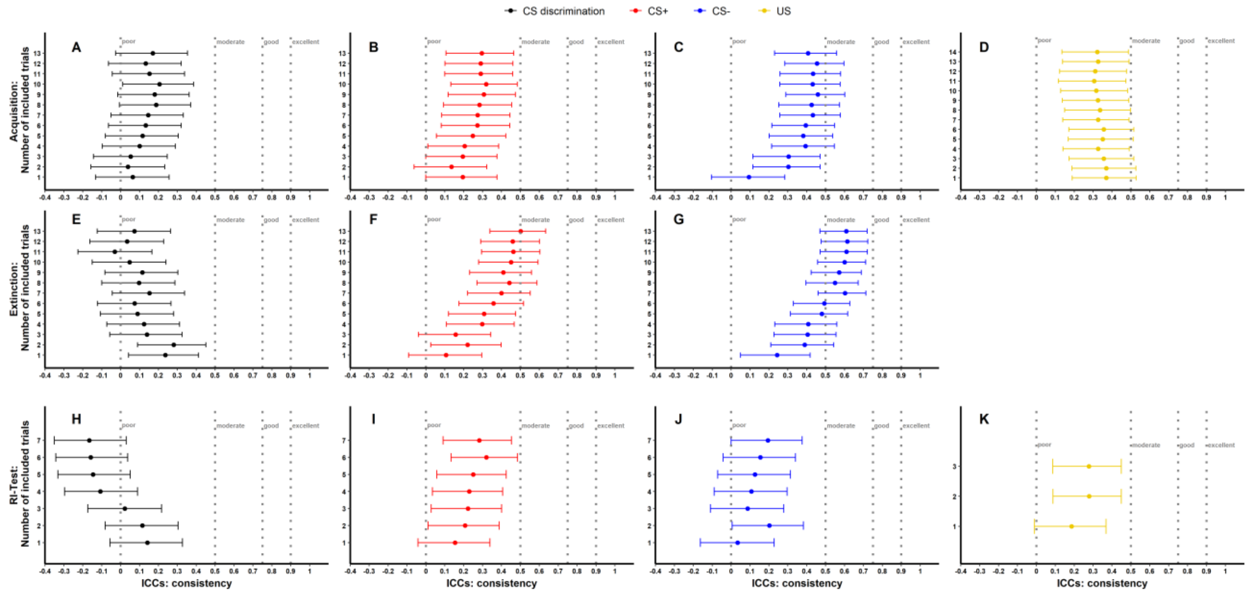


*Supplementary Figure 4.* Illustration of $ICC_{con}$ of trial-by-trial raw SCRs for phases (A-D: Acquisition, E-G: Extinction, H-J: Reinstatement-Test, K: Reinstatement) and stimulus-types separately. Trials were averaged starting with the first (i.e., reinstatement-test and US trials) or second trial (i.e., acquisition and extinction training), adding all preceding trials trial-by-trial and averaged. ICCs < 0.5, < 0.75, < 0.9 and > 0.9 (Koo & Li, 2016) were interpreted as poor, moderate, good and excellent respectively. Error bars represent 95% confidence intervals. Non-overlapping error bars indicate significant differences between ICCs within one figure. RI-Test = reinstatement-test.

**ICCs of trial-by-trial log-transformed SCRs**

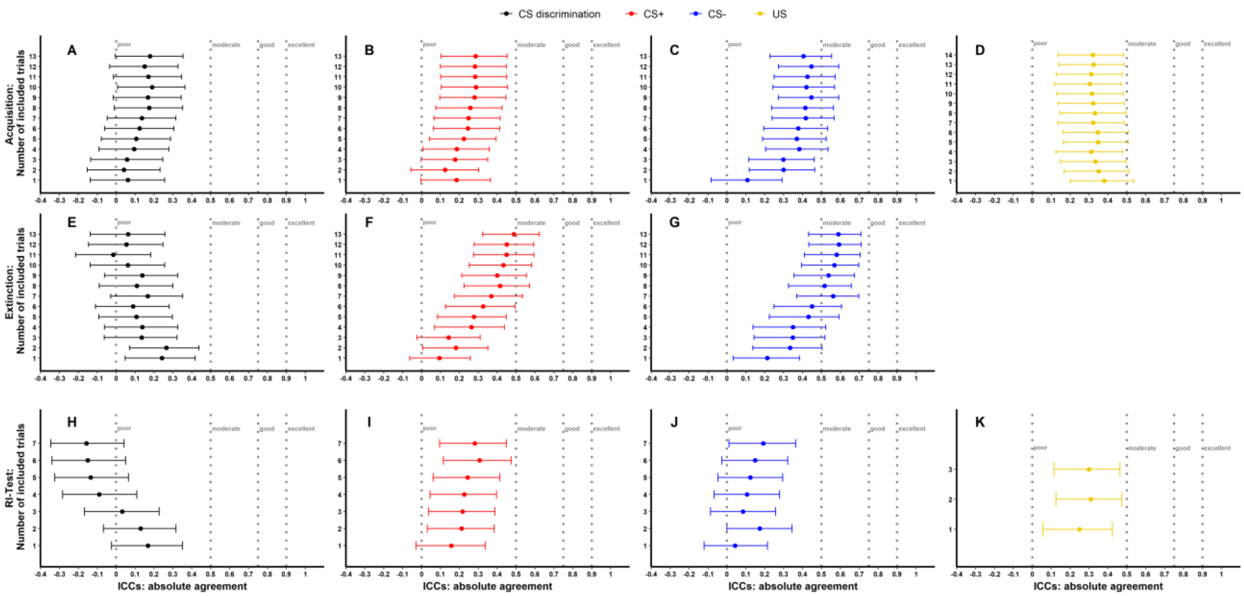


*Supplementary Figure 5.* Illustration of $ICC_{abs}$ of trial-by-trial log-transformed SCRs for phases (A-D: Acquisition, E-G: Extinction, H-J: Reinstatement-Test, K: Reinstatement) and stimulus-types separately. Trials were averaged starting with the first (i.e., reinstatement-test and US trials) or second trial (i.e., acquisition and extinction training), adding all preceding trials trial-by-trial and averaged. ICCs < 0.5, < 0.75, < 0.9 and > 0.9 (Koo & Li, 2016) were interpreted as poor, moderate, good and excellent respectively. Error bars represent 95% confidence intervals. Non-overlapping error bars indicate significant differences between ICCs within one figure. RI-Test = reinstatement-test.

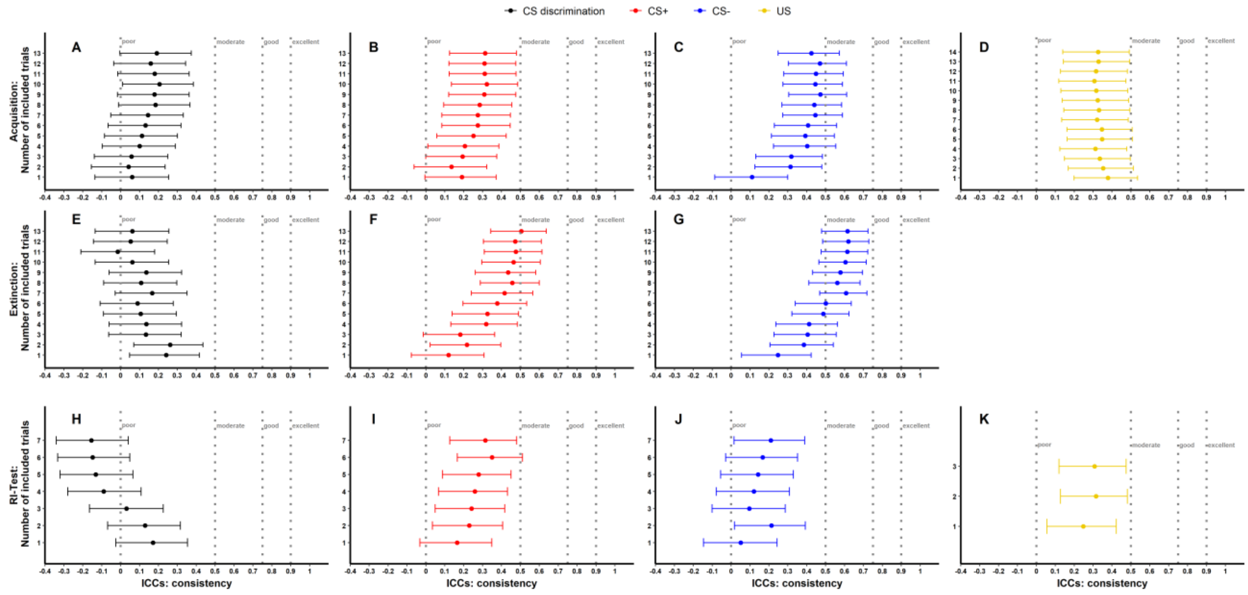


*Supplementary Figure 6.* Illustration of $ICC_{con}$ of trial-by-trial log-transformed SCRs for phases (A-D: Acquisition, E-G: Extinction, H-J: Reinstatement-Test, K: Reinstatement) and stimulus-types separately. Trials were averaged starting with the first (i.e., reinstatement-test and US trials) or second trial (i.e., acquisition and extinction training), adding all preceding trials trial-by-trial and averaged. ICCs < 0.5, < 0.75, < 0.9 and > 0.9 (Koo & Li, 2016) were interpreted as poor, moderate, good and excellent respectively. Error bars represent 95% confidence intervals. Non-overlapping error bars indicate significant differences between ICCs within one figure. RI-Test = reinstatement-test.

**ICCs of trial-by-trial log-transformed and range corrected SCRs**

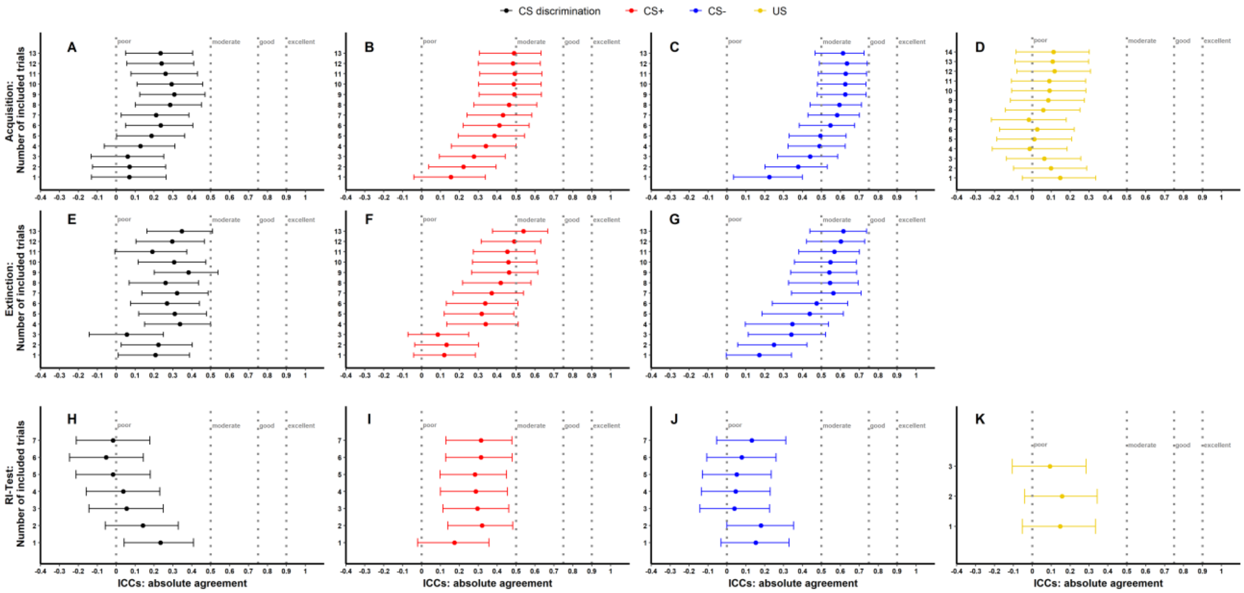


*Supplementary Figure 7.* Illustration of $ICC_{abs}$ of trial-by-trial log-transformed and range corrected SCRs for phases (A-D: Acquisition, E-G: Extinction, H-J: Reinstatement-Test, K: Reinstatement) and stimulus-types separately. Trials were averaged starting with the first (i.e., reinstatement-test and US trials) or second trial (i.e., acquisition and extinction training), adding all preceding trials trial-by-trial and averaged. ICCs < 0.5, < 0.75, < 0.9 and > 0.9 (Koo & Li, 2016) were interpreted as poor, moderate, good and excellent respectively. Error bars represent 95% confidence intervals. Non-overlapping error bars indicate significant differences between ICCs within one figure. RI-Test = reinstatement-test.

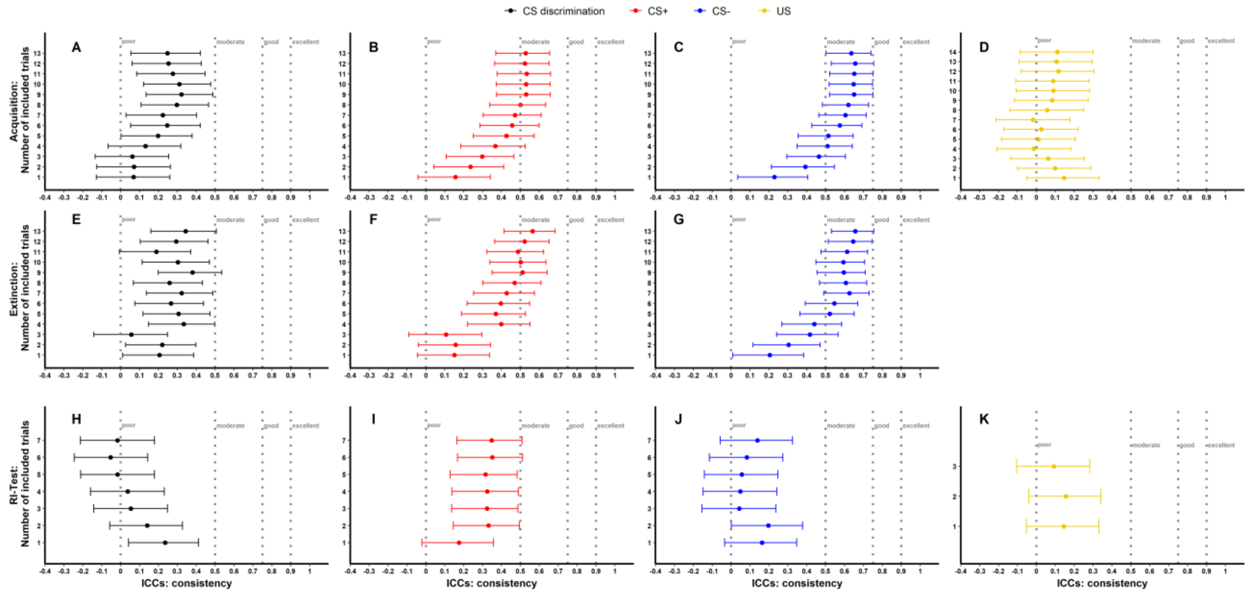


*Supplementary Figure 8.* Illustration of $ICC_{con}$ of trial-by-trial log-transformed and range corrected SCRs for phases (A-D: Acquisition, E-G: Extinction, H-J: Reinstatement-Test, K: Reinstatement) and stimulus-types separately. Trials were averaged starting with the first (i.e., reinstatement-test and US trials) or second trial (i.e., acquisition and extinction training), adding all preceding trials trial-by-trial and averaged. ICCs < 0.5, < 0.75, < 0.9 and > 0.9 (Koo & Li, 2016) were interpreted as poor, moderate, good and excellent respectively. Error bars represent 95% confidence intervals. Non-overlapping error bars indicate significant differences between ICCs within one figure. RI-Test = reinstatement-test.

### Detailed results of ICC calculations: fear ratings

***Supplementary Table 4:*** *$ICC_{abs}$ and $ICC_{con}$ for all data specifications of fear ratings.*

| Outcome | Stim.-type | Phase | Op. | $ICC_{abs}$ | | | | $ICC_{con}$ | | | |
|---|---|---|---|---|---|---|---|---|---|---|---|
| | | | | Value | Lower 95% CI | Upper 95% CI | p-value | Value | Lower 95% CI | Upper 95% CI | p-value |
| Ratings | CS dis. | Acq | post-pre | 0.190 | 0.007 | 0.364 | .043 | 0.203 | 0.008 | 0.384 | .043 |
| Ratings | CS+ | Acq | post-pre | 0.436 | 0.262 | 0.582 | < .001 | 0.433 | 0.260 | 0.579 | < .001 |
| Ratings | CS- | Acq | post-pre | -0.162 | -0.328 | 0.018 | .945 | -0.190 | -0.372 | 0.005 | .945 |
| Ratings | CS dis. | Acq | post | 0.424 | 0.230 | 0.581 | < .001 | 0.470 | 0.302 | 0.609 | < .001 |
| Ratings | CS+ | Acq | post | 0.343 | 0.163 | 0.502 | .001 | 0.362 | 0.179 | 0.521 | .001 |
| Ratings | CS- | Acq | post | 0.228 | 0.045 | 0.400 | .020 | 0.242 | 0.049 | 0.417 | .020 |
| Ratings | US | Acq | post | 0.310 | 0.120 | 0.470 | .005 | 0.300 | 0.110 | 0.470 | .005 |
| Ratings | CS dis. | Ext | pre | 0.459 | 0.250 | 0.617 | < .001 | 0.516 | 0.357 | 0.646 | < .001 |
| Ratings | CS+ | Ext | pre | 0.485 | 0.266 | 0.643 | < .001 | 0.548 | 0.395 | 0.671 | < .001 |
| Ratings | CS- | Ext | pre | 0.702 | 0.587 | 0.789 | < .001 | 0.700 | 0.585 | 0.788 | < .001 |
| Ratings | CS dis. | Ext | pre-post | 0.482 | 0.308 | 0.623 | < .001 | 0.512 | 0.352 | 0.643 | < .001 |
| Ratings | CS+ | Ext | pre-post | 0.494 | 0.282 | 0.648 | < .001 | 0.552 | 0.400 | 0.675 | < .001 |
| Ratings | CS- | Ext | pre-post | 0.188 | 0.003 | 0.363 | .048 | 0.198 | 0.002 | 0.378 | .048 |
| Ratings | CS dis. | Ext | post | 0.165 | -0.025 | 0.345 | .078 | 0.169 | -0.027 | 0.353 | .078 |
| Ratings | CS+ | Ext | post | 0.474 | 0.307 | 0.613 | < .001 | 0.472 | 0.305 | 0.611 | < .001 |
| Ratings | CS- | Ext | post | 0.686 | 0.563 | 0.779 | < .001 | 0.700 | 0.584 | 0.787 | < .001 |
| Ratings | US | RI | post | 0.430 | 0.260 | 0.580 | < .001 | 0.450 | 0.280 | 0.590 | < .001 |
| Ratings | CS dis. | RI-T | pre | 0.172 | -0.021 | 0.354 | .072 | 0.174 | -0.022 | 0.357 | .072 |
| Ratings | CS+ | RI-T | pre | 0.437 | 0.264 | 0.582 | < .001 | 0.436 | 0.263 | 0.582 | < .001 |
| Ratings | CS- | RI-T | pre | 0.538 | 0.382 | 0.663 | < .001 | 0.536 | 0.381 | 0.662 | < .001 |

Note. Stim. = Stimulus, Op. = Operationalization, CI = Confidence Interval, CS dis. = CS discrimination, Acq = Acquisition training, Ext = Extinction training, RI = Reinstatement, RI-T = Reinstatement-Test, pre = prior to the experimental phase, post = subsequent to the experimental phase.

### Detailed results of ICC and similarity calculations: BOLD fMRI

***Supplementary Table 5:*** *$ICC_{abs}$ and $ICC_{con}$ for CS discrimination during fear acquisition (Acq) and extinction training (Ext).*

| Phase | ICC-type | Whole Brain | Insula | Amygdala | Hippocampus | dmPFC | vmPFC |
|---|---|---|---|---|---|---|---|
| **Acq** | ICCabs | 0.175 | 0.001 | < 0.001 | < 0.001 | 0.001 | 0.001 |
| | ICCcon | 0.175 | 0.001 | < 0.001 | < 0.001 | 0.001 | 0.001 |
| **Ext** | ICCabs | 0.008 | < 0.001 | < 0.001 | < 0.001 | < 0.001 | < 0.001 |
| | ICCcon | 0.008 | < 0.001 | < 0.001 | < 0.001 | < 0.001 | < 0.001 |

*Note.* dmPFC = dorsomedial prefrontal cortex; vmPFC = ventromedial prefrontal cortex.

***Supplementary Table 6:*** *Paired sample t-tests comparing between- and within-subject similarity for whole brain activation pattern as well as activation pattern in the ROIs for acquisition training (Acq) and extinction training (Ext).*

| Phase | ROI | *t* | *df* | *p* | *Cohen's d* |
|---|---|---|---|---|---|
| **Acq** | Whole Brain | 4.09 | 70 | **< .001** | 0.49 |
| | Insula | 4.33 | 70 | **< .001** | 0.51 |
| | Amygdala | 2.01 | 70 | **.048** | 0.24 |
| | Hippocampus | 2.18 | 70 | **.033** | 0.26 |
| | dmPFC | 2.97 | 70 | **.004** | 0.35 |
| | vmPFC | 2.39 | 70 | **.019** | 0.28 |
| **Ext** | Whole Brain | 0.33 | 70 | .740 | 0.04 |
| | Insula | 0.86 | 70 | .394 | 0.10 |
| | Amygdala | 1.26 | 70 | .211 | 0.15 |
| | Hippocampus | -0.35 | 70 | .726 | -0.04 |
| | dmPFC | -0.35 | 70 | .726 | -0.04 |
| | vmPFC | -0.06 | 70 | .955 | -0.01 |

*Note.* dmPFC = dorsomedial prefrontal cortex; vmPFC = ventromedial prefrontal cortex.

**Longitudinal reliability at the group-level: log-transformed as well as log-transformed and range corrected SCRs**

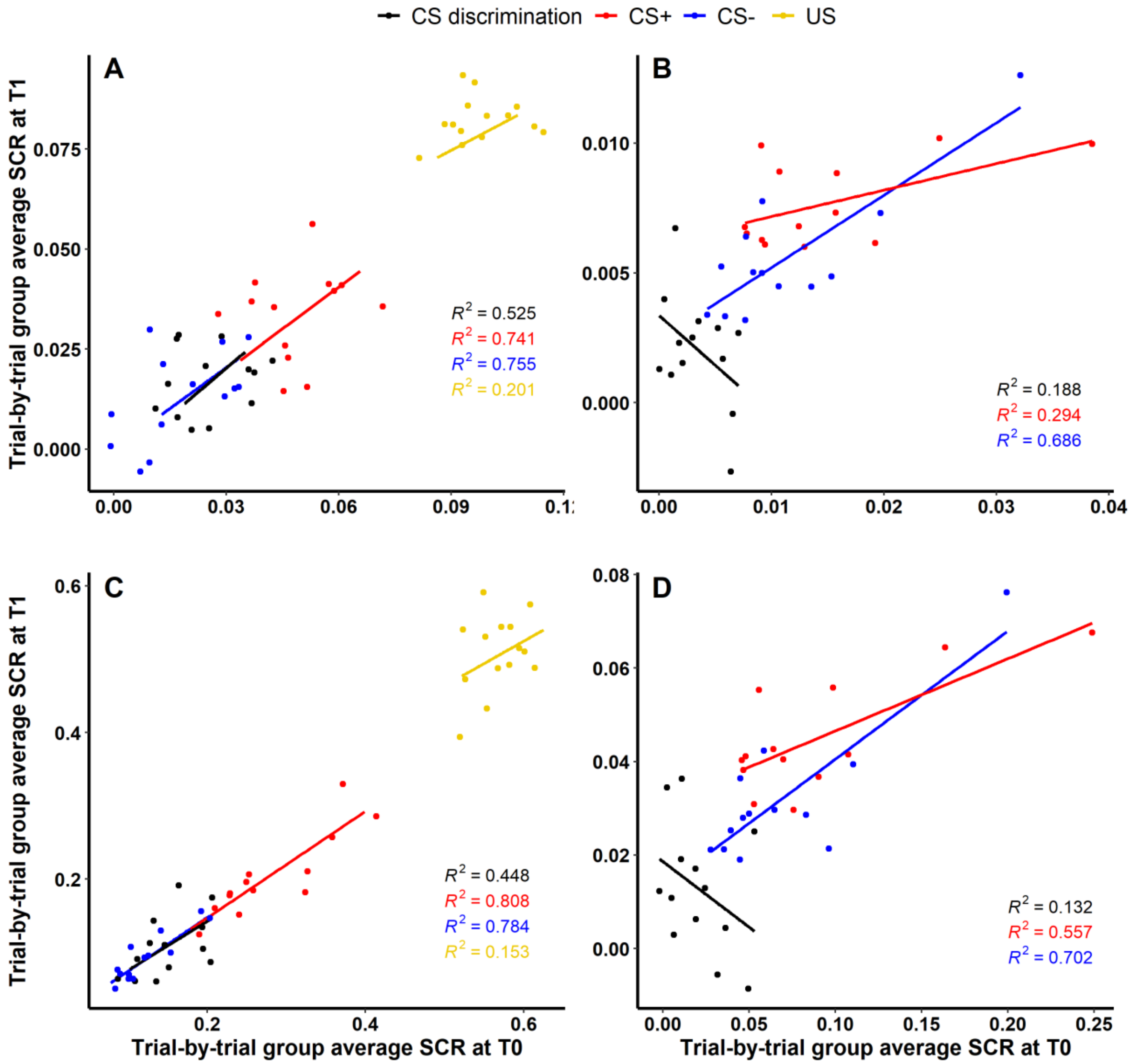


*Supplementary Figure 9.* Scatter plots illustrating longitudinal reliability at the group level during (A,C) acquisition and (B,D) extinction training for log-transformed (A,B) as well as log-transformed and range corrected (C,D) SCRs. Longitudinal reliability at the group level refers to the extent of explained variance in linear regressions comprising SCRs at T0 as independent and

SCRs at T1 as dependent variable. Results are shown for trial-by-trial group average SCRs to the CS+ (red), CS- (blue), the US (yellow) and CS discrimination (black). Single data points represent pairs of single trials at T0 and T1 averaged across participants. Note that no US was presented during extinction training and hence, no reliability of the US is shown in (B) and (D).

### Detailed results of predictability analysis: SCR and fear ratings

Cohen's $f^2$ (formula: $f^2 = R^2/1 - R^2$) was calculated as effect size. According to the guidelines of Cohen (1988), $f^2 \geq .02$, $f^2 \geq .15$ and $f^2 \geq .34$ represent small, medium and large effect sizes respectively. Since Cohen's $f^2$ is informative, but less common (Selya, Rose, Dierker, Hedeker, & Mermelstein, 2012), additionally R squared is reported as effect size.

***Supplementary Table 7:*** *Detailed results of linear regressions: SCR.*

| Outcome | Stim.-type | Ampl.-type | Ranking | Predictor | Criterion | *b* | $SE_b$ | Lower 95% CI | Upper 95% CI | *t* | *df* | *p* | $R^2$ | Cohen's $f^2$ |
|---|---|---|---|---|---|---|---|---|---|---|---|---|---|---|
| SCR | CS dis. | raw | not ranked | AVE ACQ | 1st trial EXT | 0.329 | 0.129 | 0.076 | 0.582 | 2.543 | 105 | 0.012 | 0.038 | 0.040 |
| SCR | CS dis. | raw | not ranked | AVE last 2 trials ACQ | 1st trial EXT | 0.264 | 0.080 | 0.107 | 0.421 | 3.288 | 105 | 0.001 | 0.066 | 0.071 |
| SCR | CS dis. | raw | not ranked | AVE ACQ | AVE EXT | 0.109 | 0.062 | -0.013 | 0.231 | 1.762 | 105 | 0.081 | 0.050 | 0.052 |
| SCR | CS dis. | raw | not ranked | AVE last 2 trials ACQ | AVE EXT | 0.031 | 0.031 | -0.030 | 0.092 | 0.986 | 105 | 0.327 | 0.011 | 0.011 |
| SCR | CS dis. | raw | not ranked | AVE ACQ | AVE last 2 trials EXT | 0.081 | 0.115 | -0.144 | 0.306 | 0.705 | 105 | 0.483 | 0.007 | 0.007 |
| SCR | CS dis. | raw | not ranked | AVE last 2 trials ACQ | AVE last 2 trials EXT | -0.039 | 0.105 | -0.245 | 0.167 | -0.371 | 105 | 0.711 | 0.005 | 0.005 |
| SCR | CS dis. | raw | not ranked | AVE ACQ | 1st trial RI-Test | 0.195 | 0.276 | -0.346 | 0.736 | 0.708 | 105 | 0.480 | 0.008 | 0.008 |
| SCR | CS dis. | raw | not ranked | AVE last 2 trials ACQ | 1st trial RI-Test | 0.218 | 0.230 | -0.233 | 0.669 | 0.945 | 105 | 0.347 | 0.028 | 0.029 |
| SCR | CS dis. | raw | not ranked | 1st trial EXT | 1st trial RI-Test | 0.038 | 0.165 | -0.285 | 0.361 | 0.231 | 105 | 0.817 | 0.001 | 0.001 |
| SCR | CS dis. | raw | not ranked | AVE EXT | 1st trial RI-Test | 0.222 | 0.501 | -0.760 | 1.204 | 0.443 | 105 | 0.659 | 0.003 | 0.003 |
| SCR | CS dis. | raw | not ranked | AVE last 2 trials EXT | 1st trial RI-Test | -0.316 | 0.824 | -1.931 | 1.299 | -0.384 | 105 | 0.702 | 0.020 | 0.020 |
| SCR | CS+ | raw | not ranked | AVE ACQ | 1st trial EXT | 0.686 | 0.128 | 0.435 | 0.937 | 5.347 | 105 | 0.000 | 0.291 | 0.410 |

| Outcome | Stim.-type | Ampl.-type | Ranking | Predictor | Criterion | *b* | $SE_b$ | Lower 95% CI | Upper 95% CI | *t* | *df* | *p* | $R^2$ | Cohen's $f^2$ |
|---|---|---|---|---|---|---|---|---|---|---|---|---|---|---|
| SCR | CS+ | raw | not ranked | AVE last 2 trials ACQ | 1st trial EXT | 0.508 | 0.130 | 0.253 | 0.763 | 3.909 | 105 | 0.000 | 0.212 | 0.269 |
| SCR | CS+ | raw | not ranked | AVE ACQ | AVE EXT | 0.283 | 0.076 | 0.134 | 0.432 | 3.705 | 105 | 0.000 | 0.273 | 0.375 |
| SCR | CS+ | raw | not ranked | AVE last 2 trials ACQ | AVE EXT | 0.216 | 0.077 | 0.065 | 0.367 | 2.827 | 105 | 0.006 | 0.212 | 0.270 |
| SCR | CS+ | raw | not ranked | AVE ACQ | AVE last 2 trials EXT | 0.200 | 0.099 | 0.006 | 0.394 | 2.006 | 105 | 0.047 | 0.120 | 0.137 |
| SCR | CS+ | raw | not ranked | AVE last 2 trials ACQ | AVE last 2 trials EXT | 0.143 | 0.092 | -0.037 | 0.323 | 1.550 | 105 | 0.124 | 0.082 | 0.089 |
| SCR | CS+ | raw | not ranked | AVE ACQ | 1st trial RI-Test | 0.676 | 0.132 | 0.417 | 0.935 | 5.099 | 105 | 0.000 | 0.213 | 0.270 |
| SCR | CS+ | raw | not ranked | AVE last 2 trials ACQ | 1st trial RI-Test | 0.434 | 0.147 | 0.146 | 0.722 | 2.956 | 105 | 0.004 | 0.117 | 0.132 |
| SCR | CS+ | raw | not ranked | 1st trial EXT | 1st trial RI-Test | 0.608 | 0.101 | 0.410 | 0.806 | 5.993 | 105 | 0.000 | 0.279 | 0.386 |
| SCR | CS+ | raw | not ranked | AVE EXT | 1st trial RI-Test | 1.123 | 0.250 | 0.633 | 1.613 | 4.486 | 105 | 0.000 | 0.172 | 0.208 |
| SCR | CS+ | raw | not ranked | AVE last 2 trials EXT | 1st trial RI-Test | 0.386 | 0.238 | -0.080 | 0.852 | 1.627 | 105 | 0.107 | 0.023 | 0.024 |
| SCR | CS- | raw | not ranked | AVE ACQ | 1st trial EXT | 0.728 | 0.193 | 0.350 | 1.106 | 3.774 | 105 | 0.000 | 0.132 | 0.152 |
| SCR | CS- | raw | not ranked | AVE last 2 trials ACQ | 1st trial EXT | 0.520 | 0.150 | 0.226 | 0.814 | 3.469 | 105 | 0.001 | 0.086 | 0.094 |
| SCR | CS- | raw | not ranked | AVE ACQ | AVE EXT | 0.369 | 0.082 | 0.208 | 0.530 | 4.518 | 105 | 0.000 | 0.280 | 0.390 |
| SCR | CS- | raw | not ranked | AVE last 2 trials ACQ | AVE EXT | 0.231 | 0.074 | 0.086 | 0.376 | 3.126 | 105 | 0.002 | 0.140 | 0.163 |
| SCR | CS- | raw | not ranked | AVE ACQ | AVE last 2 trials EXT | 0.370 | 0.148 | 0.080 | 0.660 | 2.495 | 105 | 0.014 | 0.165 | 0.197 |
| SCR | CS- | raw | not ranked | AVE last 2 trials ACQ | AVE last 2 trials EXT | 0.265 | 0.107 | 0.055 | 0.475 | 2.475 | 105 | 0.015 | 0.107 | 0.120 |
| SCR | CS- | raw | not ranked | AVE ACQ | 1st trial RI-Test | 0.640 | 0.240 | 0.170 | 1.110 | 2.661 | 105 | 0.009 | 0.086 | 0.094 |
| SCR | CS- | raw | not ranked | AVE last 2 trials ACQ | 1st trial RI-Test | 0.449 | 0.239 | -0.019 | 0.917 | 1.878 | 105 | 0.063 | 0.054 | 0.057 |
| SCR | CS- | raw | not ranked | 1st trial EXT | 1st trial RI-Test | 0.336 | 0.118 | 0.105 | 0.567 | 2.849 | 105 | 0.005 | 0.096 | 0.106 |
| SCR | CS- | raw | not ranked | AVE EXT | 1st trial RI-Test | 0.584 | 0.299 | -0.002 | 1.170 | 1.953 | 105 | 0.054 | 0.035 | 0.036 |
| SCR | CS- | raw | not ranked | AVE last 2 trials EXT | 1st trial RI-Test | 0.145 | 0.319 | -0.480 | 0.770 | 0.453 | 105 | 0.651 | 0.004 | 0.004 |
| SCR | CS dis. | log | not ranked | AVE ACQ | 1st trial EXT | 0.328 | 0.134 | 0.065 | 0.591 | 2.446 | 105 | 0.016 | 0.037 | 0.039 |
| SCR | CS dis. | log | not ranked | AVE last 2 trials ACQ | 1st trial EXT | 0.260 | 0.085 | 0.093 | 0.427 | 3.045 | 105 | 0.003 | 0.062 | 0.066 |
| SCR | CS dis. | log | not ranked | AVE ACQ | AVE EXT | 0.109 | 0.056 | -0.001 | 0.219 | 1.956 | 105 | 0.053 | 0.047 | 0.050 |
| SCR | CS dis. | log | not ranked | AVE last 2 trials ACQ | AVE EXT | 0.031 | 0.031 | -0.030 | 0.092 | 1.000 | 105 | 0.320 | 0.010 | 0.010 |

| Outcome | Stim.-type | Ampl.-type | Ranking | Predictor | Criterion | *b* | $SE_b$ | Lower 95% CI | Upper 95% CI | *t* | *df* | *p* | $R^2$ | Cohen's $f^2$ |
|---|---|---|---|---|---|---|---|---|---|---|---|---|---|---|
| SCR | CS dis. | log | not ranked | AVE ACQ | AVE last 2 trials EXT | 0.074 | 0.103 | -0.128 | 0.276 | 0.719 | 105 | 0.474 | 0.006 | 0.006 |
| SCR | CS dis. | log | not ranked | AVE last 2 trials ACQ | AVE last 2 trials EXT | -0.039 | 0.105 | -0.245 | 0.167 | -0.373 | 105 | 0.710 | 0.004 | 0.004 |
| SCR | CS dis. | log | not ranked | AVE ACQ | 1st trial RI-Test | 0.135 | 0.259 | -0.373 | 0.643 | 0.521 | 105 | 0.603 | 0.004 | 0.004 |
| SCR | CS dis. | log | not ranked | AVE last 2 trials ACQ | 1st trial RI-Test | 0.173 | 0.221 | -0.260 | 0.606 | 0.784 | 105 | 0.435 | 0.018 | 0.018 |
| SCR | CS dis. | log | not ranked | 1st trial EXT | 1st trial RI-Test | 0.043 | 0.149 | -0.249 | 0.335 | 0.291 | 105 | 0.771 | 0.001 | 0.001 |
| SCR | CS dis. | log | not ranked | AVE EXT | 1st trial RI-Test | 0.149 | 0.450 | -0.733 | 1.031 | 0.331 | 105 | 0.741 | 0.001 | 0.001 |
| SCR | CS dis. | log | not ranked | AVE last 2 trials EXT | 1st trial RI-Test | -0.282 | 0.685 | -1.625 | 1.061 | -0.412 | 105 | 0.681 | 0.017 | 0.017 |
| SCR | CS+ | log | not ranked | AVE ACQ | 1st trial EXT | 0.679 | 0.115 | 0.454 | 0.904 | 5.906 | 105 | 0.000 | 0.297 | 0.423 |
| SCR | CS+ | log | not ranked | AVE last 2 trials ACQ | 1st trial EXT | 0.502 | 0.117 | 0.273 | 0.731 | 4.305 | 105 | 0.000 | 0.210 | 0.265 |
| SCR | CS+ | log | not ranked | AVE ACQ | AVE EXT | 0.294 | 0.074 | 0.149 | 0.439 | 3.995 | 105 | 0.000 | 0.277 | 0.383 |
| SCR | CS+ | log | not ranked | AVE last 2 trials ACQ | AVE EXT | 0.232 | 0.074 | 0.087 | 0.377 | 3.145 | 105 | 0.002 | 0.223 | 0.287 |
| SCR | CS+ | log | not ranked | AVE ACQ | AVE last 2 trials EXT | 0.202 | 0.094 | 0.018 | 0.386 | 2.158 | 105 | 0.033 | 0.117 | 0.133 |
| SCR | CS+ | log | not ranked | AVE last 2 trials ACQ | AVE last 2 trials EXT | 0.149 | 0.088 | -0.023 | 0.321 | 1.686 | 105 | 0.095 | 0.082 | 0.089 |
| SCR | CS+ | log | not ranked | AVE ACQ | 1st trial RI-Test | 0.659 | 0.123 | 0.418 | 0.900 | 5.361 | 105 | 0.000 | 0.216 | 0.275 |
| SCR | CS+ | log | not ranked | AVE last 2 trials ACQ | 1st trial RI-Test | 0.418 | 0.135 | 0.153 | 0.683 | 3.106 | 105 | 0.002 | 0.112 | 0.126 |
| SCR | CS+ | log | not ranked | 1st trial EXT | 1st trial RI-Test | 0.603 | 0.096 | 0.415 | 0.791 | 6.255 | 105 | 0.000 | 0.280 | 0.390 |
| SCR | CS+ | log | not ranked | AVE EXT | 1st trial RI-Test | 1.032 | 0.219 | 0.603 | 1.461 | 4.706 | 105 | 0.000 | 0.165 | 0.198 |
| SCR | CS+ | log | not ranked | AVE last 2 trials EXT | 1st trial RI-Test | 0.364 | 0.214 | -0.055 | 0.783 | 1.701 | 105 | 0.092 | 0.023 | 0.024 |
| SCR | CS- | log | not ranked | AVE ACQ | 1st trial EXT | 0.712 | 0.183 | 0.353 | 1.071 | 3.904 | 105 | 0.000 | 0.133 | 0.154 |
| SCR | CS- | log | not ranked | AVE last 2 trials ACQ | 1st trial EXT | 0.518 | 0.146 | 0.232 | 0.804 | 3.546 | 105 | 0.001 | 0.089 | 0.098 |
| SCR | CS- | log | not ranked | AVE ACQ | AVE EXT | 0.384 | 0.081 | 0.225 | 0.543 | 4.729 | 105 | 0.000 | 0.295 | 0.418 |
| SCR | CS- | log | not ranked | AVE last 2 trials ACQ | AVE EXT | 0.245 | 0.073 | 0.102 | 0.388 | 3.356 | 105 | 0.001 | 0.152 | 0.179 |
| SCR | CS- | log | not ranked | AVE ACQ | AVE last 2 trials EXT | 0.379 | 0.145 | 0.095 | 0.663 | 2.608 | 105 | 0.010 | 0.175 | 0.213 |
| SCR | CS- | log | not ranked | AVE last 2 trials ACQ | AVE last 2 trials EXT | 0.285 | 0.106 | 0.077 | 0.493 | 2.687 | 105 | 0.008 | 0.126 | 0.144 |
| SCR | CS- | log | not ranked | AVE ACQ | 1st trial RI-Test | 0.612 | 0.216 | 0.189 | 1.035 | 2.833 | 105 | 0.006 | 0.086 | 0.094 |

| Outcome | Stim.-type | Ampl.-type | Ranking | Predictor | Criterion | *b* | *SE*$_b$ | Lower 95% CI | Upper 95% CI | *t* | *df* | *p* | $R^2$ | Cohen's $f^2$ |
|---|---|---|---|---|---|---|---|---|---|---|---|---|---|---|
| SCR | CS- | log | not ranked | AVE last 2 trials ACQ | 1st trial RI-Test | 0.449 | 0.213 | 0.032 | 0.866 | 2.108 | 105 | 0.037 | 0.058 | 0.062 |
| SCR | CS- | log | not ranked | 1st trial EXT | 1st trial RI-Test | 0.341 | 0.113 | 0.120 | 0.562 | 3.021 | 105 | 0.003 | 0.101 | 0.113 |
| SCR | CS- | log | not ranked | AVE EXT | 1st trial RI-Test | 0.578 | 0.280 | 0.029 | 1.127 | 2.066 | 105 | 0.041 | 0.038 | 0.040 |
| SCR | CS- | log | not ranked | AVE last 2 trials EXT | 1st trial RI-Test | 0.163 | 0.287 | -0.400 | 0.726 | 0.568 | 105 | 0.571 | 0.005 | 0.005 |
| SCR | CS dis. | log rc | not ranked | AVE ACQ | 1st trial EXT | 0.378 | 0.200 | -0.014 | 0.770 | 1.894 | 105 | 0.061 | 0.030 | 0.031 |
| SCR | CS dis. | log rc | not ranked | AVE last 2 trials ACQ | 1st trial EXT | 0.298 | 0.109 | 0.084 | 0.512 | 2.733 | 105 | 0.007 | 0.046 | 0.049 |
| SCR | CS dis. | log rc | not ranked | AVE ACQ | AVE EXT | 0.071 | 0.054 | -0.035 | 0.177 | 1.328 | 105 | 0.187 | 0.017 | 0.017 |
| SCR | CS dis. | log rc | not ranked | AVE last 2 trials ACQ | AVE EXT | 0.022 | 0.038 | -0.052 | 0.096 | 0.572 | 105 | 0.568 | 0.004 | 0.004 |
| SCR | CS dis. | log rc | not ranked | AVE ACQ | AVE last 2 trials EXT | 0.103 | 0.106 | -0.105 | 0.311 | 0.974 | 105 | 0.332 | 0.010 | 0.010 |
| SCR | CS dis. | log rc | not ranked | AVE last 2 trials ACQ | AVE last 2 trials EXT | -0.007 | 0.093 | -0.189 | 0.175 | -0.073 | 105 | 0.942 | 0.000 | 0.000 |
| SCR | CS dis. | log rc | not ranked | AVE ACQ | 1st trial RI-Test | -0.212 | 0.225 | -0.653 | 0.229 | -0.943 | 105 | 0.348 | 0.009 | 0.009 |
| SCR | CS dis. | log rc | not ranked | AVE last 2 trials ACQ | 1st trial RI-Test | -0.041 | 0.158 | -0.351 | 0.269 | -0.259 | 105 | 0.796 | 0.001 | 0.001 |
| SCR | CS dis. | log rc | not ranked | 1st trial EXT | 1st trial RI-Test | 0.032 | 0.112 | -0.188 | 0.252 | 0.290 | 105 | 0.773 | 0.001 | 0.001 |
| SCR | CS dis. | log rc | not ranked | AVE EXT | 1st trial RI-Test | -0.025 | 0.381 | -0.772 | 0.722 | -0.066 | 105 | 0.948 | 0.000 | 0.000 |
| SCR | CS dis. | log rc | not ranked | AVE last 2 trials EXT | 1st trial RI-Test | -0.360 | 0.376 | -1.097 | 0.377 | -0.958 | 105 | 0.340 | 0.027 | 0.028 |
| SCR | CS+ | log rc | not ranked | AVE ACQ | 1st trial EXT | 0.435 | 0.122 | 0.196 | 0.674 | 3.564 | 105 | 0.001 | 0.108 | 0.121 |
| SCR | CS+ | log rc | not ranked | AVE last 2 trials ACQ | 1st trial EXT | 0.320 | 0.111 | 0.102 | 0.538 | 2.886 | 105 | 0.005 | 0.069 | 0.074 |
| SCR | CS+ | log rc | not ranked | AVE ACQ | AVE EXT | 0.267 | 0.058 | 0.153 | 0.381 | 4.578 | 105 | 0.000 | 0.215 | 0.274 |
| SCR | CS+ | log rc | not ranked | AVE last 2 trials ACQ | AVE EXT | 0.239 | 0.059 | 0.123 | 0.355 | 4.069 | 105 | 0.000 | 0.204 | 0.256 |
| SCR | CS+ | log rc | not ranked | AVE ACQ | AVE last 2 trials EXT | 0.181 | 0.073 | 0.038 | 0.324 | 2.495 | 105 | 0.014 | 0.079 | 0.086 |
| SCR | CS+ | log rc | not ranked | AVE last 2 trials ACQ | AVE last 2 trials EXT | 0.157 | 0.069 | 0.022 | 0.292 | 2.268 | 105 | 0.025 | 0.070 | 0.075 |
| SCR | CS+ | log rc | not ranked | AVE ACQ | 1st trial RI-Test | 0.233 | 0.138 | -0.037 | 0.503 | 1.681 | 105 | 0.096 | 0.023 | 0.024 |
| SCR | CS+ | log rc | not ranked | AVE last 2 trials ACQ | 1st trial RI-Test | 0.070 | 0.130 | -0.185 | 0.325 | 0.543 | 105 | 0.588 | 0.003 | 0.003 |
| SCR | CS+ | log rc | not ranked | 1st trial EXT | 1st trial RI-Test | 0.403 | 0.103 | 0.201 | 0.605 | 3.910 | 105 | 0.000 | 0.122 | 0.138 |
| SCR | CS+ | log rc | not ranked | AVE EXT | 1st trial RI-Test | 0.500 | 0.206 | 0.096 | 0.904 | 2.425 | 105 | 0.017 | 0.035 | 0.037 |

| Outcome | Stim.-type | Ampl.-type | Ranking | Predictor | Criterion | *b* | $SE_b$ | Lower 95% CI | Upper 95% CI | *t* | *df* | *p* | $R^2$ | Cohen's $f^2$ |
|---|---|---|---|---|---|---|---|---|---|---|---|---|---|---|
| SCR | CS+ | log rc | not ranked | AVE last 2 trials EXT | 1st trial RI-Test | -0.021 | 0.196 | -0.405 | 0.363 | -0.106 | 105 | 0.916 | 0.000 | 0.000 |
| SCR | CS- | log rc | not ranked | AVE ACQ | 1st trial EXT | 0.247 | 0.193 | -0.131 | 0.625 | 1.278 | 105 | 0.204 | 0.014 | 0.014 |
| SCR | CS- | log rc | not ranked | AVE last 2 trials ACQ | 1st trial EXT | 0.192 | 0.150 | -0.102 | 0.486 | 1.275 | 105 | 0.205 | 0.010 | 0.011 |
| SCR | CS- | log rc | not ranked | AVE ACQ | AVE EXT | 0.370 | 0.078 | 0.217 | 0.523 | 4.719 | 105 | 0.000 | 0.272 | 0.375 |
| SCR | CS- | log rc | not ranked | AVE last 2 trials ACQ | AVE EXT | 0.246 | 0.069 | 0.111 | 0.381 | 3.577 | 105 | 0.001 | 0.151 | 0.178 |
| SCR | CS- | log rc | not ranked | AVE ACQ | AVE last 2 trials EXT | 0.310 | 0.104 | 0.106 | 0.514 | 2.971 | 105 | 0.004 | 0.125 | 0.143 |
| SCR | CS- | log rc | not ranked | AVE last 2 trials ACQ | AVE last 2 trials EXT | 0.246 | 0.078 | 0.093 | 0.399 | 3.143 | 105 | 0.002 | 0.099 | 0.110 |
| SCR | CS- | log rc | not ranked | AVE ACQ | 1st trial RI-Test | 0.397 | 0.240 | -0.073 | 0.867 | 1.654 | 105 | 0.101 | 0.031 | 0.032 |
| SCR | CS- | log rc | not ranked | AVE last 2 trials ACQ | 1st trial RI-Test | 0.255 | 0.216 | -0.168 | 0.678 | 1.179 | 105 | 0.241 | 0.016 | 0.017 |
| SCR | CS- | log rc | not ranked | 1st trial EXT | 1st trial RI-Test | 0.192 | 0.118 | -0.039 | 0.423 | 1.619 | 105 | 0.108 | 0.032 | 0.033 |
| SCR | CS- | log rc | not ranked | AVE EXT | 1st trial RI-Test | 0.178 | 0.278 | -0.367 | 0.723 | 0.639 | 105 | 0.524 | 0.003 | 0.003 |
| SCR | CS- | log rc | not ranked | AVE last 2 trials EXT | 1st trial RI-Test | -0.108 | 0.189 | -0.478 | 0.262 | -0.569 | 105 | 0.571 | 0.002 | 0.002 |
| SCR | CS dis. | raw | ranked | AVE ACQ | 1st trial EXT | 0.180 | 0.089 | 0.006 | 0.354 | 2.009 | 105 | 0.047 | 0.032 | 0.033 |
| SCR | CS dis. | raw | ranked | AVE last 2 trials ACQ | 1st trial EXT | 0.273 | 0.086 | 0.104 | 0.442 | 3.167 | 105 | 0.002 | 0.087 | 0.095 |
| SCR | CS dis. | raw | ranked | AVE ACQ | AVE EXT | 0.211 | 0.097 | 0.021 | 0.401 | 2.169 | 105 | 0.032 | 0.043 | 0.045 |
| SCR | CS dis. | raw | ranked | AVE last 2 trials ACQ | AVE EXT | 0.228 | 0.092 | 0.048 | 0.408 | 2.489 | 105 | 0.014 | 0.059 | 0.063 |
| SCR | CS dis. | raw | ranked | AVE ACQ | AVE last 2 trials EXT | 0.125 | 0.120 | -0.110 | 0.360 | 1.045 | 105 | 0.299 | 0.012 | 0.012 |
| SCR | CS dis. | raw | ranked | AVE last 2 trials ACQ | AVE last 2 trials EXT | 0.200 | 0.106 | -0.008 | 0.408 | 1.888 | 105 | 0.062 | 0.036 | 0.038 |
| SCR | CS dis. | raw | ranked | AVE ACQ | 1st trial RI-Test | 0.037 | 0.103 | -0.165 | 0.239 | 0.362 | 105 | 0.718 | 0.001 | 0.001 |
| SCR | CS dis. | raw | ranked | AVE last 2 trials ACQ | 1st trial RI-Test | -0.071 | 0.096 | -0.259 | 0.117 | -0.740 | 105 | 0.461 | 0.006 | 0.006 |
| SCR | CS dis. | raw | ranked | 1st trial EXT | 1st trial RI-Test | 0.034 | 0.112 | -0.186 | 0.254 | 0.303 | 105 | 0.763 | 0.001 | 0.001 |
| SCR | CS dis. | raw | ranked | AVE EXT | 1st trial RI-Test | 0.017 | 0.109 | -0.197 | 0.231 | 0.154 | 105 | 0.878 | 0.000 | 0.000 |
| SCR | CS dis. | raw | ranked | AVE last 2 trials EXT | 1st trial RI-Test | 0.068 | 0.087 | -0.103 | 0.239 | 0.773 | 105 | 0.442 | 0.006 | 0.006 |
| SCR | CS+ | raw | ranked | AVE ACQ | 1st trial EXT | 0.594 | 0.075 | 0.447 | 0.741 | 7.958 | 105 | 0.000 | 0.319 | 0.469 |
| SCR | CS+ | raw | ranked | AVE last 2 trials ACQ | 1st trial EXT | 0.381 | 0.078 | 0.228 | 0.534 | 4.860 | 105 | 0.000 | 0.187 | 0.230 |
| SCR | CS+ | raw | ranked | AVE ACQ | AVE EXT | 0.607 | 0.071 | 0.468 | 0.746 | 8.500 | 105 | 0.000 | 0.324 | 0.480 |

| Outcome | Stim.-type | Ampl.-type | Ranking | Predictor | Criterion | *b* | $SE_b$ | Lower 95% CI | Upper 95% CI | *t* | *df* | *p* | $R^2$ | Cohen's $f^2$ |
|---|---|---|---|---|---|---|---|---|---|---|---|---|---|---|
| SCR | CS+ | raw | ranked | AVE last 2 trials ACQ | AVE EXT | 0.451 | 0.077 | 0.300 | 0.602 | 5.852 | 105 | 0.000 | 0.256 | 0.343 |
| SCR | CS+ | raw | ranked | AVE ACQ | AVE last 2 trials EXT | 0.364 | 0.125 | 0.119 | 0.609 | 2.912 | 105 | 0.004 | 0.072 | 0.078 |
| SCR | CS+ | raw | ranked | AVE last 2 trials ACQ | AVE last 2 trials EXT | 0.281 | 0.108 | 0.069 | 0.493 | 2.608 | 105 | 0.010 | 0.061 | 0.065 |
| SCR | CS+ | raw | ranked | AVE ACQ | 1st trial RI-Test | 0.485 | 0.083 | 0.322 | 0.648 | 5.828 | 105 | 0.000 | 0.215 | 0.274 |
| SCR | CS+ | raw | ranked | AVE last 2 trials ACQ | 1st trial RI-Test | 0.216 | 0.088 | 0.044 | 0.388 | 2.441 | 105 | 0.016 | 0.061 | 0.064 |
| SCR | CS+ | raw | ranked | 1st trial EXT | 1st trial RI-Test | 0.518 | 0.083 | 0.355 | 0.681 | 6.282 | 105 | 0.000 | 0.272 | 0.374 |
| SCR | CS+ | raw | ranked | AVE EXT | 1st trial RI-Test | 0.340 | 0.097 | 0.150 | 0.530 | 3.507 | 105 | 0.001 | 0.120 | 0.136 |
| SCR | CS+ | raw | ranked | AVE last 2 trials EXT | 1st trial RI-Test | 0.009 | 0.075 | -0.138 | 0.156 | 0.113 | 105 | 0.910 | 0.000 | 0.000 |
| SCR | CS- | raw | ranked | AVE ACQ | 1st trial EXT | 0.388 | 0.096 | 0.200 | 0.576 | 4.057 | 105 | 0.000 | 0.129 | 0.148 |
| SCR | CS- | raw | ranked | AVE last 2 trials ACQ | 1st trial EXT | 0.196 | 0.078 | 0.043 | 0.349 | 2.507 | 105 | 0.014 | 0.056 | 0.060 |
| SCR | CS- | raw | ranked | AVE ACQ | AVE EXT | 0.670 | 0.070 | 0.533 | 0.807 | 9.586 | 105 | 0.000 | 0.384 | 0.623 |
| SCR | CS- | raw | ranked | AVE last 2 trials ACQ | AVE EXT | 0.353 | 0.072 | 0.212 | 0.494 | 4.905 | 105 | 0.000 | 0.184 | 0.225 |
| SCR | CS- | raw | ranked | AVE ACQ | AVE last 2 trials EXT | 0.427 | 0.115 | 0.202 | 0.652 | 3.702 | 105 | 0.000 | 0.109 | 0.122 |
| SCR | CS- | raw | ranked | AVE last 2 trials ACQ | AVE last 2 trials EXT | 0.388 | 0.094 | 0.204 | 0.572 | 4.117 | 105 | 0.000 | 0.155 | 0.183 |
| SCR | CS- | raw | ranked | AVE ACQ | 1st trial RI-Test | 0.340 | 0.104 | 0.136 | 0.544 | 3.281 | 105 | 0.001 | 0.099 | 0.110 |
| SCR | CS- | raw | ranked | AVE last 2 trials ACQ | 1st trial RI-Test | 0.206 | 0.080 | 0.049 | 0.363 | 2.567 | 105 | 0.012 | 0.062 | 0.067 |
| SCR | CS- | raw | ranked | 1st trial EXT | 1st trial RI-Test | 0.327 | 0.100 | 0.131 | 0.523 | 3.265 | 105 | 0.001 | 0.107 | 0.119 |
| SCR | CS- | raw | ranked | AVE EXT | 1st trial RI-Test | 0.298 | 0.096 | 0.110 | 0.486 | 3.096 | 105 | 0.003 | 0.089 | 0.097 |
| SCR | CS- | raw | ranked | AVE last 2 trials EXT | 1st trial RI-Test | 0.110 | 0.069 | -0.025 | 0.245 | 1.583 | 105 | 0.117 | 0.017 | 0.018 |
| SCR | CS dis. | log | ranked | AVE ACQ | 1st trial EXT | 0.177 | 0.090 | 0.001 | 0.353 | 1.971 | 105 | 0.051 | 0.031 | 0.032 |
| SCR | CS dis. | log | ranked | AVE last 2 trials ACQ | 1st trial EXT | 0.269 | 0.086 | 0.100 | 0.438 | 3.135 | 105 | 0.002 | 0.084 | 0.092 |
| SCR | CS dis. | log | ranked | AVE ACQ | AVE EXT | 0.206 | 0.098 | 0.014 | 0.398 | 2.108 | 105 | 0.037 | 0.041 | 0.043 |
| SCR | CS dis. | log | ranked | AVE last 2 trials ACQ | AVE EXT | 0.214 | 0.092 | 0.034 | 0.394 | 2.319 | 105 | 0.022 | 0.052 | 0.055 |
| SCR | CS dis. | log | ranked | AVE ACQ | AVE last 2 trials EXT | 0.132 | 0.120 | -0.103 | 0.367 | 1.102 | 105 | 0.273 | 0.014 | 0.014 |
| SCR | CS dis. | log | ranked | AVE last 2 trials ACQ | AVE last 2 trials EXT | 0.194 | 0.106 | -0.014 | 0.402 | 1.838 | 105 | 0.069 | 0.034 | 0.036 |
| SCR | CS dis. | log | ranked | AVE ACQ | 1st trial RI-Test | 0.039 | 0.103 | -0.163 | 0.241 | 0.380 | 105 | 0.704 | 0.002 | 0.002 |

| Outcome | Stim.-type | Ampl.-type | Ranking | Predictor | Criterion | *b* | $SE_b$ | Lower 95% CI | Upper 95% CI | *t* | *df* | *p* | $R^2$ | Cohen's $f^2$ |
|---|---|---|---|---|---|---|---|---|---|---|---|---|---|---|
| SCR | CS dis. | log | ranked | AVE last 2 trials ACQ | 1st trial RI-Test | -0.081 | 0.096 | -0.269 | 0.107 | -0.845 | 105 | 0.400 | 0.008 | 0.008 |
| SCR | CS dis. | log | ranked | 1st trial EXT | 1st trial RI-Test | 0.030 | 0.110 | -0.186 | 0.246 | 0.270 | 105 | 0.787 | 0.001 | 0.001 |
| SCR | CS dis. | log | ranked | AVE EXT | 1st trial RI-Test | 0.009 | 0.109 | -0.205 | 0.223 | 0.084 | 105 | 0.933 | 0.000 | 0.000 |
| SCR | CS dis. | log | ranked | AVE last 2 trials EXT | 1st trial RI-Test | 0.060 | 0.087 | -0.111 | 0.231 | 0.696 | 105 | 0.488 | 0.005 | 0.005 |
| SCR | CS+ | log | ranked | AVE ACQ | 1st trial EXT | 0.591 | 0.075 | 0.444 | 0.738 | 7.906 | 105 | 0.000 | 0.316 | 0.462 |
| SCR | CS+ | log | ranked | AVE last 2 trials ACQ | 1st trial EXT | 0.382 | 0.078 | 0.229 | 0.535 | 4.880 | 105 | 0.000 | 0.188 | 0.231 |
| SCR | CS+ | log | ranked | AVE ACQ | AVE EXT | 0.606 | 0.073 | 0.463 | 0.749 | 8.363 | 105 | 0.000 | 0.324 | 0.479 |
| SCR | CS+ | log | ranked | AVE last 2 trials ACQ | AVE EXT | 0.455 | 0.077 | 0.304 | 0.606 | 5.942 | 105 | 0.000 | 0.260 | 0.351 |
| SCR | CS+ | log | ranked | AVE ACQ | AVE last 2 trials EXT | 0.375 | 0.125 | 0.130 | 0.620 | 3.003 | 105 | 0.003 | 0.077 | 0.083 |
| SCR | CS+ | log | ranked | AVE last 2 trials ACQ | AVE last 2 trials EXT | 0.285 | 0.108 | 0.073 | 0.497 | 2.646 | 105 | 0.009 | 0.063 | 0.067 |
| SCR | CS+ | log | ranked | AVE ACQ | 1st trial RI-Test | 0.484 | 0.084 | 0.319 | 0.649 | 5.789 | 105 | 0.000 | 0.214 | 0.272 |
| SCR | CS+ | log | ranked | AVE last 2 trials ACQ | 1st trial RI-Test | 0.221 | 0.088 | 0.049 | 0.393 | 2.514 | 105 | 0.013 | 0.063 | 0.068 |
| SCR | CS+ | log | ranked | 1st trial EXT | 1st trial RI-Test | 0.518 | 0.083 | 0.355 | 0.681 | 6.282 | 105 | 0.000 | 0.272 | 0.374 |
| SCR | CS+ | log | ranked | AVE EXT | 1st trial RI-Test | 0.337 | 0.097 | 0.147 | 0.527 | 3.488 | 105 | 0.001 | 0.118 | 0.134 |
| SCR | CS+ | log | ranked | AVE last 2 trials EXT | 1st trial RI-Test | 0.008 | 0.075 | -0.139 | 0.155 | 0.110 | 105 | 0.912 | 0.000 | 0.000 |
| SCR | CS- | log | ranked | AVE ACQ | 1st trial EXT | 0.387 | 0.095 | 0.201 | 0.573 | 4.057 | 105 | 0.000 | 0.128 | 0.147 |
| SCR | CS- | log | ranked | AVE last 2 trials ACQ | 1st trial EXT | 0.197 | 0.078 | 0.044 | 0.350 | 2.520 | 105 | 0.013 | 0.057 | 0.060 |
| SCR | CS- | log | ranked | AVE ACQ | AVE EXT | 0.674 | 0.070 | 0.537 | 0.811 | 9.688 | 105 | 0.000 | 0.388 | 0.634 |
| SCR | CS- | log | ranked | AVE last 2 trials ACQ | AVE EXT | 0.356 | 0.072 | 0.215 | 0.497 | 4.959 | 105 | 0.000 | 0.187 | 0.230 |
| SCR | CS- | log | ranked | AVE ACQ | AVE last 2 trials EXT | 0.432 | 0.115 | 0.207 | 0.657 | 3.751 | 105 | 0.000 | 0.111 | 0.125 |
| SCR | CS- | log | ranked | AVE last 2 trials ACQ | AVE last 2 trials EXT | 0.391 | 0.094 | 0.207 | 0.575 | 4.151 | 105 | 0.000 | 0.157 | 0.186 |
| SCR | CS- | log | ranked | AVE ACQ | 1st trial RI-Test | 0.341 | 0.104 | 0.137 | 0.545 | 3.289 | 105 | 0.001 | 0.099 | 0.110 |
| SCR | CS- | log | ranked | AVE last 2 trials ACQ | 1st trial RI-Test | 0.206 | 0.080 | 0.049 | 0.363 | 2.573 | 105 | 0.011 | 0.063 | 0.067 |
| SCR | CS- | log | ranked | 1st trial EXT | 1st trial RI-Test | 0.327 | 0.100 | 0.131 | 0.523 | 3.265 | 105 | 0.001 | 0.107 | 0.119 |
| SCR | CS- | log | ranked | AVE EXT | 1st trial RI-Test | 0.298 | 0.096 | 0.110 | 0.486 | 3.108 | 105 | 0.002 | 0.089 | 0.097 |
| SCR | CS- | log | ranked | AVE last 2 trials EXT | 1st trial RI-Test | 0.109 | 0.069 | -0.026 | 0.244 | 1.578 | 105 | 0.118 | 0.017 | 0.017 |
| SCR | CS dis. | log rc | ranked | AVE ACQ | 1st trial EXT | 0.150 | 0.096 | -0.038 | 0.338 | 1.571 | 105 | 0.119 | 0.023 | 0.023 |

| Outcome | Stim.-type | Ampl.-type | Ranking | Predictor | Criterion | *b* | $SE_b$ | Lower 95% CI | Upper 95% CI | *t* | *df* | *p* | $R^2$ | Cohen's $f^2$ |
|---|---|---|---|---|---|---|---|---|---|---|---|---|---|---|
| SCR | CS dis. | log rc | ranked | AVE last 2 trials ACQ | 1st trial EXT | 0.248 | 0.085 | 0.081 | 0.415 | 2.930 | 105 | 0.004 | 0.071 | 0.077 |
| SCR | CS dis. | log rc | ranked | AVE ACQ | AVE EXT | 0.136 | 0.096 | -0.052 | 0.324 | 1.411 | 105 | 0.161 | 0.018 | 0.018 |
| SCR | CS dis. | log rc | ranked | AVE last 2 trials ACQ | AVE EXT | 0.164 | 0.094 | -0.020 | 0.348 | 1.739 | 105 | 0.085 | 0.031 | 0.032 |
| SCR | CS dis. | log rc | ranked | AVE ACQ | AVE last 2 trials EXT | 0.135 | 0.120 | -0.100 | 0.370 | 1.130 | 105 | 0.261 | 0.014 | 0.015 |
| SCR | CS dis. | log rc | ranked | AVE last 2 trials ACQ | AVE last 2 trials EXT | 0.167 | 0.105 | -0.039 | 0.373 | 1.594 | 105 | 0.114 | 0.025 | 0.026 |
| SCR | CS dis. | log rc | ranked | AVE ACQ | 1st trial RI-Test | -0.038 | 0.100 | -0.234 | 0.158 | -0.381 | 105 | 0.704 | 0.001 | 0.001 |
| SCR | CS dis. | log rc | ranked | AVE last 2 trials ACQ | 1st trial RI-Test | -0.099 | 0.093 | -0.281 | 0.083 | -1.064 | 105 | 0.290 | 0.011 | 0.012 |
| SCR | CS dis. | log rc | ranked | 1st trial EXT | 1st trial RI-Test | 0.040 | 0.100 | -0.156 | 0.236 | 0.399 | 105 | 0.691 | 0.002 | 0.002 |
| SCR | CS dis. | log rc | ranked | AVE EXT | 1st trial RI-Test | -0.014 | 0.101 | -0.212 | 0.184 | -0.137 | 105 | 0.892 | 0.000 | 0.000 |
| SCR | CS dis. | log rc | ranked | AVE last 2 trials EXT | 1st trial RI-Test | -0.010 | 0.084 | -0.175 | 0.155 | -0.121 | 105 | 0.904 | 0.000 | 0.000 |
| SCR | CS+ | log rc | ranked | AVE ACQ | 1st trial EXT | 0.358 | 0.096 | 0.170 | 0.546 | 3.722 | 105 | 0.000 | 0.116 | 0.131 |
| SCR | CS+ | log rc | ranked | AVE last 2 trials ACQ | 1st trial EXT | 0.244 | 0.082 | 0.083 | 0.405 | 2.957 | 105 | 0.004 | 0.076 | 0.083 |
| SCR | CS+ | log rc | ranked | AVE ACQ | AVE EXT | 0.558 | 0.089 | 0.384 | 0.732 | 6.264 | 105 | 0.000 | 0.274 | 0.377 |
| SCR | CS+ | log rc | ranked | AVE last 2 trials ACQ | AVE EXT | 0.437 | 0.076 | 0.288 | 0.586 | 5.786 | 105 | 0.000 | 0.240 | 0.316 |
| SCR | CS+ | log rc | ranked | AVE ACQ | AVE last 2 trials EXT | 0.397 | 0.131 | 0.140 | 0.654 | 3.044 | 105 | 0.003 | 0.086 | 0.094 |
| SCR | CS+ | log rc | ranked | AVE last 2 trials ACQ | AVE last 2 trials EXT | 0.299 | 0.110 | 0.083 | 0.515 | 2.729 | 105 | 0.007 | 0.069 | 0.075 |
| SCR | CS+ | log rc | ranked | AVE ACQ | 1st trial RI-Test | 0.200 | 0.097 | 0.010 | 0.390 | 2.058 | 105 | 0.042 | 0.037 | 0.038 |
| SCR | CS+ | log rc | ranked | AVE last 2 trials ACQ | 1st trial RI-Test | 0.074 | 0.085 | -0.093 | 0.241 | 0.869 | 105 | 0.387 | 0.007 | 0.007 |
| SCR | CS+ | log rc | ranked | 1st trial EXT | 1st trial RI-Test | 0.349 | 0.097 | 0.159 | 0.539 | 3.587 | 105 | 0.001 | 0.124 | 0.141 |
| SCR | CS+ | log rc | ranked | AVE EXT | 1st trial RI-Test | 0.161 | 0.096 | -0.027 | 0.349 | 1.681 | 105 | 0.096 | 0.027 | 0.028 |
| SCR | CS+ | log rc | ranked | AVE last 2 trials EXT | 1st trial RI-Test | -0.067 | 0.071 | -0.206 | 0.072 | -0.937 | 105 | 0.351 | 0.007 | 0.008 |
| SCR | CS- | log rc | ranked | AVE ACQ | 1st trial EXT | 0.244 | 0.100 | 0.048 | 0.440 | 2.446 | 105 | 0.016 | 0.051 | 0.053 |
| SCR | CS- | log rc | ranked | AVE last 2 trials ACQ | 1st trial EXT | 0.111 | 0.078 | -0.042 | 0.264 | 1.418 | 105 | 0.159 | 0.018 | 0.018 |
| SCR | CS- | log rc | ranked | AVE ACQ | AVE EXT | 0.682 | 0.072 | 0.541 | 0.823 | 9.479 | 105 | 0.000 | 0.397 | 0.659 |
| SCR | CS- | log rc | ranked | AVE last 2 trials ACQ | AVE EXT | 0.347 | 0.071 | 0.208 | 0.486 | 4.913 | 105 | 0.000 | 0.177 | 0.215 |
| SCR | CS- | log rc | ranked | AVE ACQ | AVE last 2 trials EXT | 0.487 | 0.117 | 0.258 | 0.716 | 4.148 | 105 | 0.000 | 0.141 | 0.164 |
| SCR | CS- | log rc | ranked | AVE last 2 trials ACQ | AVE last 2 trials EXT | 0.383 | 0.093 | 0.201 | 0.565 | 4.107 | 105 | 0.000 | 0.150 | 0.177 |

| Outcome | Stim.-type | Ampl.-type | Ranking | Predictor | Criterion | *b* | *SE_b* | Lower 95% CI | Upper 95% CI | *t* | *df* | *p* | $R^2$ | Cohen's $f^2$ |
|---|---|---|---|---|---|---|---|---|---|---|---|---|---|---|
| SCR | CS- | log rc | ranked | AVE ACQ | 1st trial RI-Test | 0.251 | 0.097 | 0.061 | 0.441 | 2.582 | 105 | 0.011 | 0.054 | 0.057 |
| SCR | CS- | log rc | ranked | AVE last 2 trials ACQ | 1st trial RI-Test | 0.146 | 0.080 | -0.011 | 0.303 | 1.815 | 105 | 0.072 | 0.031 | 0.032 |
| SCR | CS- | log rc | ranked | 1st trial EXT | 1st trial RI-Test | 0.189 | 0.104 | -0.015 | 0.393 | 1.822 | 105 | 0.071 | 0.036 | 0.037 |
| SCR | CS- | log rc | ranked | AVE EXT | 1st trial RI-Test | 0.145 | 0.100 | -0.051 | 0.341 | 1.454 | 105 | 0.149 | 0.021 | 0.022 |
| SCR | CS- | log rc | ranked | AVE last 2 trials EXT | 1st trial RI-Test | 0.039 | 0.070 | -0.098 | 0.176 | 0.556 | 105 | 0.579 | 0.002 | 0.002 |

Note. Ampl. = Amplitude, Stim. = Stimulus, CI = Confidence Interval, CS dis. = CS discrimination, log = log-transformed, log rc = log-transformed and range corrected, AVE = average, ACQ = Acquisition training, EXT = Extinction training, RI = Reinstatement, RI-Test = Reinstatement-Test.

***Supplementary Table 8:*** *Detailed results of linear regressions: fear ratings.*

| Outcome | Stim.-type | Ranking | Predictor | Criterion | *b* | *SE_b* | Lower 95% CI | Upper 95% CI | *t* | *df* | *p* | $R^2$ | Cohen's $f^2$ |
|---|---|---|---|---|---|---|---|---|---|---|---|---|---|
| Fear Ratings | CS dis. | not ranked | post-pre ACQ | pre EXT | 0.519 | 0.094 | 0.335 | 0.703 | 5.543 | 77 | 0.000 | 0.270 | 0.369 |
| Fear Ratings | CS dis. | not ranked | post ACQ | pre EXT | 0.570 | 0.088 | 0.398 | 0.742 | 6.454 | 92 | 0.000 | 0.265 | 0.360 |
| Fear Ratings | CS dis. | not ranked | post-pre ACQ | pre-post EXT | 0.372 | 0.132 | 0.113 | 0.631 | 2.827 | 76 | 0.006 | 0.175 | 0.213 |
| Fear Ratings | CS dis. | not ranked | post ACQ | pre-post EXT | 0.413 | 0.107 | 0.203 | 0.623 | 3.851 | 91 | 0.000 | 0.177 | 0.215 |
| Fear Ratings | CS dis. | not ranked | post-pre ACQ | post EXT | 0.144 | 0.087 | -0.027 | 0.315 | 1.648 | 79 | 0.103 | 0.076 | 0.082 |
| Fear Ratings | CS dis. | not ranked | post ACQ | post EXT | 0.139 | 0.077 | -0.012 | 0.290 | 1.818 | 98 | 0.072 | 0.063 | 0.068 |
| Fear Ratings | CS dis. | not ranked | post ACQ | 1st trial RI-Test | 0.257 | 0.084 | 0.092 | 0.422 | 3.056 | 74 | 0.003 | 0.099 | 0.110 |
| Fear Ratings | CS dis. | not ranked | post-pre ACQ | 1st trial RI-Test | 0.240 | 0.081 | 0.081 | 0.399 | 2.967 | 60 | 0.004 | 0.117 | 0.132 |
| Fear Ratings | CS dis. | not ranked | pre EXT | 1st trial RI-Test | 0.236 | 0.086 | 0.067 | 0.405 | 2.739 | 69 | 0.008 | 0.112 | 0.127 |
| Fear Ratings | CS dis. | not ranked | pre-post EXT | 1st trial RI-Test | 0.187 | 0.104 | -0.017 | 0.391 | 1.793 | 68 | 0.077 | 0.052 | 0.055 |
| Fear Ratings | CS dis. | not ranked | post EXT | 1st trial RI-Test | 0.301 | 0.187 | -0.066 | 0.668 | 1.610 | 71 | 0.112 | 0.043 | 0.045 |
| Fear Ratings | CS+ | not ranked | post-pre ACQ | pre EXT | 0.509 | 0.095 | 0.323 | 0.695 | 5.365 | 91 | 0.000 | 0.208 | 0.263 |
| Fear Ratings | CS+ | not ranked | post ACQ | pre EXT | 0.655 | 0.090 | 0.479 | 0.831 | 7.307 | 97 | 0.000 | 0.319 | 0.469 |
| Fear Ratings | CS+ | not ranked | post-pre ACQ | pre-post EXT | 0.425 | 0.081 | 0.266 | 0.584 | 5.218 | 90 | 0.000 | 0.194 | 0.241 |
| Fear Ratings | CS+ | not ranked | post ACQ | pre-post EXT | 0.461 | 0.083 | 0.298 | 0.624 | 5.547 | 96 | 0.000 | 0.209 | 0.263 |
| Fear Ratings | CS+ | not ranked | post-pre ACQ | post EXT | 0.084 | 0.069 | -0.051 | 0.219 | 1.212 | 92 | 0.229 | 0.011 | 0.011 |
| Fear Ratings | CS+ | not ranked | post ACQ | post EXT | 0.172 | 0.073 | 0.029 | 0.315 | 2.370 | 101 | 0.020 | 0.042 | 0.044 |

| Outcome | Stim.-type | Ranking | Predictor | Criterion | *b* | $SE_b$ | Lower 95% CI | Upper 95% CI | *t* | *df* | *p* | $R^2$ | Cohen's $f^2$ |
|---|---|---|---|---|---|---|---|---|---|---|---|---|---|
| Fear Ratings | CS+ | not ranked | post ACQ | 1st trial RI-Test | 0.503 | 0.102 | 0.303 | 0.703 | 4.928 | 85 | 0.000 | 0.171 | 0.207 |
| Fear Ratings | CS+ | not ranked | post-pre ACQ | 1st trial RI-Test | 0.330 | 0.105 | 0.124 | 0.536 | 3.153 | 79 | 0.002 | 0.091 | 0.100 |
| Fear Ratings | CS+ | not ranked | pre EXT | 1st trial RI-Test | 0.430 | 0.093 | 0.248 | 0.612 | 4.630 | 82 | 0.000 | 0.184 | 0.226 |
| Fear Ratings | CS+ | not ranked | pre-post EXT | 1st trial RI-Test | 0.293 | 0.119 | 0.060 | 0.526 | 2.455 | 81 | 0.016 | 0.060 | 0.064 |
| Fear Ratings | CS+ | not ranked | post EXT | 1st trial RI-Test | 0.352 | 0.134 | 0.089 | 0.615 | 2.615 | 84 | 0.011 | 0.072 | 0.077 |
| Fear Ratings | CS- | not ranked | post-pre ACQ | pre EXT | 0.077 | 0.099 | -0.117 | 0.271 | 0.773 | 86 | 0.442 | 0.019 | 0.020 |
| Fear Ratings | CS- | not ranked | post ACQ | pre EXT | 0.244 | 0.101 | 0.046 | 0.442 | 2.423 | 98 | 0.017 | 0.130 | 0.150 |
| Fear Ratings | CS- | not ranked | post-pre ACQ | pre-post EXT | -0.197 | 0.120 | -0.432 | 0.038 | -1.638 | 85 | 0.105 | 0.061 | 0.065 |
| Fear Ratings | CS- | not ranked | post ACQ | pre-post EXT | -0.147 | 0.123 | -0.388 | 0.094 | -1.198 | 97 | 0.234 | 0.030 | 0.031 |
| Fear Ratings | CS- | not ranked | post-pre ACQ | post EXT | 0.236 | 0.138 | -0.034 | 0.506 | 1.708 | 88 | 0.091 | 0.090 | 0.099 |
| Fear Ratings | CS- | not ranked | post ACQ | post EXT | 0.386 | 0.140 | 0.112 | 0.660 | 2.753 | 100 | 0.007 | 0.217 | 0.278 |
| Fear Ratings | CS- | not ranked | post ACQ | 1st trial RI-Test | 0.270 | 0.176 | -0.075 | 0.615 | 1.534 | 88 | 0.129 | 0.039 | 0.040 |
| Fear Ratings | CS- | not ranked | post-pre ACQ | 1st trial RI-Test | 0.082 | 0.160 | -0.232 | 0.396 | 0.513 | 78 | 0.610 | 0.004 | 0.004 |
| Fear Ratings | CS- | not ranked | pre EXT | 1st trial RI-Test | 0.493 | 0.275 | -0.046 | 1.032 | 1.788 | 86 | 0.077 | 0.043 | 0.045 |
| Fear Ratings | CS- | not ranked | pre-post EXT | 1st trial RI-Test | -0.279 | 0.213 | -0.696 | 0.138 | -1.308 | 85 | 0.194 | 0.028 | 0.029 |
| Fear Ratings | CS- | not ranked | post EXT | 1st trial RI-Test | 0.582 | 0.193 | 0.204 | 0.960 | 3.010 | 87 | 0.003 | 0.109 | 0.122 |
| Fear Ratings | CS dis. | ranked | post-pre ACQ | pre EXT | 0.664 | 0.112 | 0.444 | 0.884 | 5.936 | 77 | 0.000 | 0.299 | 0.427 |
| Fear Ratings | CS dis. | ranked | post ACQ | pre EXT | 0.534 | 0.081 | 0.375 | 0.693 | 6.569 | 92 | 0.000 | 0.300 | 0.428 |
| Fear Ratings | CS dis. | ranked | post-pre ACQ | pre-post EXT | 0.595 | 0.117 | 0.366 | 0.824 | 5.082 | 76 | 0.000 | 0.241 | 0.317 |
| Fear Ratings | CS dis. | ranked | post ACQ | pre-post EXT | 0.461 | 0.084 | 0.296 | 0.626 | 5.461 | 91 | 0.000 | 0.231 | 0.301 |
| Fear Ratings | CS dis. | ranked | post-pre ACQ | post EXT | 0.269 | 0.167 | -0.058 | 0.596 | 1.613 | 79 | 0.111 | 0.033 | 0.034 |
| Fear Ratings | CS dis. | ranked | post ACQ | post EXT | 0.241 | 0.120 | 0.006 | 0.476 | 2.003 | 98 | 0.048 | 0.040 | 0.042 |
| Fear Ratings | CS dis. | ranked | post ACQ | 1st trial RI-Test | 0.213 | 0.086 | 0.044 | 0.382 | 2.471 | 74 | 0.016 | 0.072 | 0.078 |
| Fear Ratings | CS dis. | ranked | post-pre ACQ | 1st trial RI-Test | 0.236 | 0.116 | 0.009 | 0.463 | 2.043 | 60 | 0.045 | 0.066 | 0.071 |
| Fear Ratings | CS dis. | ranked | pre EXT | 1st trial RI-Test | 0.217 | 0.094 | 0.033 | 0.401 | 2.309 | 69 | 0.024 | 0.075 | 0.082 |
| Fear Ratings | CS dis. | ranked | pre-post EXT | 1st trial RI-Test | 0.138 | 0.097 | -0.052 | 0.328 | 1.417 | 68 | 0.161 | 0.029 | 0.030 |
| Fear Ratings | CS dis. | ranked | post EXT | 1st trial RI-Test | 0.143 | 0.081 | -0.016 | 0.302 | 1.769 | 71 | 0.081 | 0.047 | 0.050 |

| Outcome | Stim.-type | Ranking | Predictor | Criterion | *b* | $SE_b$ | Lower 95% CI | Upper 95% CI | *t* | *df* | *p* | $R^2$ | Cohen's $f^2$ |
|---|---|---|---|---|---|---|---|---|---|---|---|---|---|
| Fear Ratings | CS+ | ranked | post-pre ACQ | pre EXT | 0.516 | 0.101 | 0.318 | 0.714 | 5.107 | 91 | 0.000 | 0.215 | 0.274 |
| Fear Ratings | CS+ | ranked | post ACQ | pre EXT | 0.584 | 0.087 | 0.413 | 0.755 | 6.675 | 97 | 0.000 | 0.326 | 0.484 |
| Fear Ratings | CS+ | ranked | post-pre ACQ | pre-post EXT | 0.497 | 0.094 | 0.313 | 0.681 | 5.267 | 90 | 0.000 | 0.202 | 0.253 |
| Fear Ratings | CS+ | ranked | post ACQ | pre-post EXT | 0.442 | 0.090 | 0.266 | 0.618 | 4.930 | 96 | 0.000 | 0.191 | 0.236 |
| Fear Ratings | CS+ | ranked | post-pre ACQ | post EXT | 0.070 | 0.136 | -0.197 | 0.337 | 0.517 | 92 | 0.607 | 0.003 | 0.003 |
| Fear Ratings | CS+ | ranked | post ACQ | post EXT | 0.208 | 0.119 | -0.025 | 0.441 | 1.744 | 101 | 0.084 | 0.029 | 0.030 |
| Fear Ratings | CS+ | ranked | post ACQ | 1st trial RI-Test | 0.364 | 0.087 | 0.193 | 0.535 | 4.192 | 85 | 0.000 | 0.162 | 0.193 |
| Fear Ratings | CS+ | ranked | post-pre ACQ | 1st trial RI-Test | 0.286 | 0.096 | 0.098 | 0.474 | 2.985 | 79 | 0.004 | 0.092 | 0.102 |
| Fear Ratings | CS+ | ranked | pre EXT | 1st trial RI-Test | 0.382 | 0.081 | 0.223 | 0.541 | 4.732 | 82 | 0.000 | 0.198 | 0.247 |
| Fear Ratings | CS+ | ranked | pre-post EXT | 1st trial RI-Test | 0.228 | 0.089 | 0.054 | 0.402 | 2.568 | 81 | 0.012 | 0.066 | 0.071 |
| Fear Ratings | CS+ | ranked | post EXT | 1st trial RI-Test | 0.166 | 0.076 | 0.017 | 0.315 | 2.199 | 84 | 0.031 | 0.056 | 0.059 |
| Fear Ratings | CS- | ranked | post-pre ACQ | pre EXT | 0.430 | 0.169 | 0.099 | 0.761 | 2.552 | 86 | 0.012 | 0.090 | 0.098 |
| Fear Ratings | CS- | ranked | post ACQ | pre EXT | 0.558 | 0.093 | 0.376 | 0.740 | 5.965 | 98 | 0.000 | 0.276 | 0.381 |
| Fear Ratings | CS- | ranked | post-pre ACQ | pre-post EXT | 0.086 | 0.136 | -0.181 | 0.353 | 0.629 | 85 | 0.531 | 0.006 | 0.006 |
| Fear Ratings | CS- | ranked | post ACQ | pre-post EXT | 0.192 | 0.080 | 0.035 | 0.349 | 2.405 | 97 | 0.018 | 0.059 | 0.062 |
| Fear Ratings | CS- | ranked | post-pre ACQ | post EXT | 0.250 | 0.167 | -0.077 | 0.577 | 1.500 | 88 | 0.137 | 0.030 | 0.031 |
| Fear Ratings | CS- | ranked | post ACQ | post EXT | 0.443 | 0.098 | 0.251 | 0.635 | 4.512 | 100 | 0.000 | 0.171 | 0.206 |
| Fear Ratings | CS- | ranked | post ACQ | 1st trial RI-Test | 0.144 | 0.081 | -0.015 | 0.303 | 1.775 | 88 | 0.079 | 0.037 | 0.038 |
| Fear Ratings | CS- | ranked | post-pre ACQ | 1st trial RI-Test | 0.050 | 0.123 | -0.191 | 0.291 | 0.411 | 78 | 0.682 | 0.002 | 0.002 |
| Fear Ratings | CS- | ranked | pre EXT | 1st trial RI-Test | 0.148 | 0.075 | 0.001 | 0.295 | 1.979 | 86 | 0.051 | 0.043 | 0.045 |
| Fear Ratings | CS- | ranked | pre-post EXT | 1st trial RI-Test | 0.003 | 0.103 | -0.199 | 0.205 | 0.025 | 85 | 0.980 | 0.000 | 0.000 |
| Fear Ratings | CS- | ranked | post EXT | 1st trial RI-Test | 0.249 | 0.071 | 0.110 | 0.388 | 3.503 | 87 | 0.001 | 0.126 | 0.145 |

Note. Stim. = Stimulus, CI = Confidence Interval, CS dis. = CS discrimination, pre = prior to the experimental phase, post = subsequent to the experimental phase, ACQ = Acquisition training, EXT = Extinction training, RI = Reinstatement, RI-Test = Reinstatement-Test.
